# The identification of DNA binding regions of the σ54 factor using artificial neural network

**DOI:** 10.1101/393736

**Authors:** L. Ferreira, R. T. Raittz, J. N. Marchaukoski, V. A. Weiss, I. C. R. Santos-Wiess, P. A. B. Costa, R. Voyceik, L. U. Rigo

## Abstract

Transcription of many bacterial genes is regulated by alternative RNA polymerase sigma factors as the sigma 54 (σ54). A single essential σ promotes transcription of thousands of genes and many alternative σ factors promote transcription of multiple specialized genes required for coping with stress or development. Bacterial genomes have two families of sigma factors, sigma 70 (σ70) and sigma 54 (σ54). σ54 uses a more complex mechanism with specialized enhancers-binding proteins and DNA melting and is well known for its role in regulation of nitrogen metabolism in proteobacteria. The identification of these regulatory elements is the main step to understand the metabolic networks. In this study, we propose a supervised pattern recognition model with neural network to identify Transcription Factor Binding Sites (TFBSs) for σ54. This approach is capable of detecting σ54 TFBSs with sensitivity higher than 98% in recent published data. False positives are reduced with the addition of ANN and feature extraction, which increase the specificity of the program. We also propose a free, fast and friendly tool for σ54 recognition and a σ54 related genes database, available for consult. S54Finder can analyze from short DNA sequences to complete genomes and is available online. The software was used to determine σ54 TFBSs on the complete bacterial genomes database from NCBI and the result is available for comparison. S54Finder does the identification of σ54 regulated genes for a large set of genomes allowing evolutionary and conservation studies of the regulation system between the organisms.

## 1. Background

Prokaryotic transcription process starts from the recognition of binding site on DNA sequence by RNA polymerase. The DNA dependent RNA Polymerase (RNAP) is the main enzyme in the transcription and gene expression [1]. It is composed of two subunits, the core enzyme α2ββ with the catalytic site of polymerization and the sigma factors, responsible for promoter recognition [1]. When the enzyme is linked with the sigma factor protein, the complex is called of RNAP holoenzyme (α2ββ’σ) and is ready for transcription [2].

Bacterial genomes have two families of sigma factors, sigma 70 (σ70) and sigma 54 (σ54). σ70 is the best studied and is responsible for transcription of the most bacterial genes in the exponential growth [3]. σ54 uses a more complex mechanism with specialized enhancers-binding proteins and DNA melting and is well known for its role in regulation of nitrogen metabolism in proteobacteria [4].

Each sigma factor recognizes different binding sites in the DNA sequence and is responsible for changing the transcription pattern mediated thorough redirection of transcription initiation [5]. The identification of these regulatory elements is the main step to understand the metabolic networks [6].

A single essential σ promotes transcription of thousands of genes and many alternative σ factors promote transcription of multiple specialized genes required for coping with stress or development [7].

The number of sigma factors varies from one in *Mycoplasma genitalia* [8], seven in *Escherichia coli* [9] to more than 60 in *Streptomyces coelicolor* [10].

It was already shown that σ70 recognize a consensus sequence of hexamers placed between -10/-35 nucleotides upstream the start site and σ54 recognize high GC sequences located -24/-12 nucleotides upstream from the start site [11].

The σ54 factor has an important role in agriculture, due to the control of gene transcription, regulating the nitrogen fixation process that provides ammonia for the plant. This process dispenses chemical soil fertilization, reduces costs and protects the environment [12].

The classical approach for promoter prediction involves the development of algorithms that uses position weight matrices (PWMs) [13-17]. More recently, several methods propose the automatic identification of Transcription Factor Binding Sites (TFBSs) like Gibbs sampling algorithm [18, 19], genetic algorithms [20], and statistical over-representation [21, 22]. These methods works with single and double motifs identification but with a high false positive rate [23], and don’t allow complete genome sequences.

In this study, we propose the use of a supervised pattern recognition model with neural network to identify TFBSs for σ54 for whole genome sequence. The advantages of using neural network in pattern recognition include its generalization ability considering both the nonlinear and the diffuse nature of the datasets [24] and also can achieve high performance when processing extended genome sequences [25].

We also propose a free, fast and friendly tool for σ54 recognition and a σ54 database available for consult.

## 2. Methods

### 2.1. Candidates sequences for σ54 TFBS and features extraction

The set of intergenic regions to search for σ54 TFBSs candidates was defined as a range of 200 bases from the transcriptional start site of each coding sequence. Candidate sequences were set on 16 bases. The selection was done considering the presence of conserved bases (GG and GC) located in -14/-15 and -2/-3 positions [11]. 18 features were extracted from the candidate sequences. The first sixteen were the nucleotides encoded to numbers (A: 0.0; C: 1.0; G: 2.0; T: 3.0); the 17^th^ feature was the result of the alignment of the candidate sequence with the consensus sequence, extracted from experimentally confirmed sequences available in literature [11, 26, 27]; and the 18^th^ feature was the result of the alignment of the sequence candidate with a consensus of the less frequent bases (present in the true sequences-called as anti-consensus).

### 2.2. Training data set

Networks were trained using groups of positive sequences collected in the literature and sequences artificially generated, considered as negatives. Those sequences were generated using the MatLab algorithm of string generation. The artificially generated strings were divided into two groups, the first group have only nucleotide base pairs and the same string length of positive sequences. In other side the second group uses also the same length, but it also includes the conserved base pairs shown in the positive sequences.

Positive patterns were defined by sequences experimentally retrieved [26, 27] and a database with 5662 sequences from more than 60 genomes, available in www.sigma54.ca web site [28]. The artificial dataset comprised two groups of random sequences. The groups had the same number of bases observed in the positive pattern and differ by the presence and absence of the highly conserved bases (GG and GC).

### 2.3. ANN test

Tests with MLP (*Multilayer perceptron*), FAN (*Free Associative Neurons*), RBF (*Radial Basis Function*) and SVM (*Support Vector Machine*) neural networks were performed. RBF and SVM did not show better results than MLP and FAN neural networks. The best architecture used to classify the sequences using MLP neural network had 7 neurons in the first hidden layer and three neurons in the second one. For FAN neural network we used the default setting with fuzzy set support of 100, diffuse neighboring of 6, shuffling the training set every time. The accuracy for each neural network tested on the same database was 95.19% for MLP and 97.95% for the chosen one, the FAN network (**Table 1**).

**Table 1.** Performance comparison for neural networks.

|  | MLP | FAN |
| --- | --- | --- |
| True positives, % | 92.15 | **95.51** |
| False positives, % | 3.29 | 2.09 |
| True negatives, % | 96.71 | 97.91 |
| False negatives, % | 7.85 | 4.49 |
| Accuracy, % | 95.19 | **97.95** |

The ANN test was performed using 3 datasets from published studies comprising 281 proposed σ54 TFBS sequences that were not used for training (**Table 2**). For more recent data, the sensitivity test showed more than 98% of right predictions for σ54 binding sites sequences. The decrease of sensitivity observed on Barrios database was due to the presence of candidate sequences in the σ54 TFBS database.

**Table 2.** ANN sensitivity in comparison with published data.

| Dataset | ANN Positive | Total | Sensitivity (%) |
| --- | --- | --- | --- |
| Studholme, 2000 [29] | 34 | 34 | 100 |
| Leang, 2009 [30] | 90 | 91 | 98.9 |
| Barrios, 1999 [11] | 127 | 156 | 81.4 |

The sequences that did not had the conserved bases GG and GC and without known amino acids, were excluded from the test.

### 2.4. S54Finder tool

S54Finder is a fast tool developed to work with complete genome sequences. It works with annotated sequences stored in a gbk file format parsing the information to perform the analysis. If the query is a flat fasta file the software uses an in-house ORFinder to identify and annotate the coding regions and extract the intergenic sequences. The output of the S54Finder is 2 files, a new gbk file with the predicted σ54 sequences marked and a text file with the predicted sequences and related ORFs positions to download.

## 3. Results and discussion

There is no TFBSs consensus for σ54, with only few sequences confirmed in laboratory. Several studies present candidate sequences as consensus but the size usually is not the same, since it differs between species. What is observed is that among the vast majority of the sequences there are at least 4 (GG and GC) conserved bases. This scenario turns difficult the pattern recognition since it increases the false positive rate. To overcome this problem we added the anti-consensus feature, which allow us to decrease candidate number usually observed in other tools. To test S54Finder false positive rate we used 20^12^ sequences generated by combination of bases preserving the high conserved ones (GG and GC). The results showed that S54Finder identified, from the total of possibilities, 1.89% of σ54 candidate sequences. This result confirms a high specificity of the S54Finder software.

We tried to compare S54Finder with other available tools such as GenomeMatScan, but the test was not possible since GenomeMatScan does not perform whole genome predictions. The main features between the two software applications are contrasted in **Table 3**.

**Table 3.** Comparison between S54Finder and GenomeMatScan software.

|  | S54Finder | GenomeMatScan |
| --- | --- | --- |
| **Web version** | Yes | Yes |
| **Local processing** | Yes | No |
| **Method** | FAN | HMM |
| **Data entry** | Genbank file (.gbk) | Input data copy-pasted into the browser |
| **Results** | Output in Genbank formats (.gbk), MatLab (.mat) and text | Shown in browser |
| **Processing large regions of DNA** | Yes (complete genomes) | No |

HMM: Hidden Markov Models.

The distinguish feature of S54Finder is the ability to perform fast whole genome predictions. This allowed us to search for TFBSs in complete bacterial genomes across the entire NCBI database. From the complete genomes, s54finder obtained 79626 CDS sequences which could be ranked by the level of TFBSs occurrence next to homologous genes of different species.

An in house tool was employed to cluster the CDS. A set of 9417 homologous CDS groups, within 2 or more homologous occurrences were determined. The minimal relative score considered to group a pair of genes was 0.5. To determine the annotation to be considered for each gene cluster we implemented a kmeans to support the choice in the set of original annotations. The generated data – TFBSs, CDSs, annotation, Homologous clusters – were gathered to perform a FASTA/BLAST formatted data base available and a web site (http://200.236.3.16/s54.php), that is available to validate the S54Finder predictions. **Table 4** presents a list with the 29 groups that occurred 100 or more times. The glutamine synthetase protein and the nitrogen regulatory protein P II appears in the top of the list with 379 and 329 occurrences, respectively. Due to the importance of these proteins in the biologic nitrogen fixation their presence in the top predicted list was expected. We proposed a score to determine the correlation of a word present in an NCBI annotation found by s54finde with conserved clusters of genes related to TFBS. A binary matrix where columns represent the occurrence of each word and the lines the homologous groups, was joined to a column containing the number of occurrences of the respective homologous group, T. The ws54score of a word is then the Pearson correlation between the column that represent the word and homologous amount column, T. Our purpose was to identify possibly related words to σ54 regulated genes. Highest ws54scores were associated to gene names like glnL (0.3711), glnB (0.2959), glnE (0.2959) and PII (0.2959), whose association with the σ54 promoter is well known. However, words like MtrA and Thi1, also appeared in the top 10 list though not common in the literature; unknown conserved domains also appeared - DUF2233 (4) and DUF2971 (11) - in the top 15 list and we suppose they are likely related to the σ54. The list with annotation words and s54scores is in the **Additional File**.

## 4. Conclusion

The S54Finder presents good results regarding research in bacterial genomes. The main advantage is the fact that it is faster than other available tools. The pre-selection of candidates without the use of neural networks has shown that the pattern of conservation, based on published sequences and biologically confirmed, is capable of locating binding sites in bacterial genomes, but generates many false positives.

False positives are reduced with the addition of ANN and feature extraction, which increase the specificity of the program. S54Finder allows the user the identification of σ54-regulated genes for a large set of genomes allowing evolutionary and conservation studies of the regulation system between the organisms. The tool and the σ54 database are freely available in web.

## 5. Authors’ contributions

Following are the contributions of the authors to the work described in this paper. LMF: acquisition of data; analysis and interpretation of data; RTR: made substantial contributions to conception and design of the study; analysis and interpretation of data; JNM: have been involved in drafting the manuscript or revising it critically for important intellectual content; have given final approval of the version to be published; VAW: have been involved in drafting the manuscript or revising it critically for important intellectual content; have given final approval of the version to be published; ICRSW: have been involved in drafting the manuscript or revising it critically for important intellectual content; have given final approval of the version to be published; PAB: analysis and interpretation of data; RV: participated in the website development and revising the manuscript; LUR: made substantial contributions to conception and design of the study. All authors read and approved the final manuscript.

## 6. Acknowledgements

CNPq (Conselho Nacional de Desenvolvimento Científico e Tecnológico) and CAPES (Coordenação de Aperfeiçoamento de Pessoal de Nível Superior) supported this work.

---

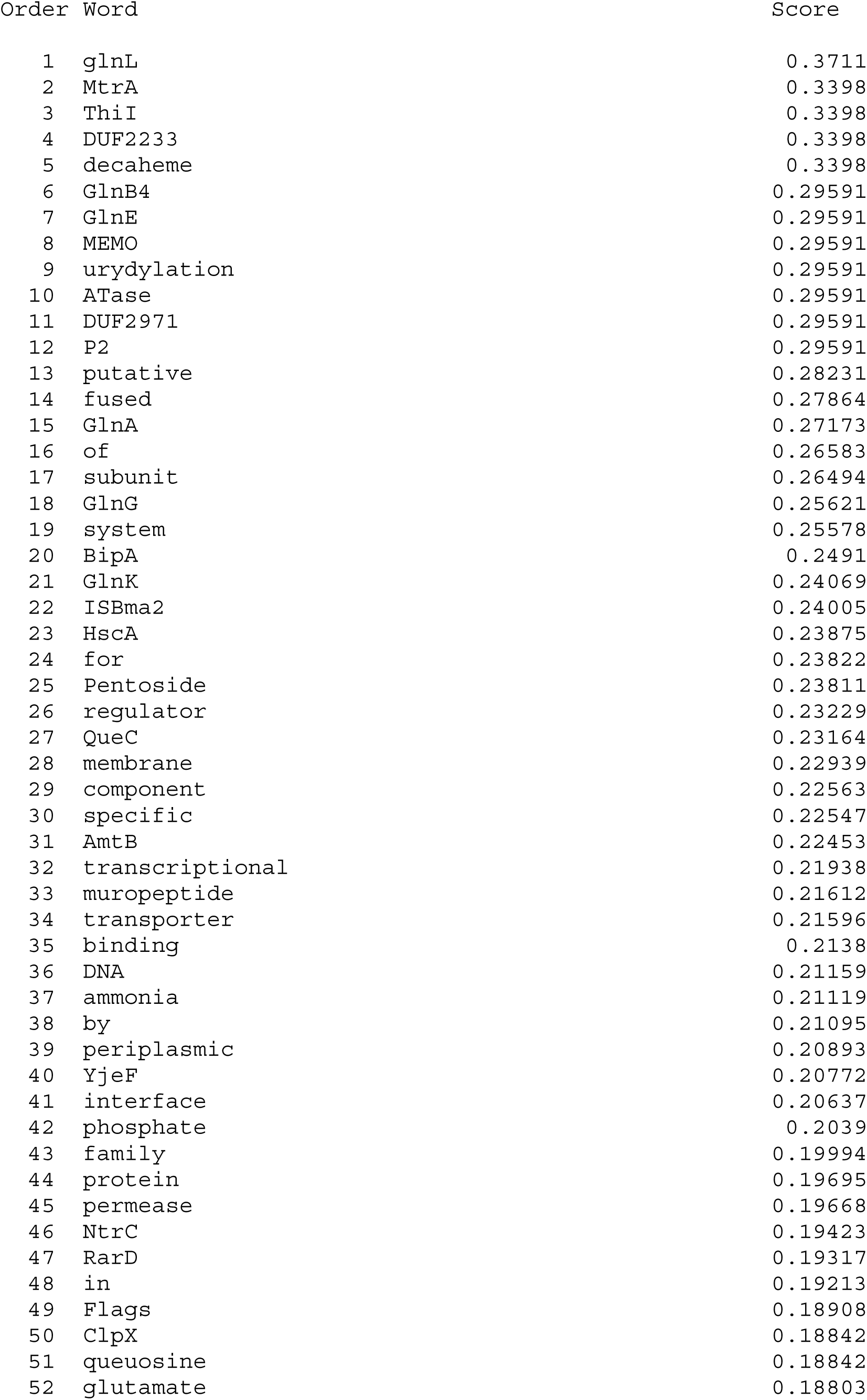

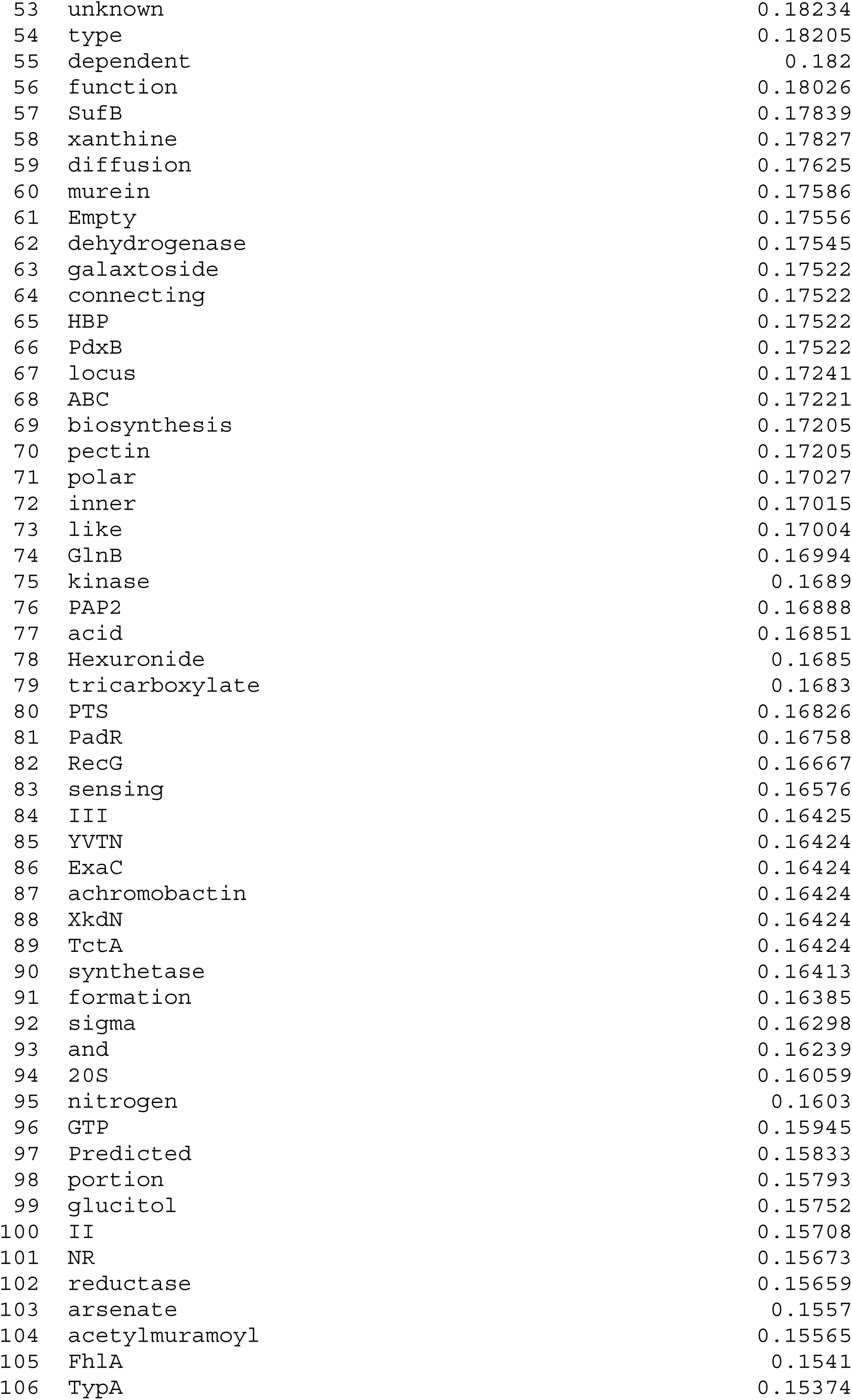

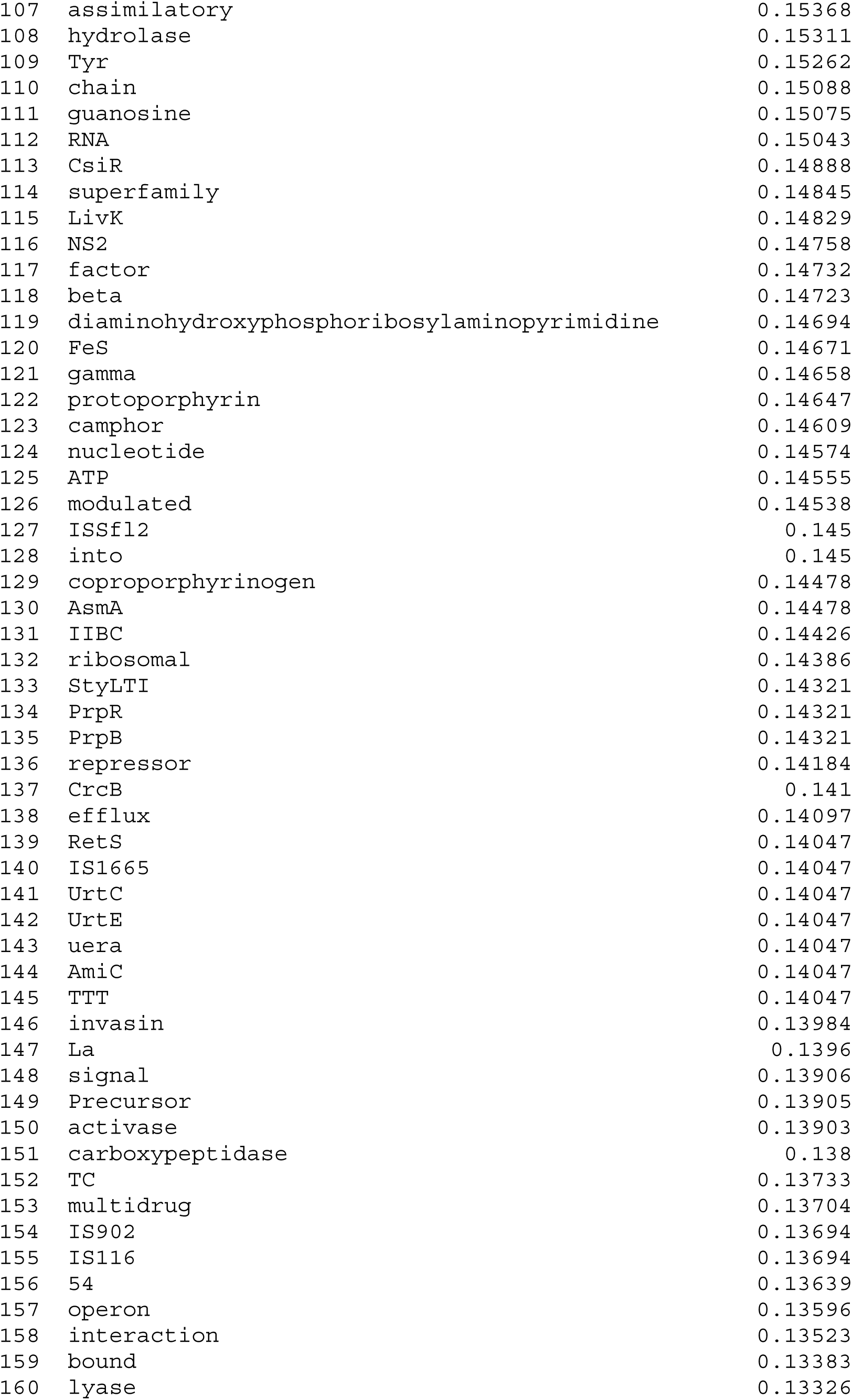

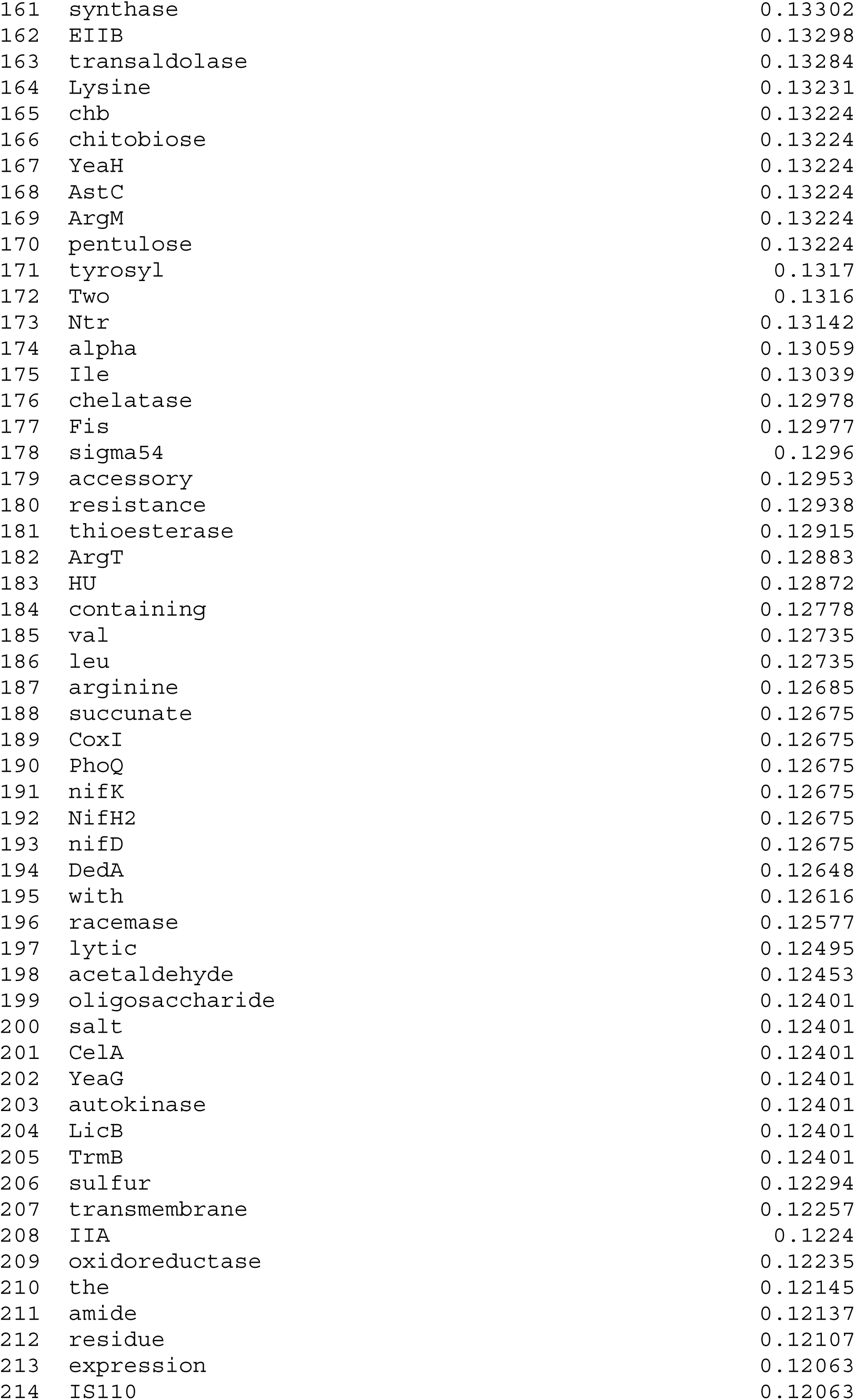

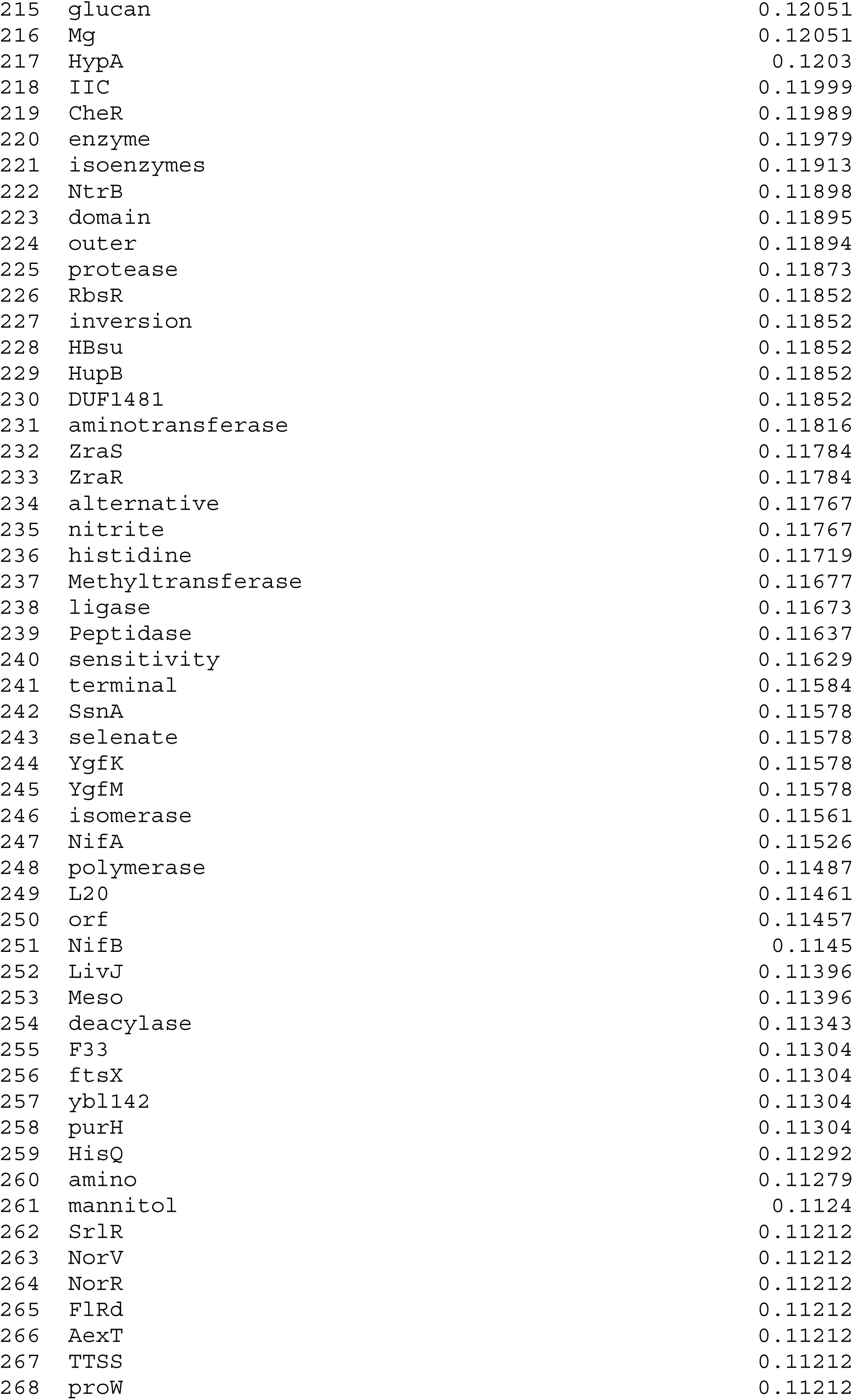

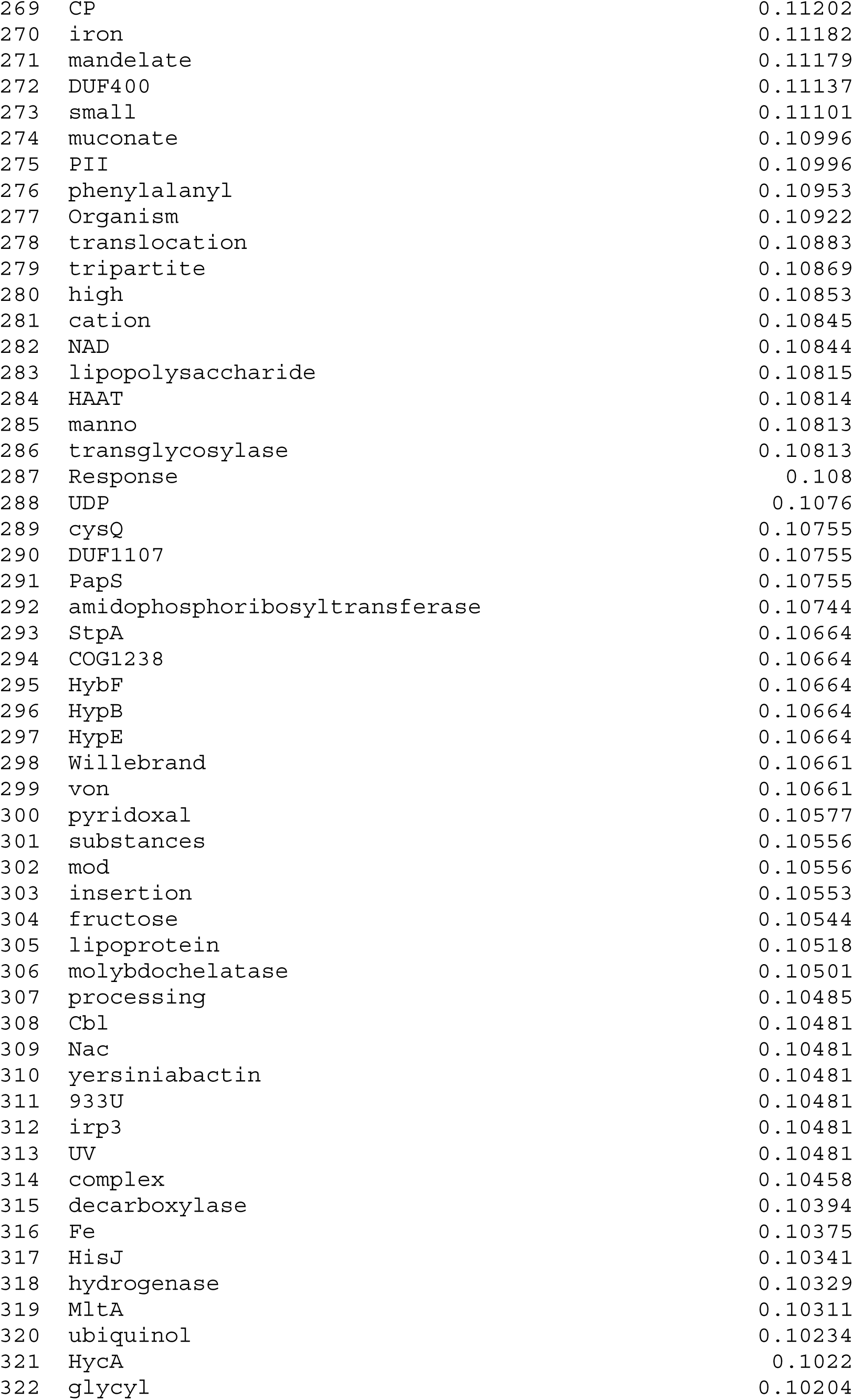

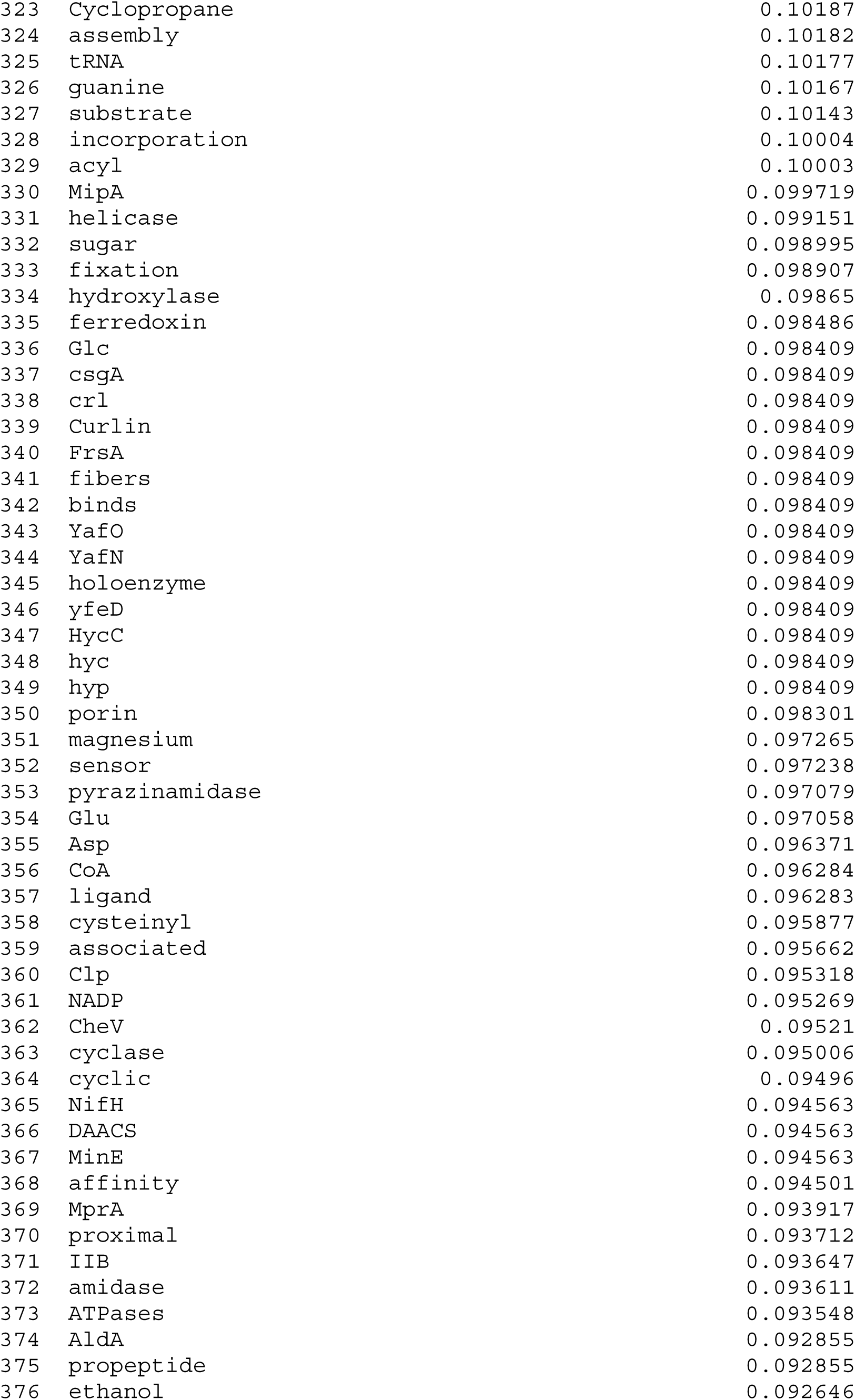

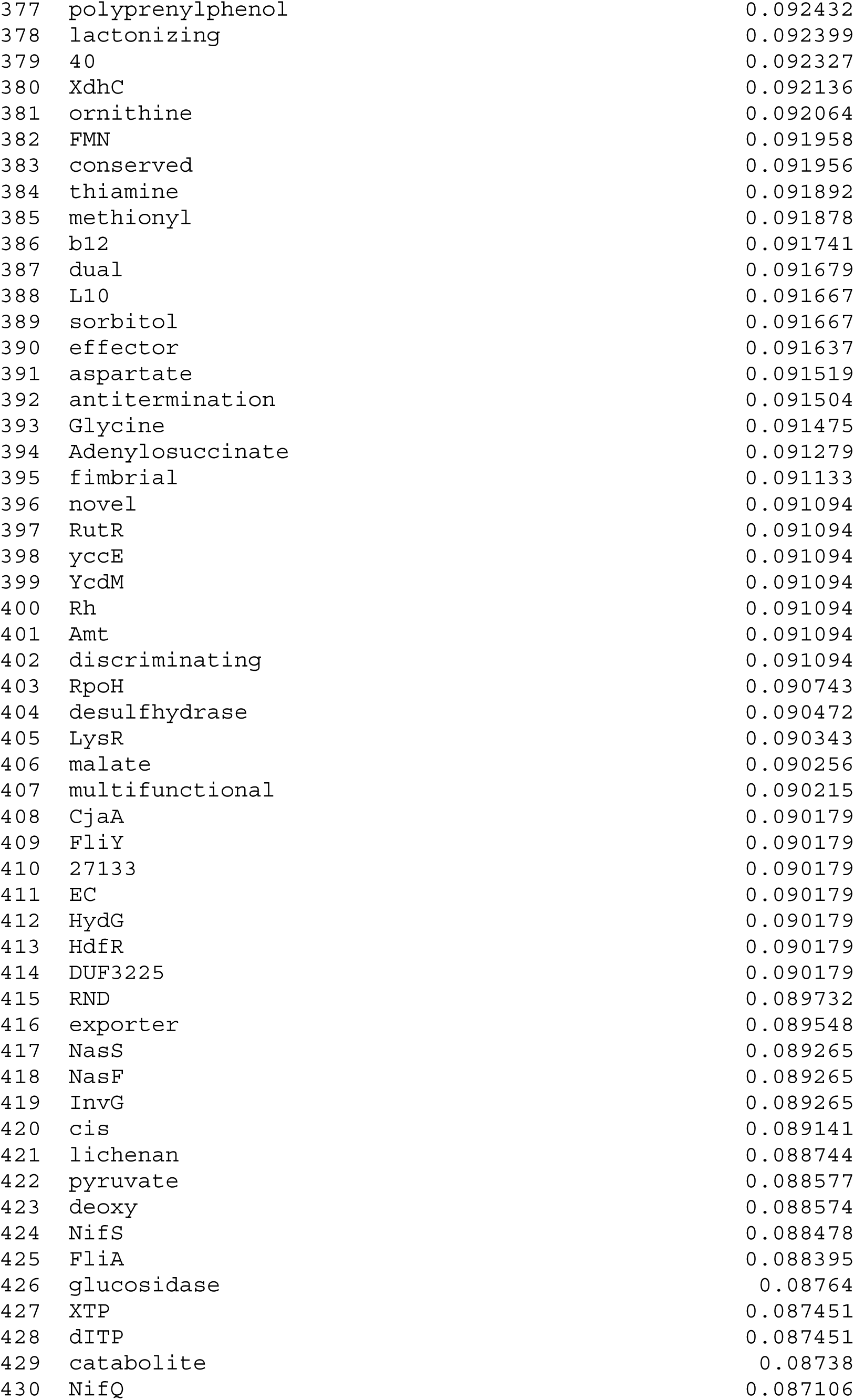

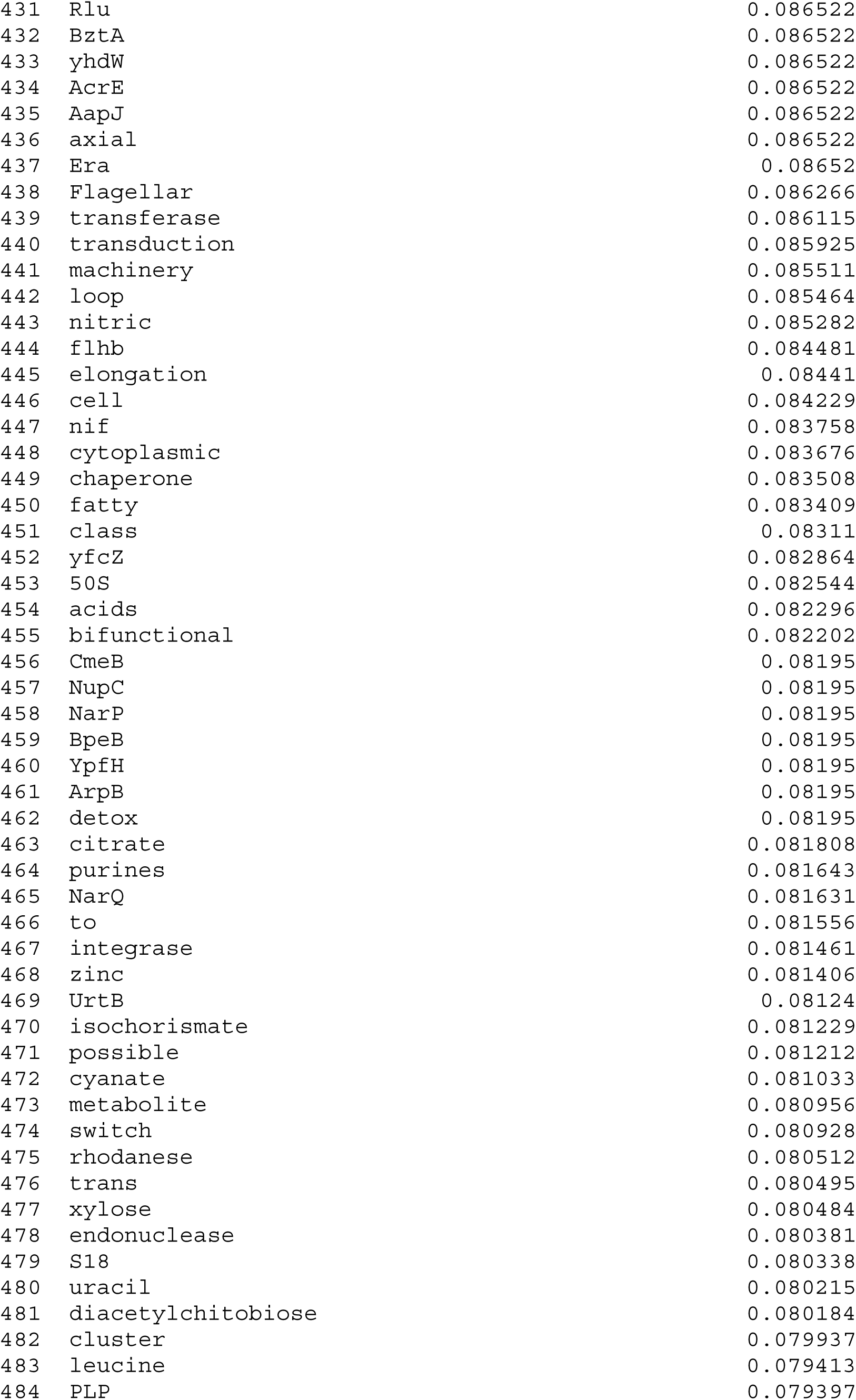

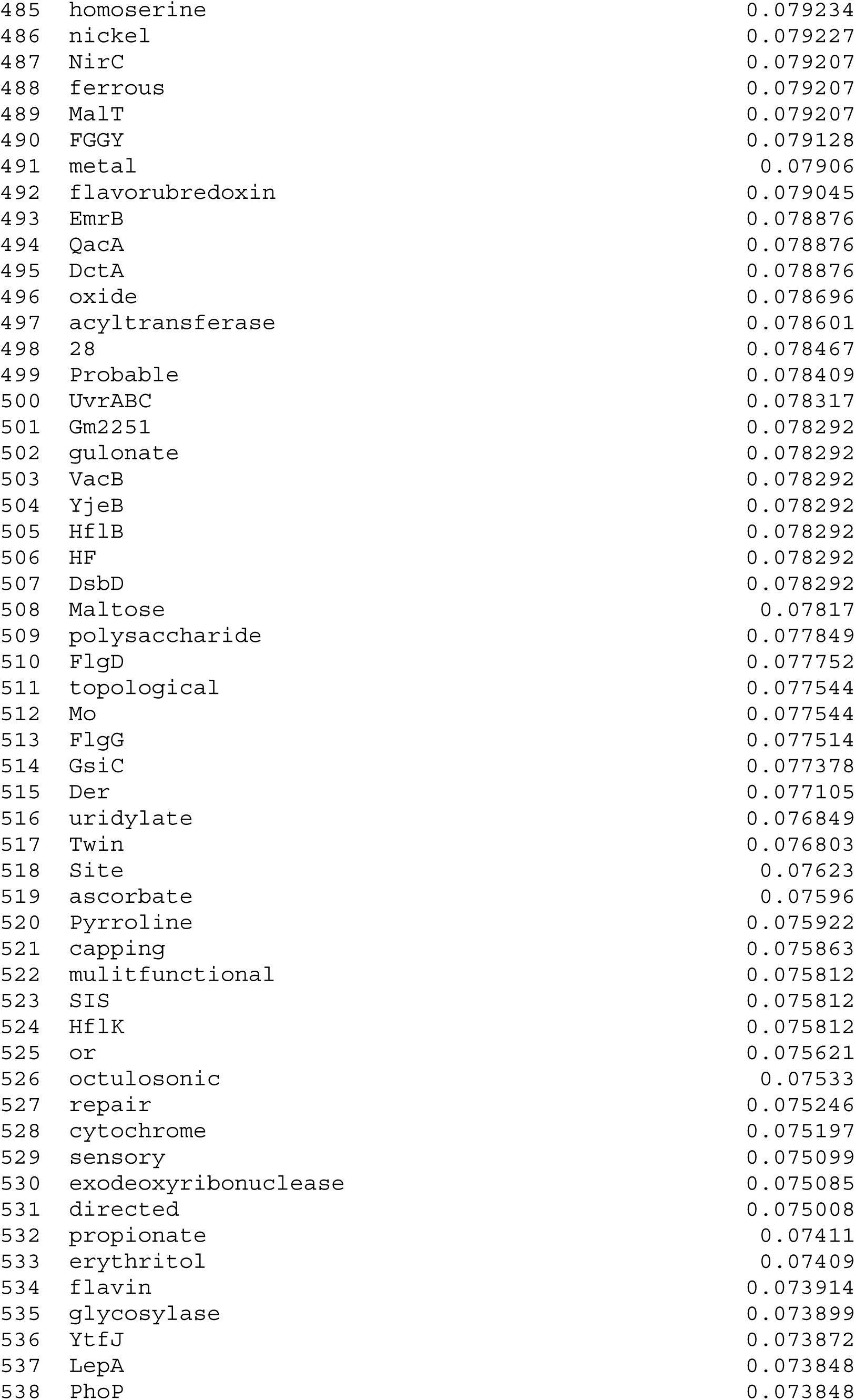

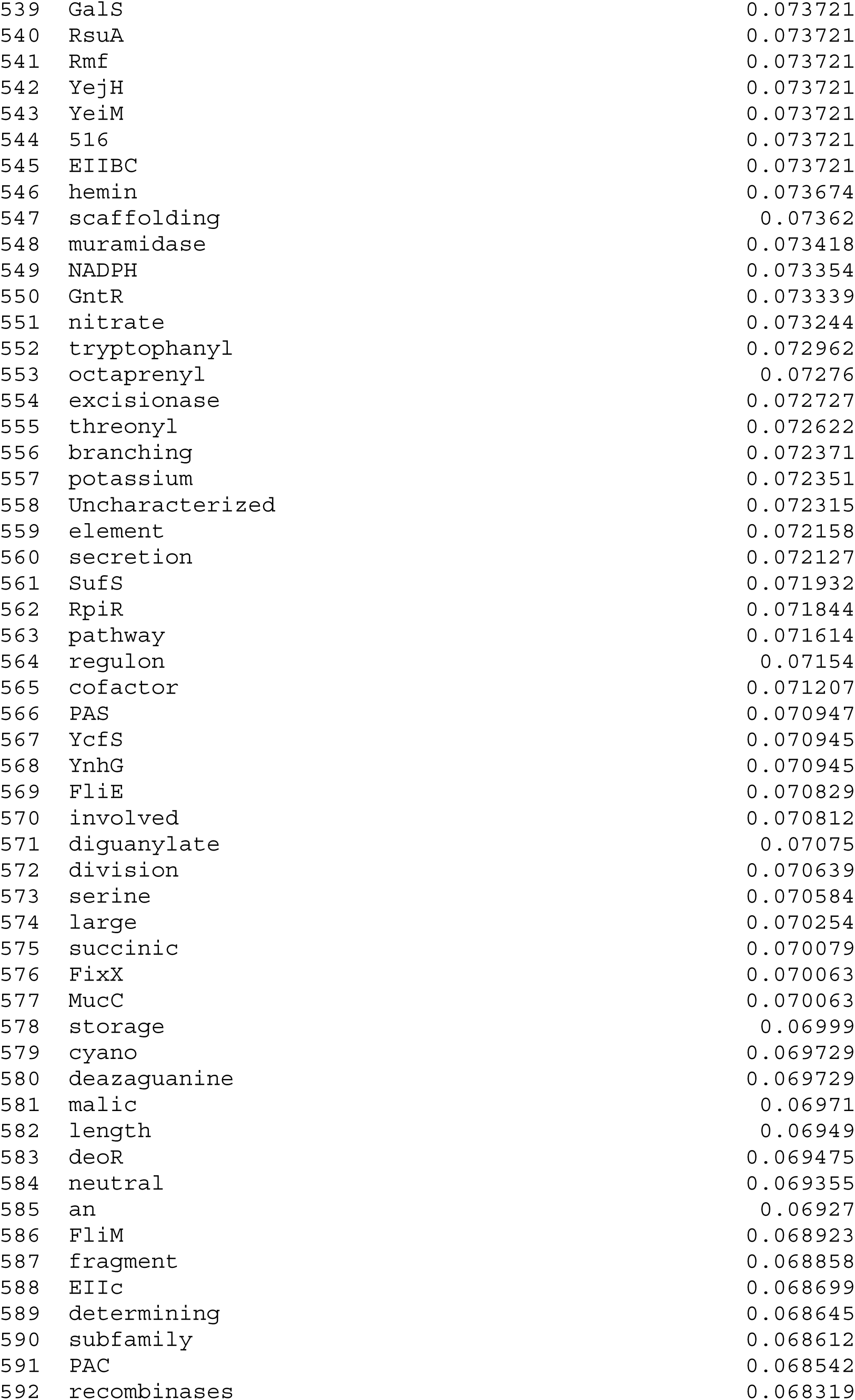

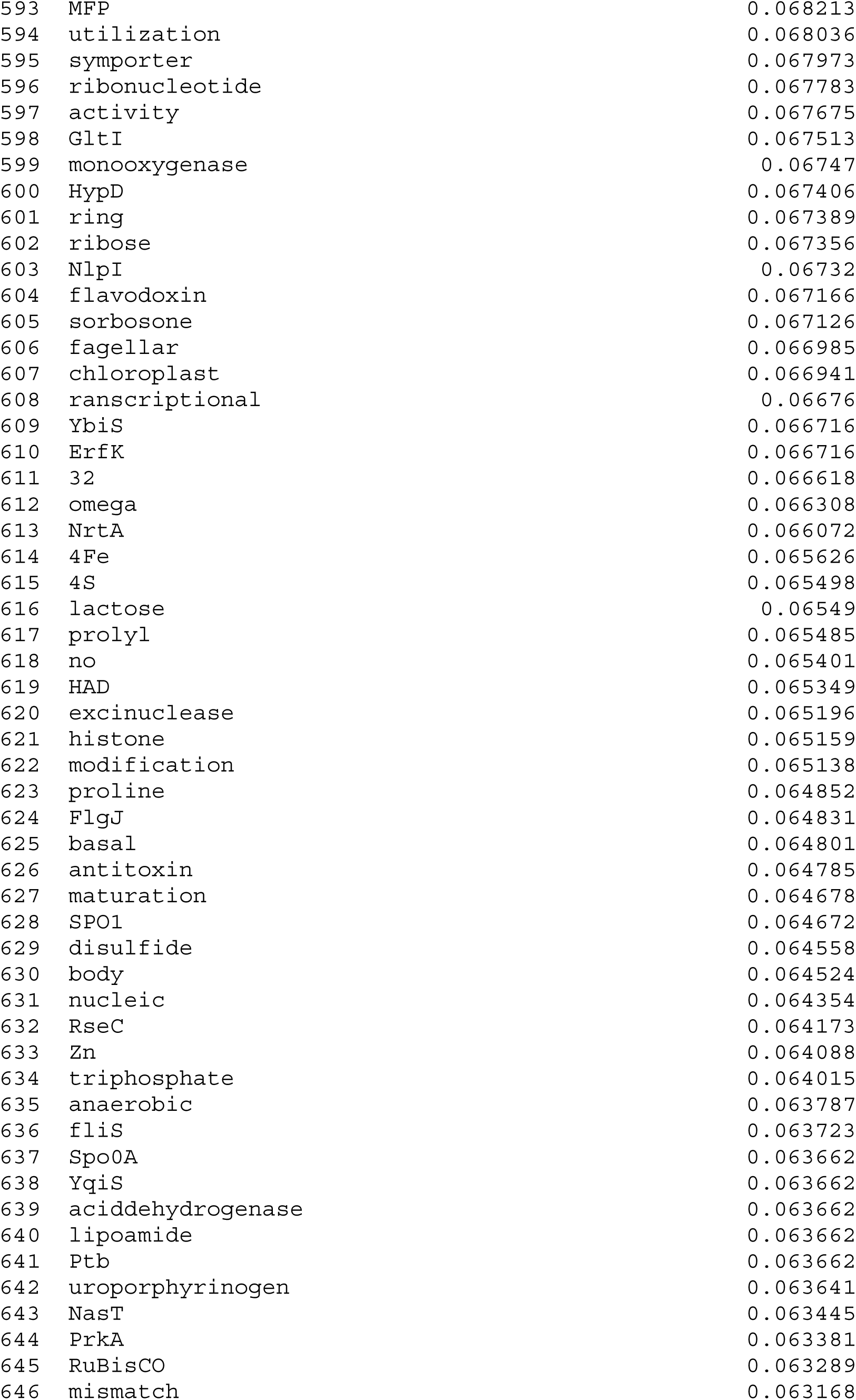

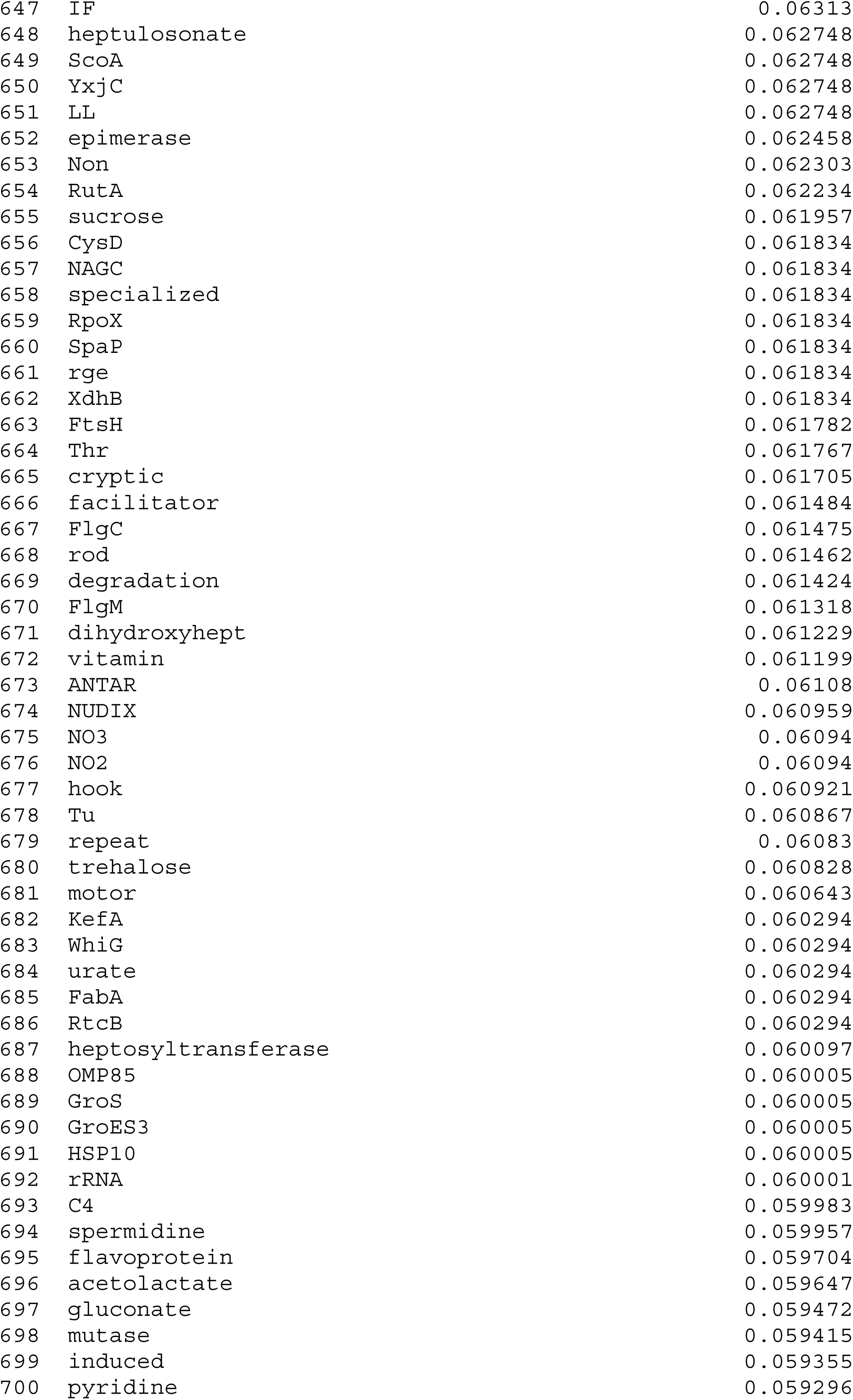

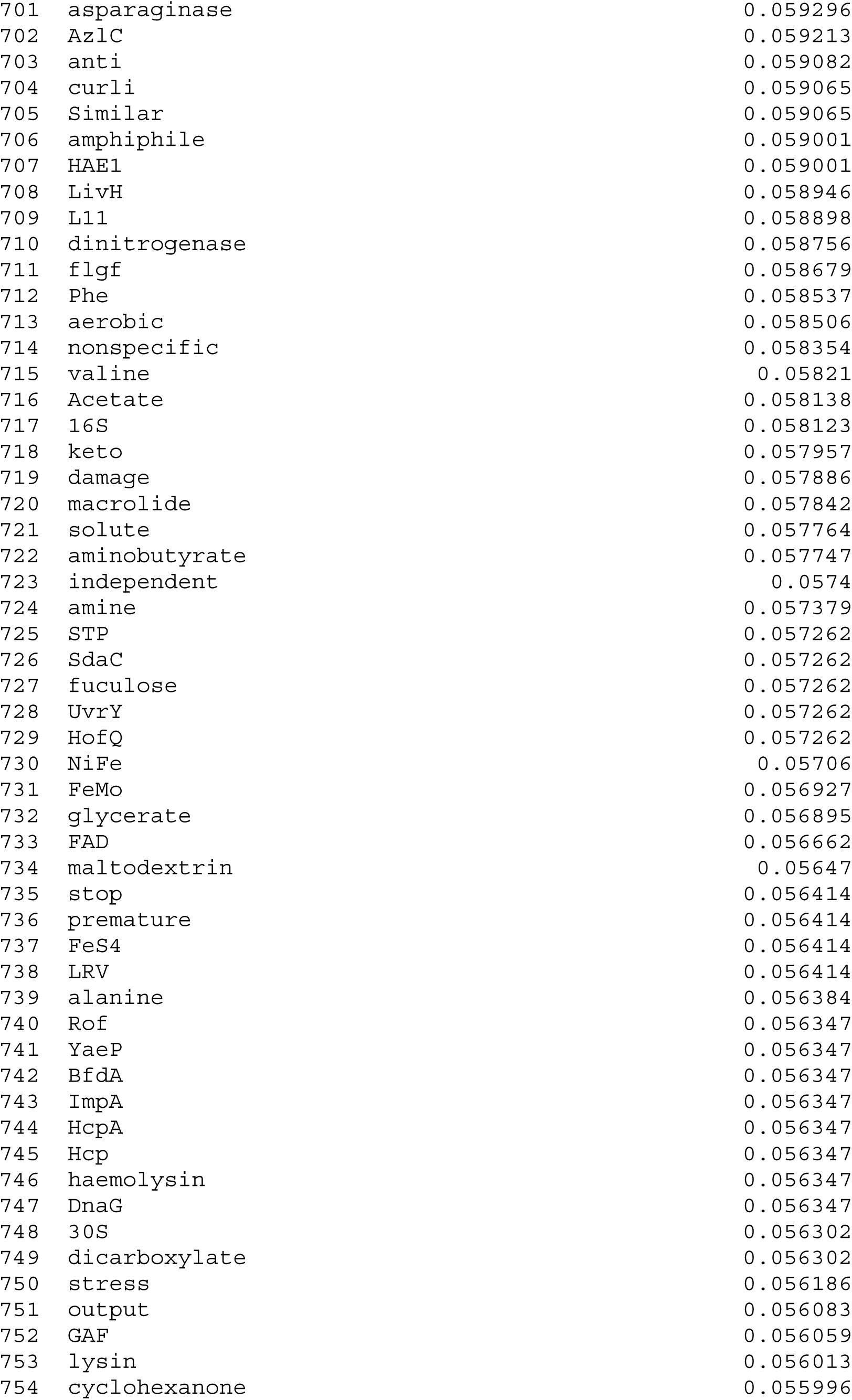

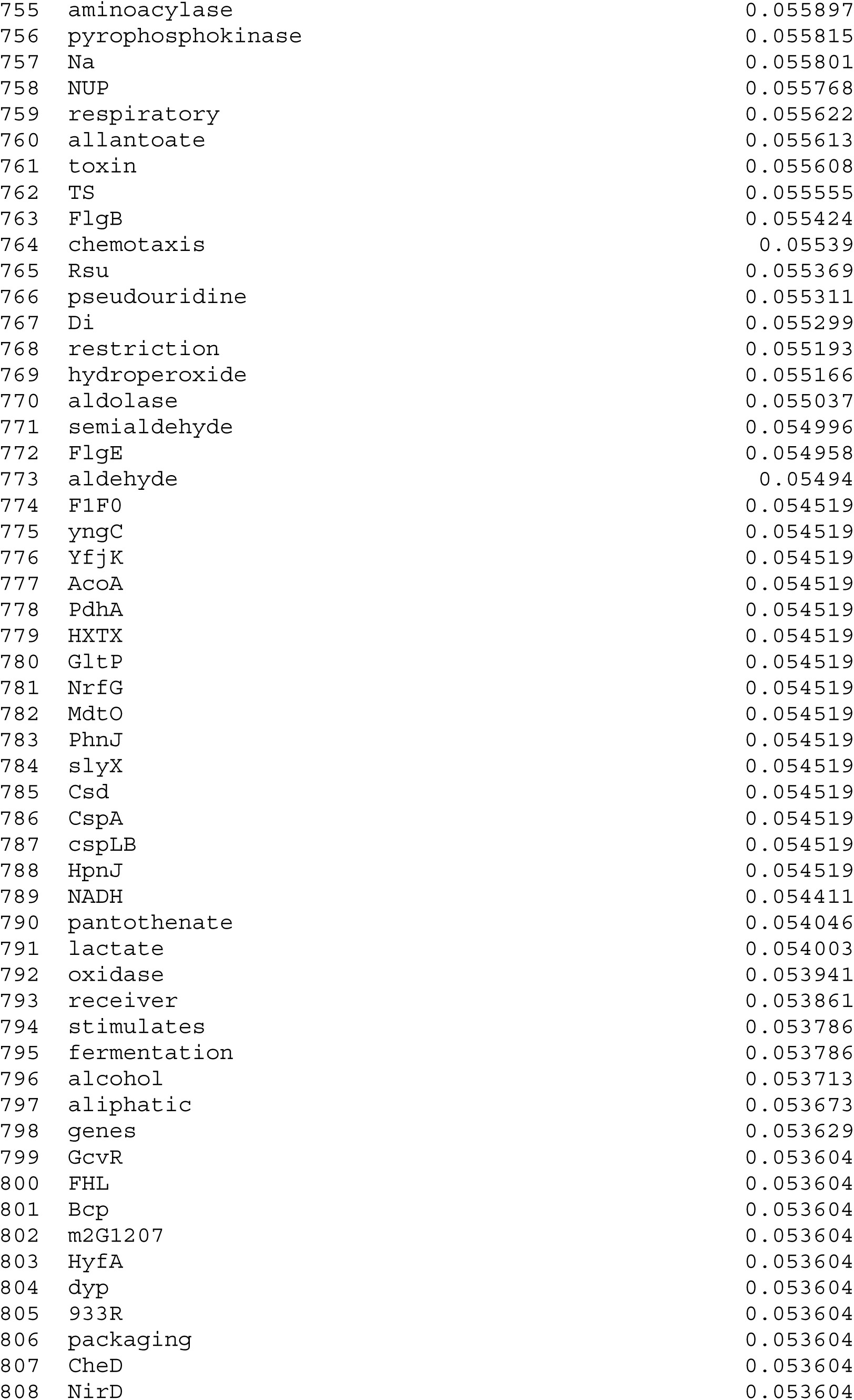

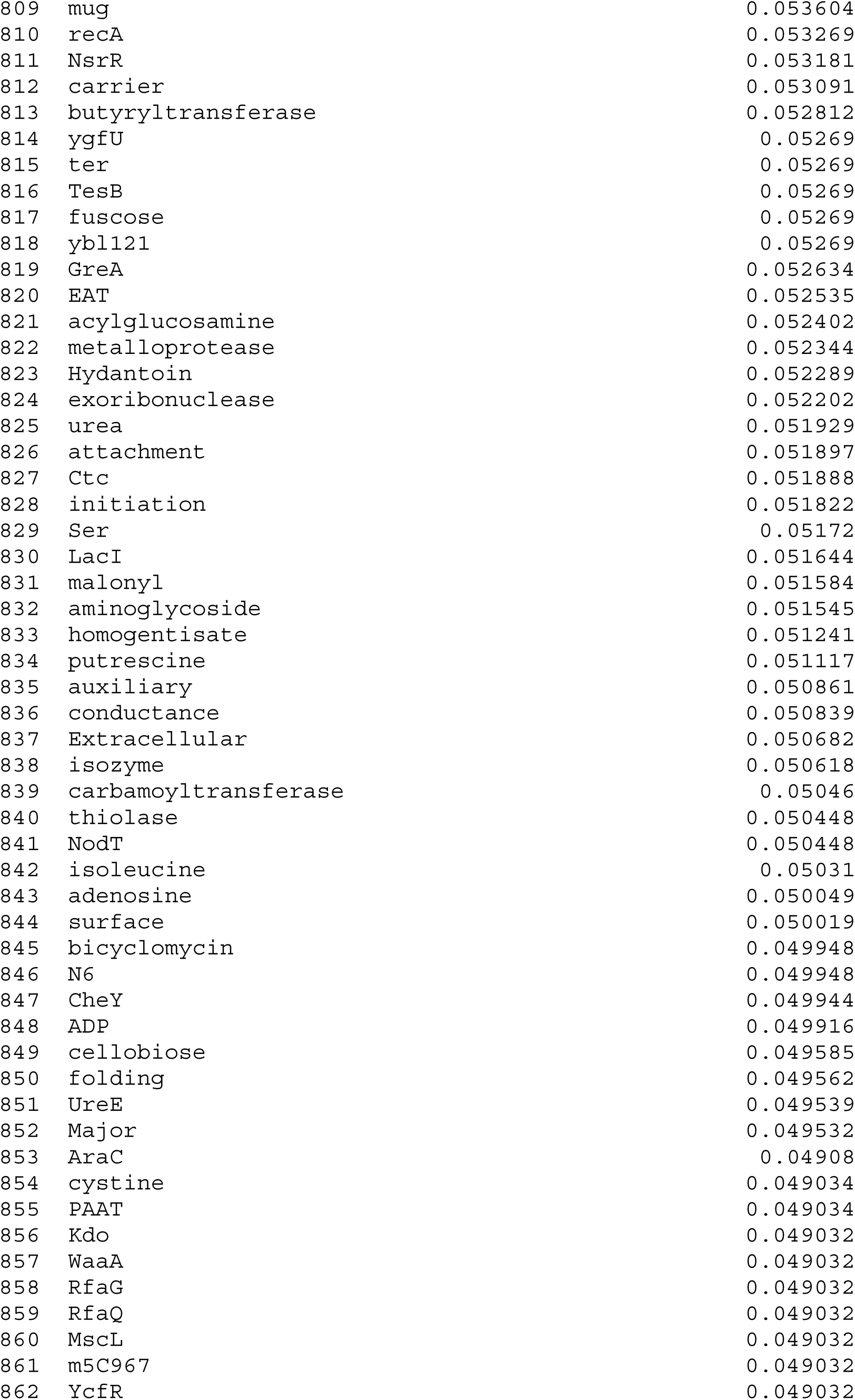

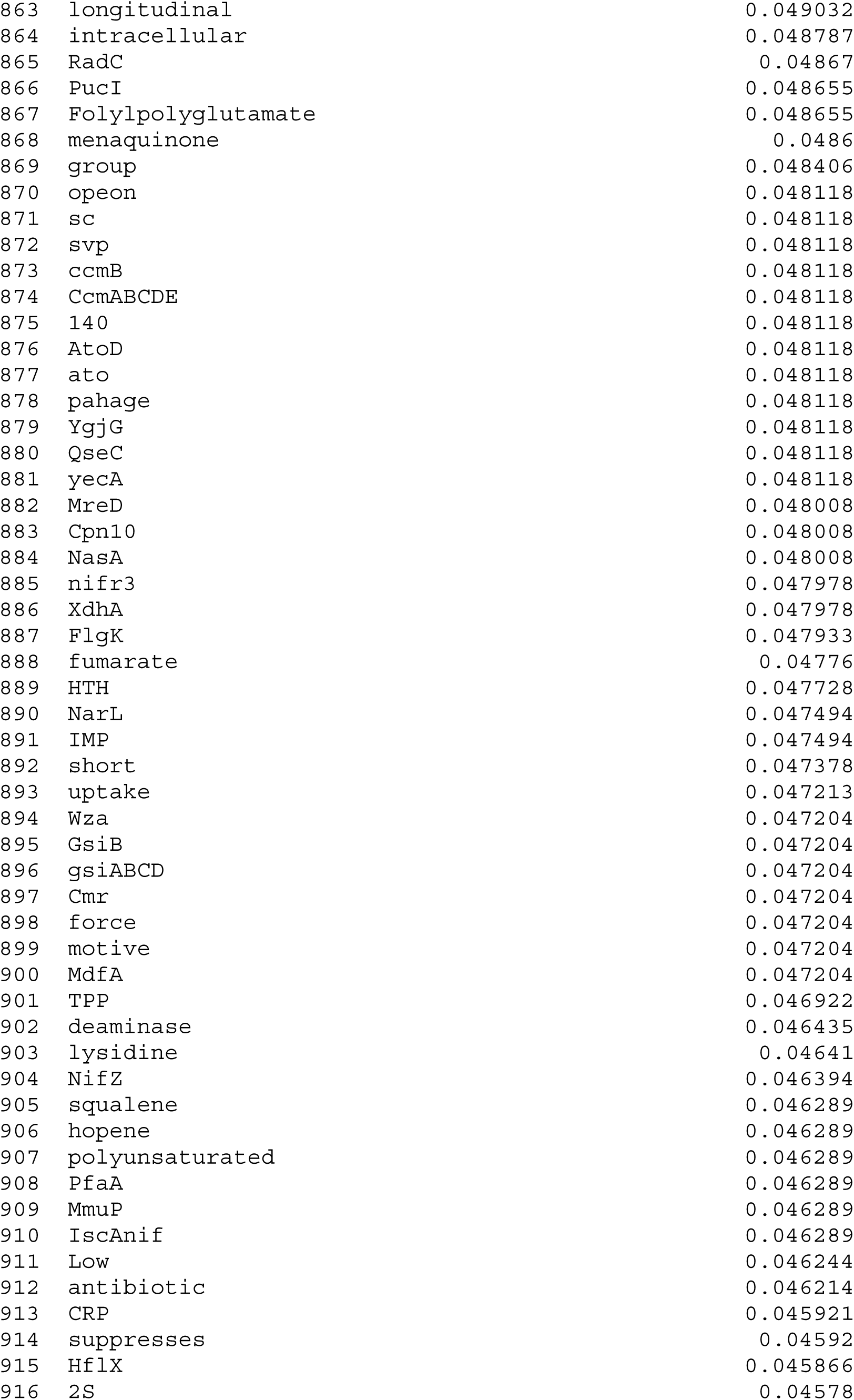

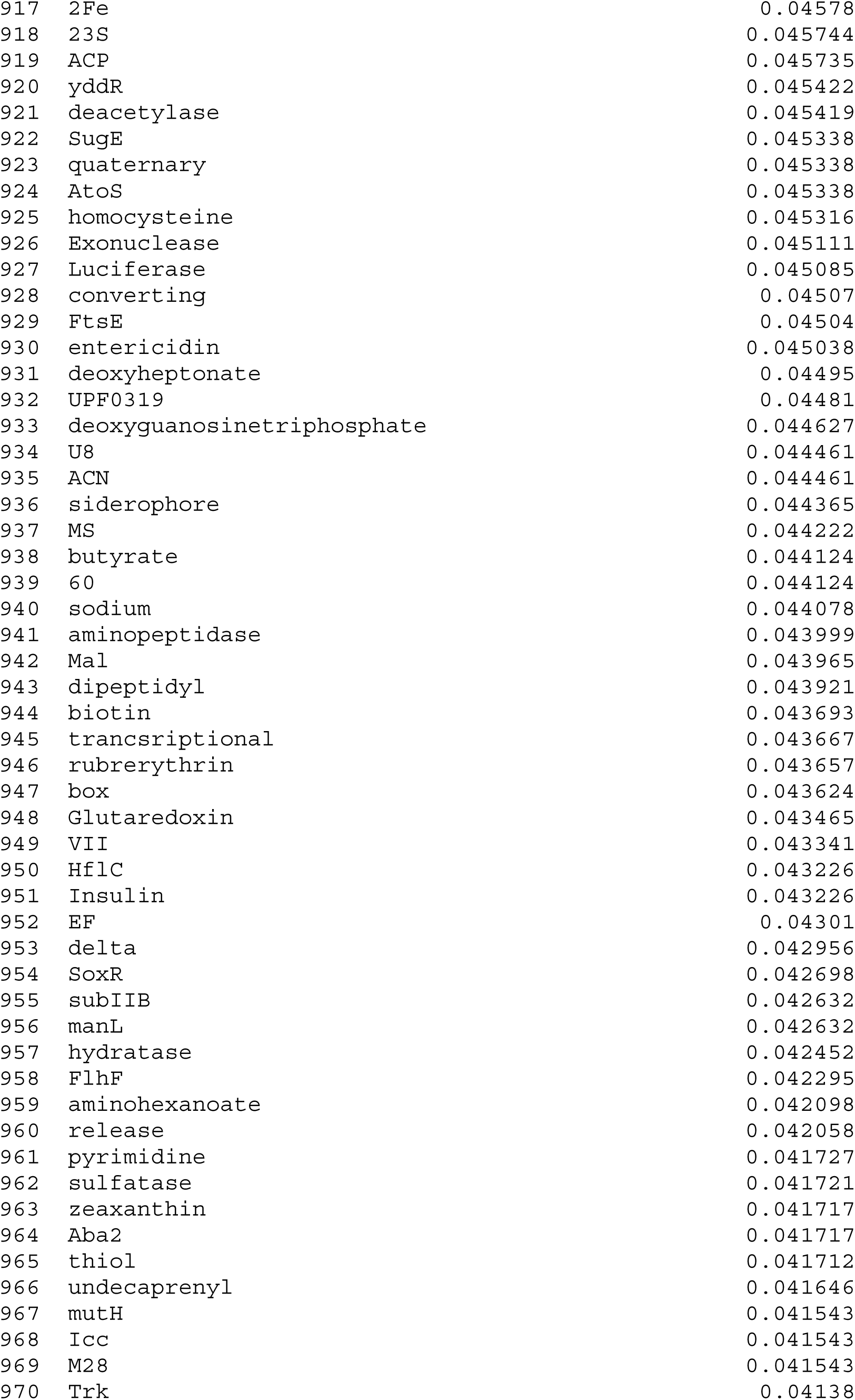

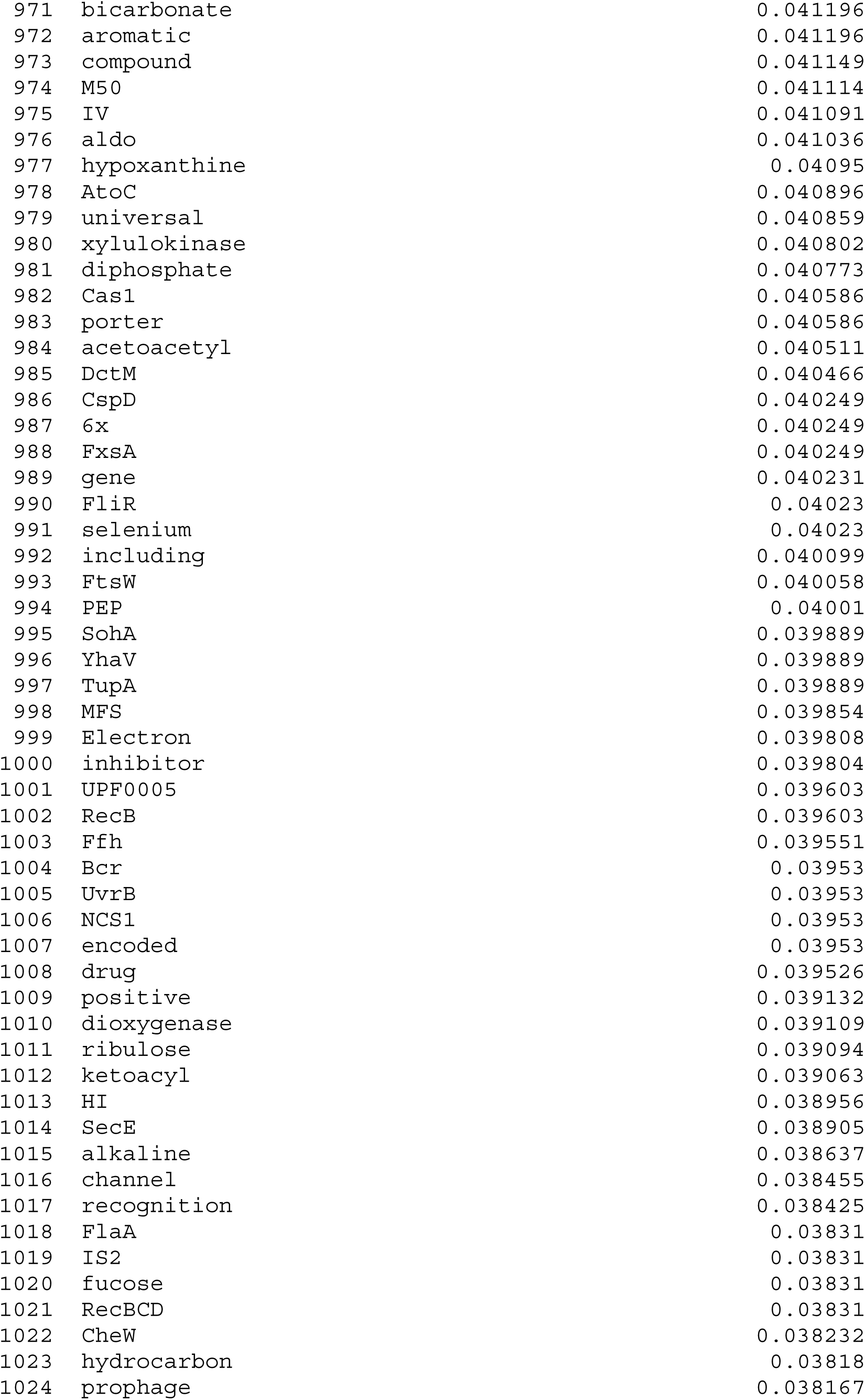

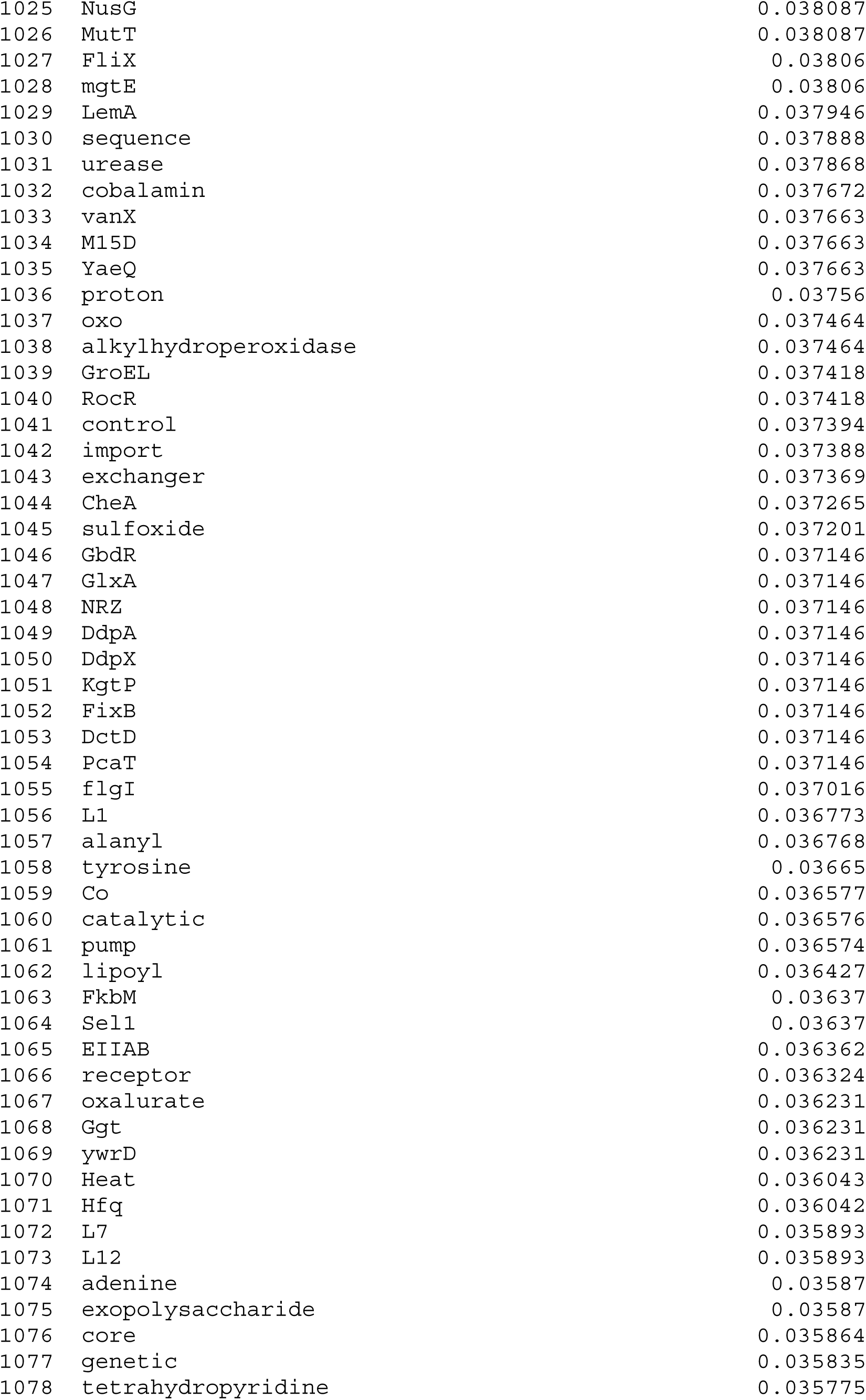

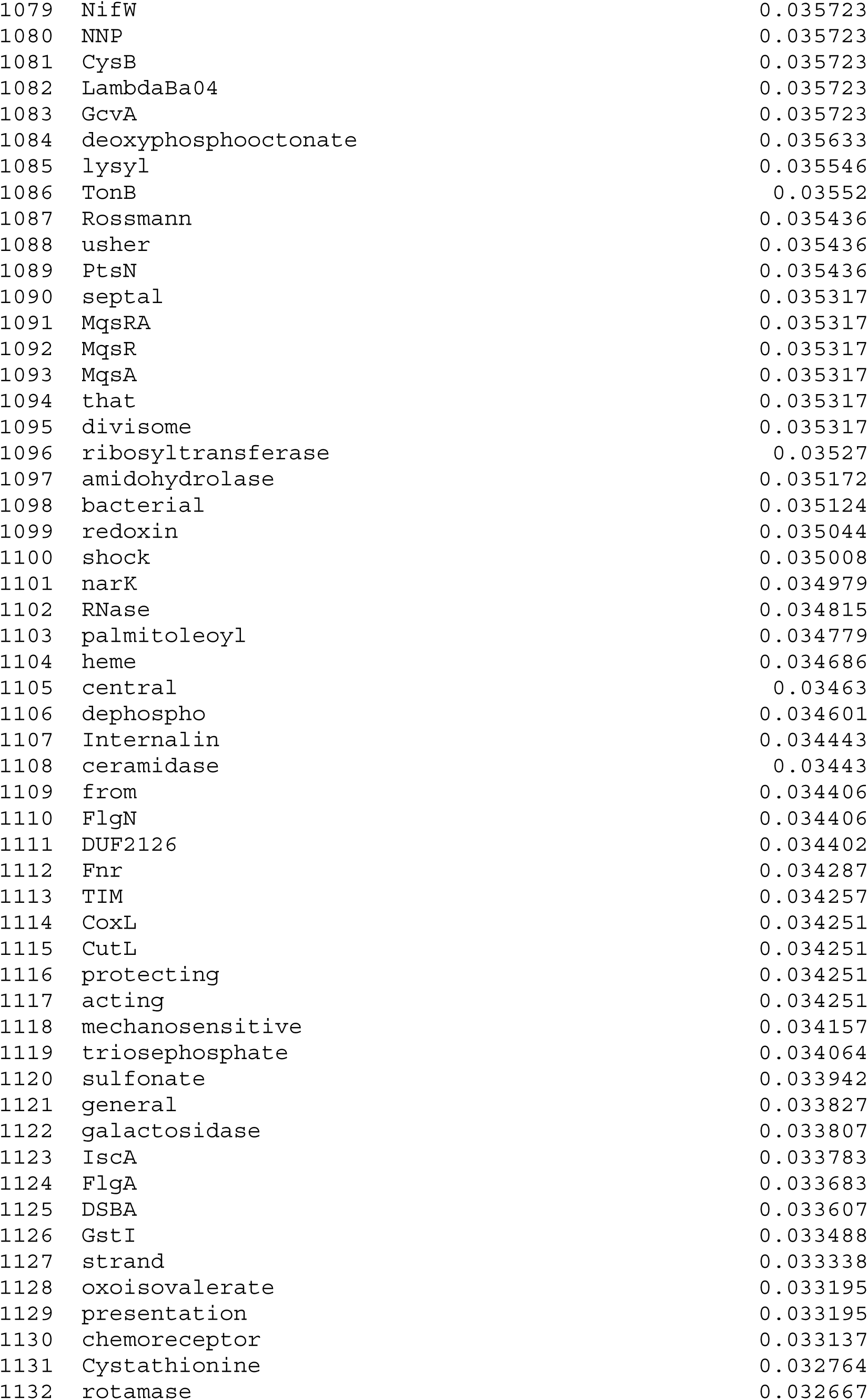

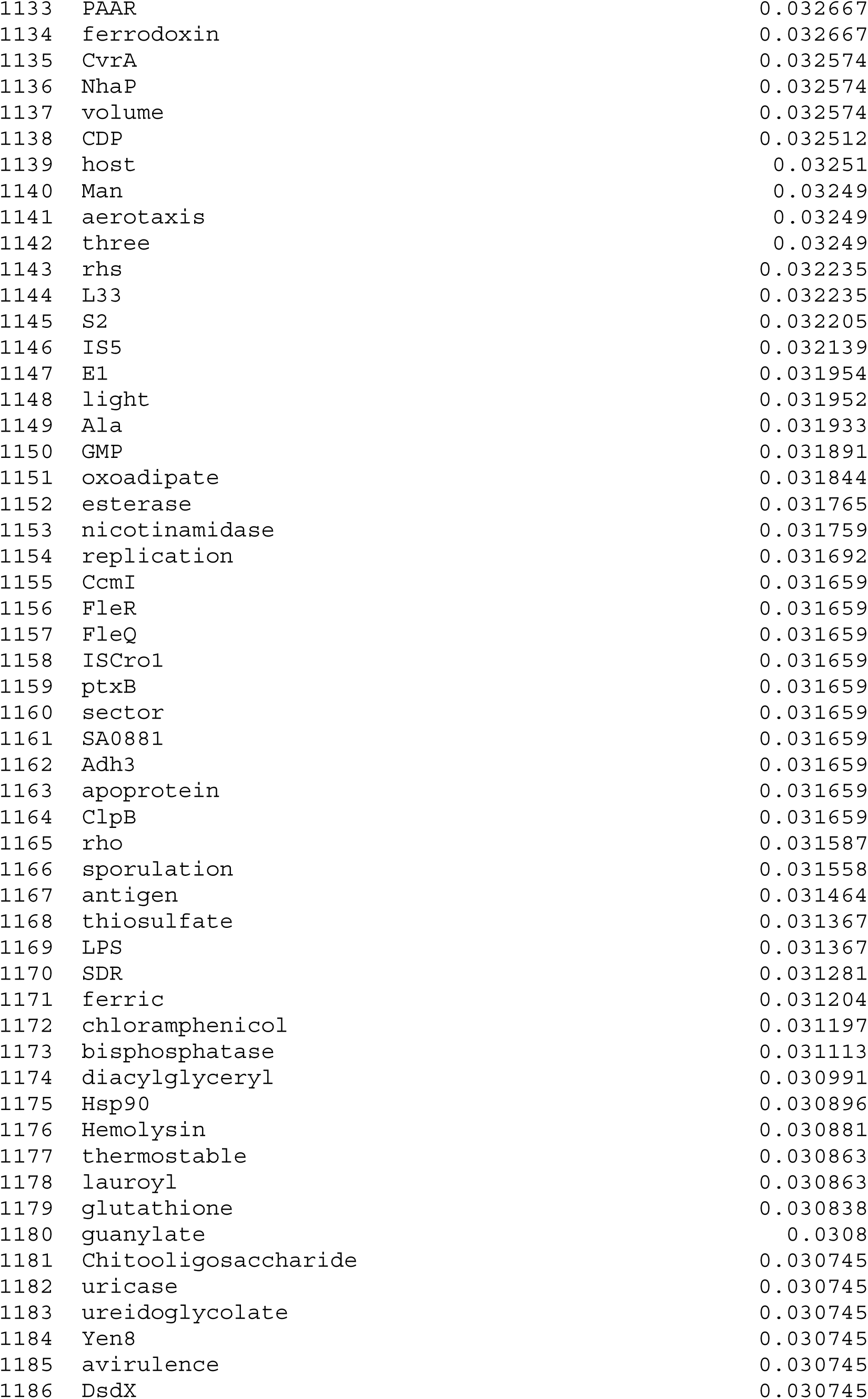

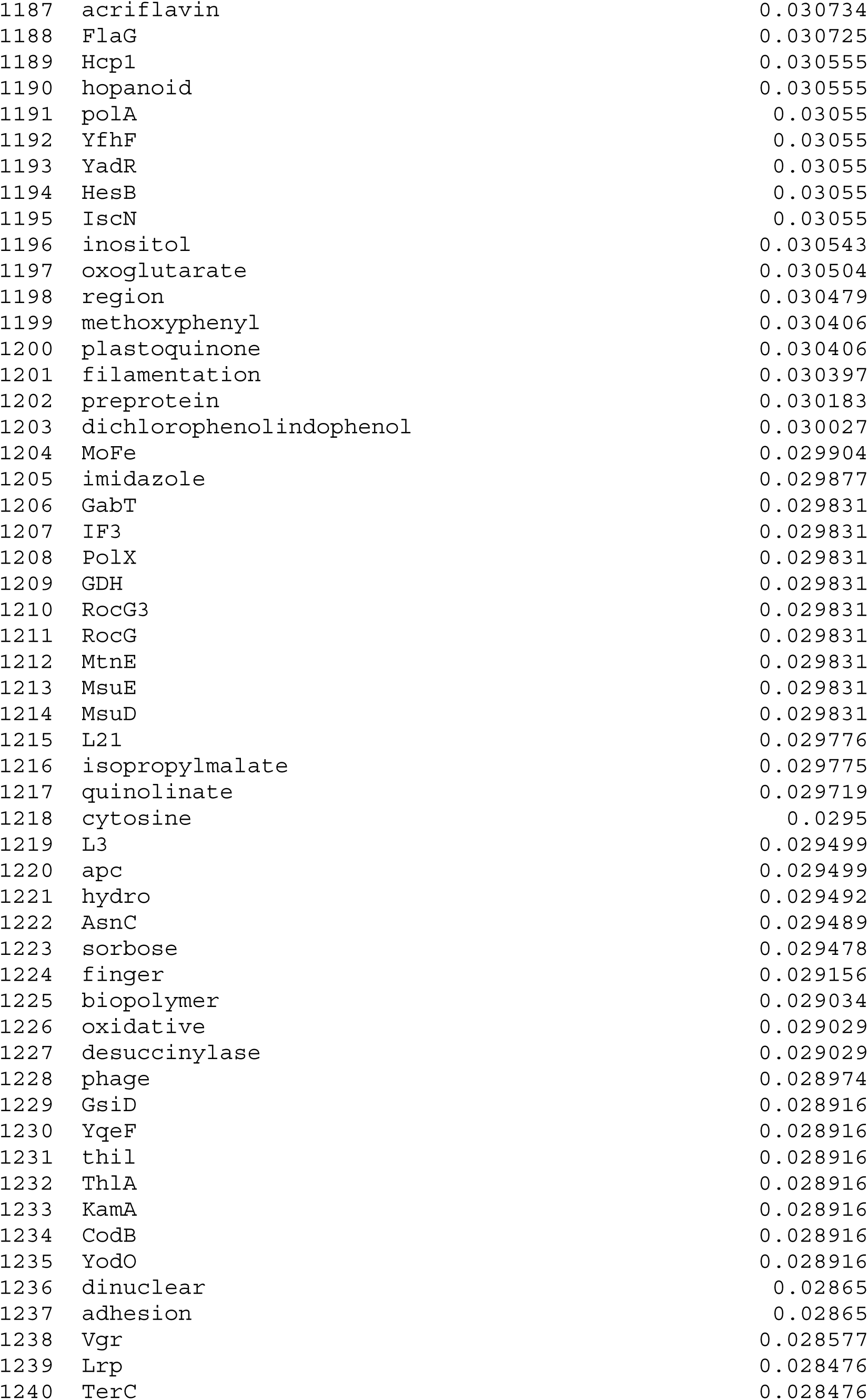

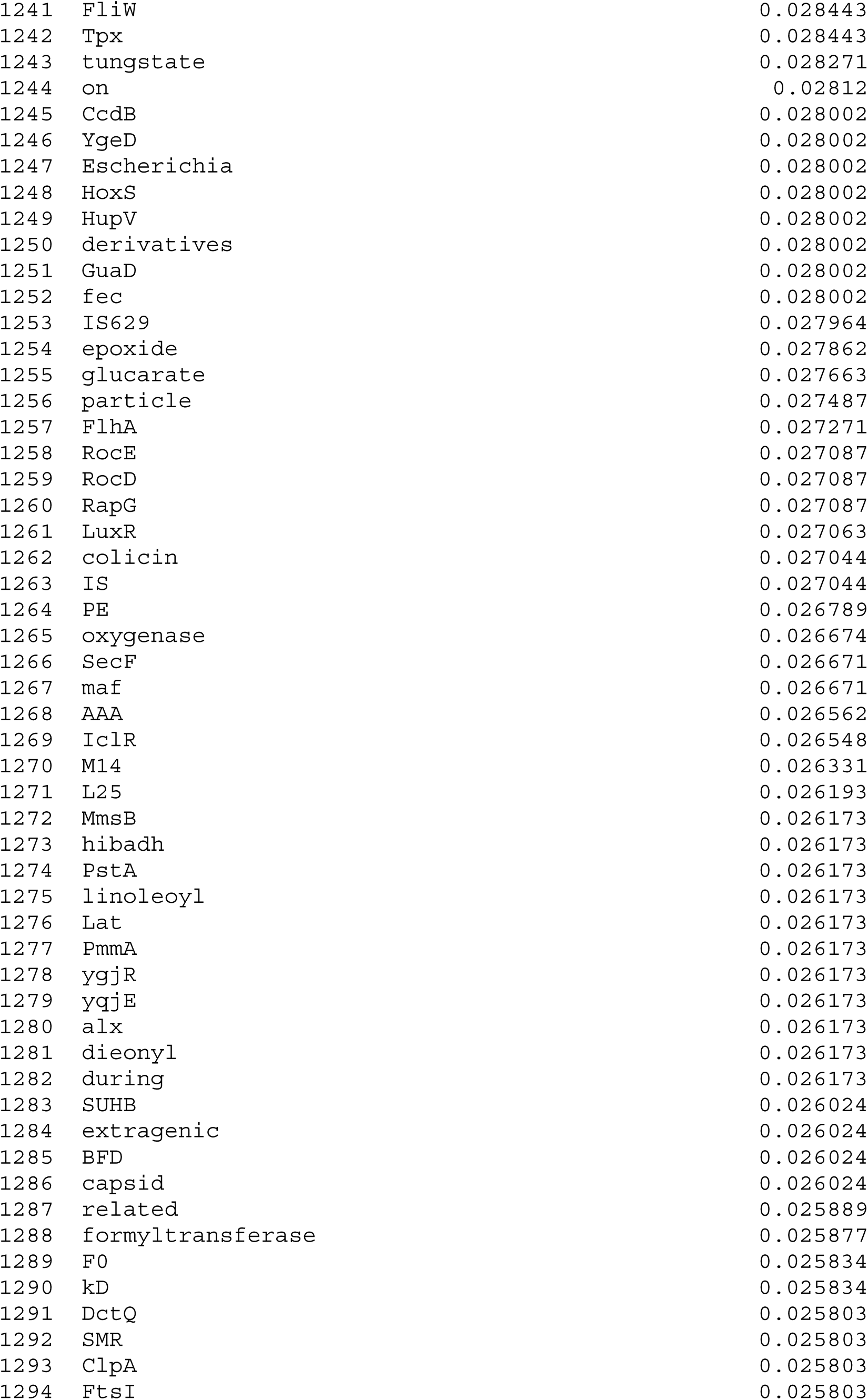

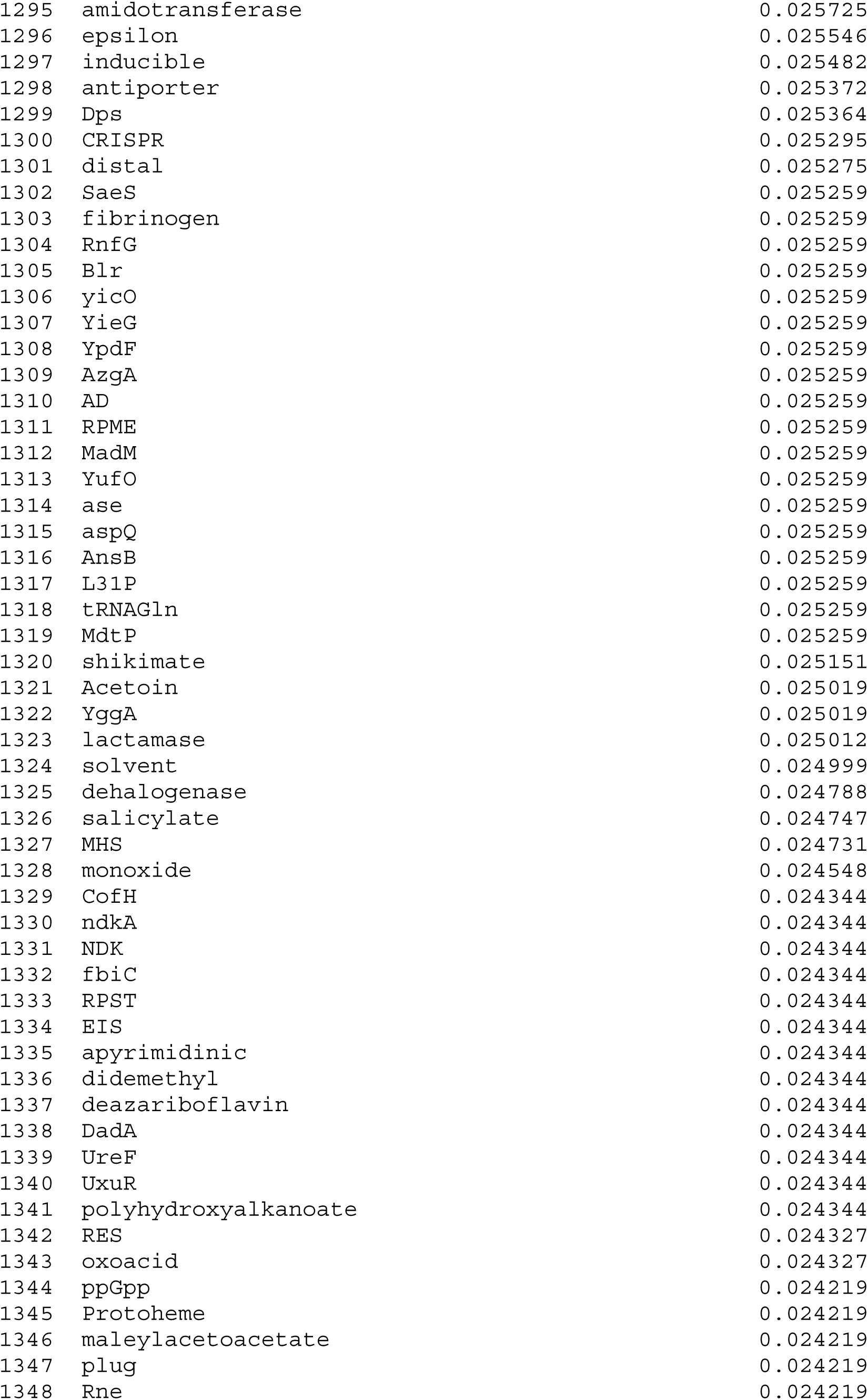

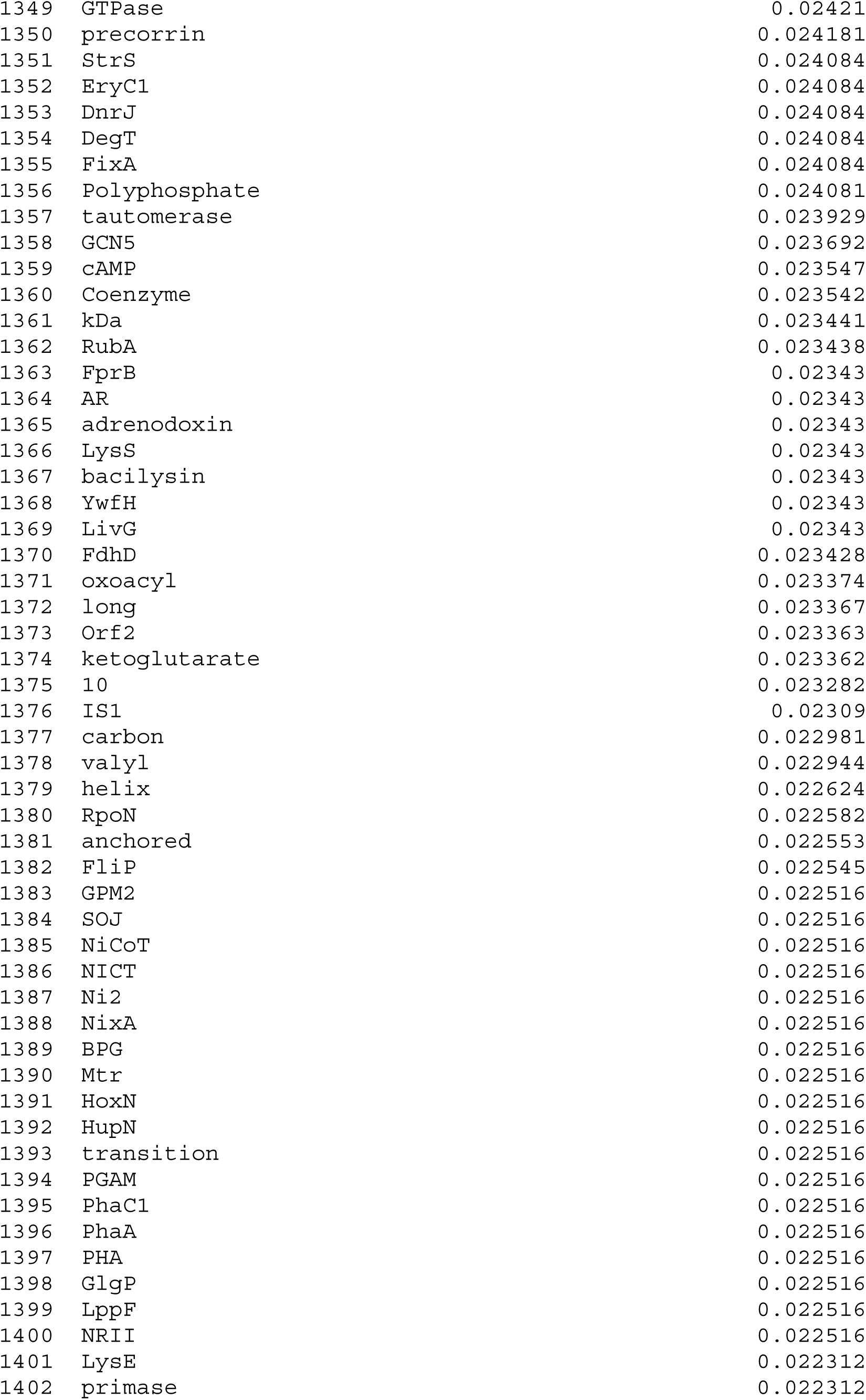

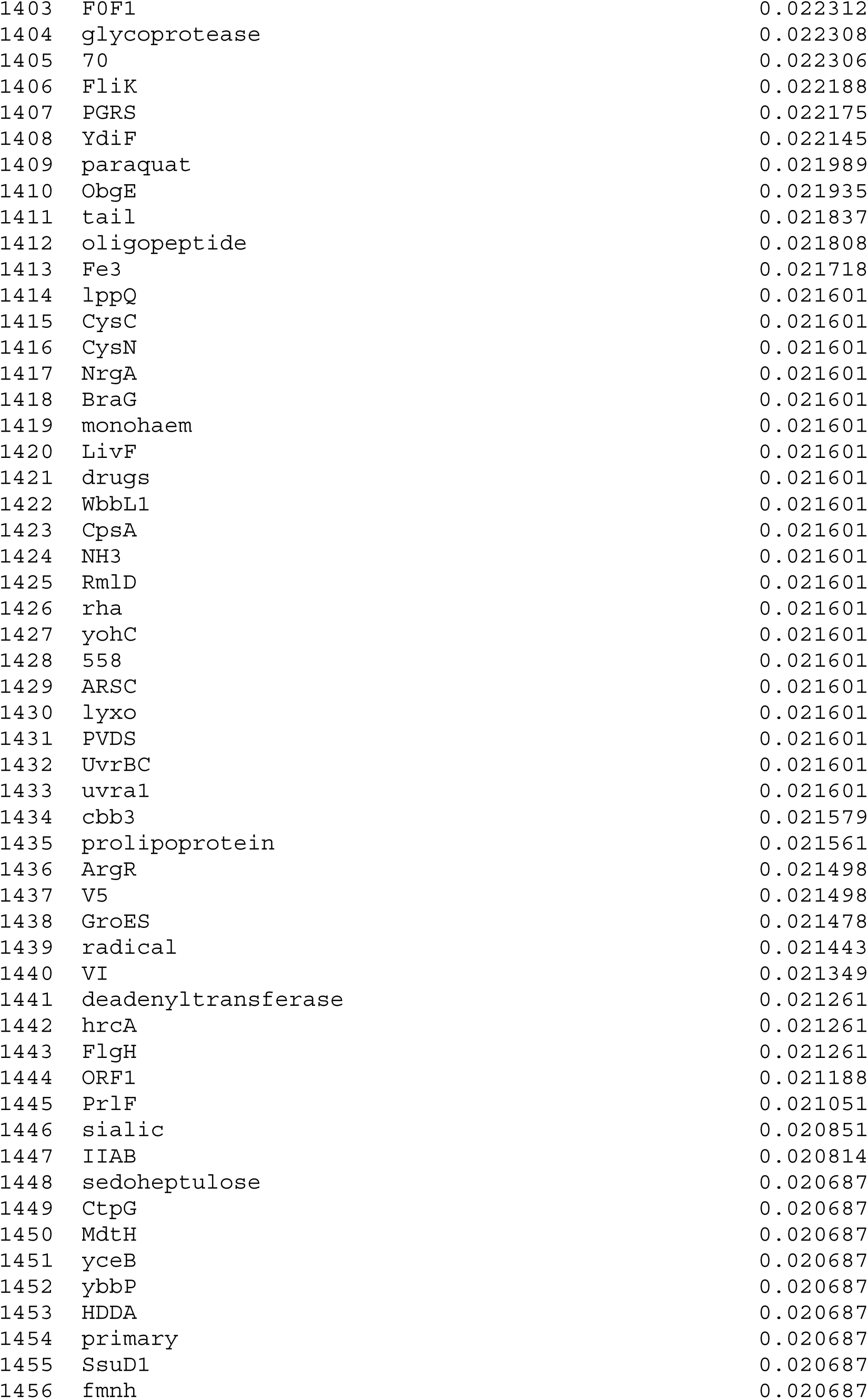

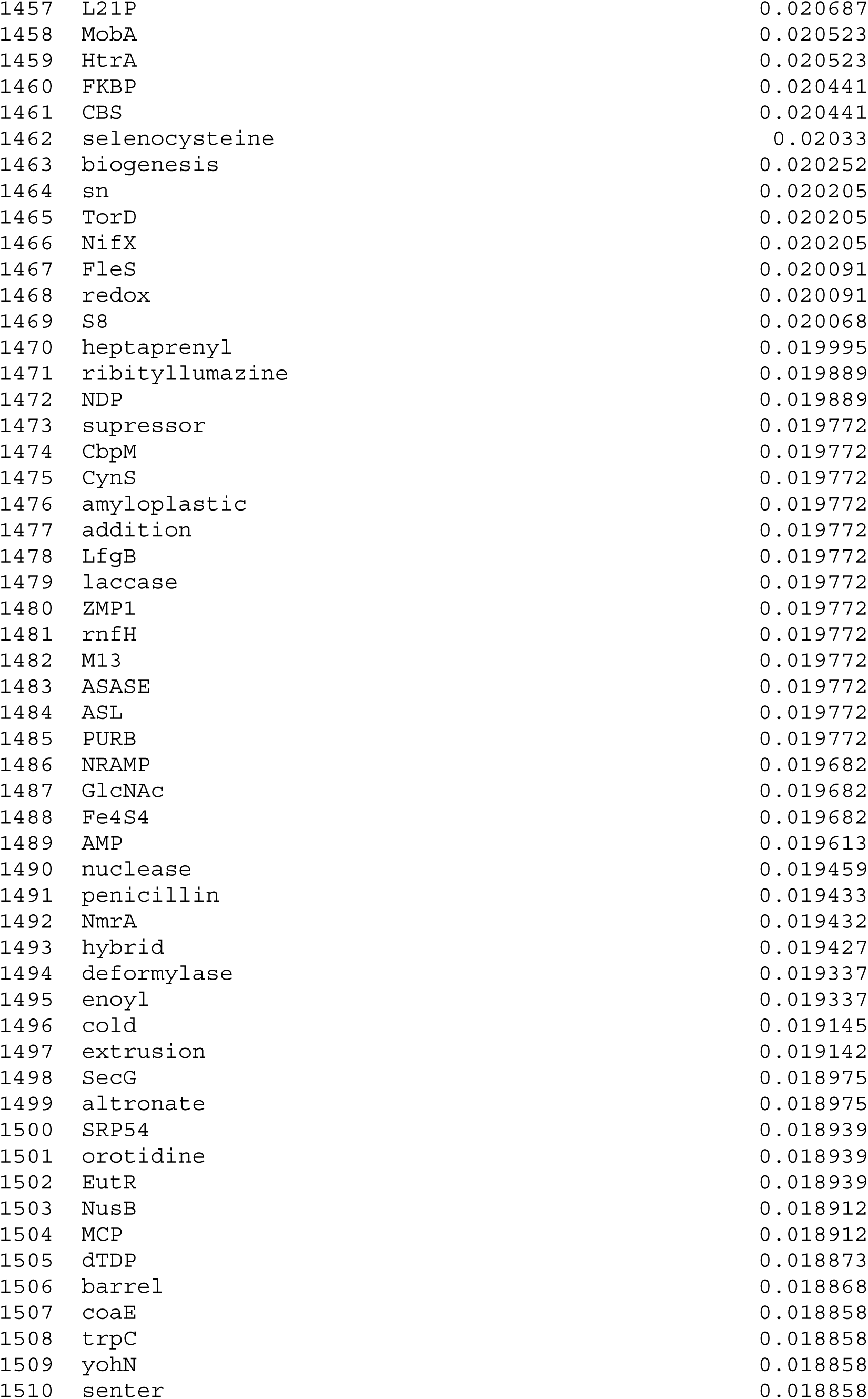

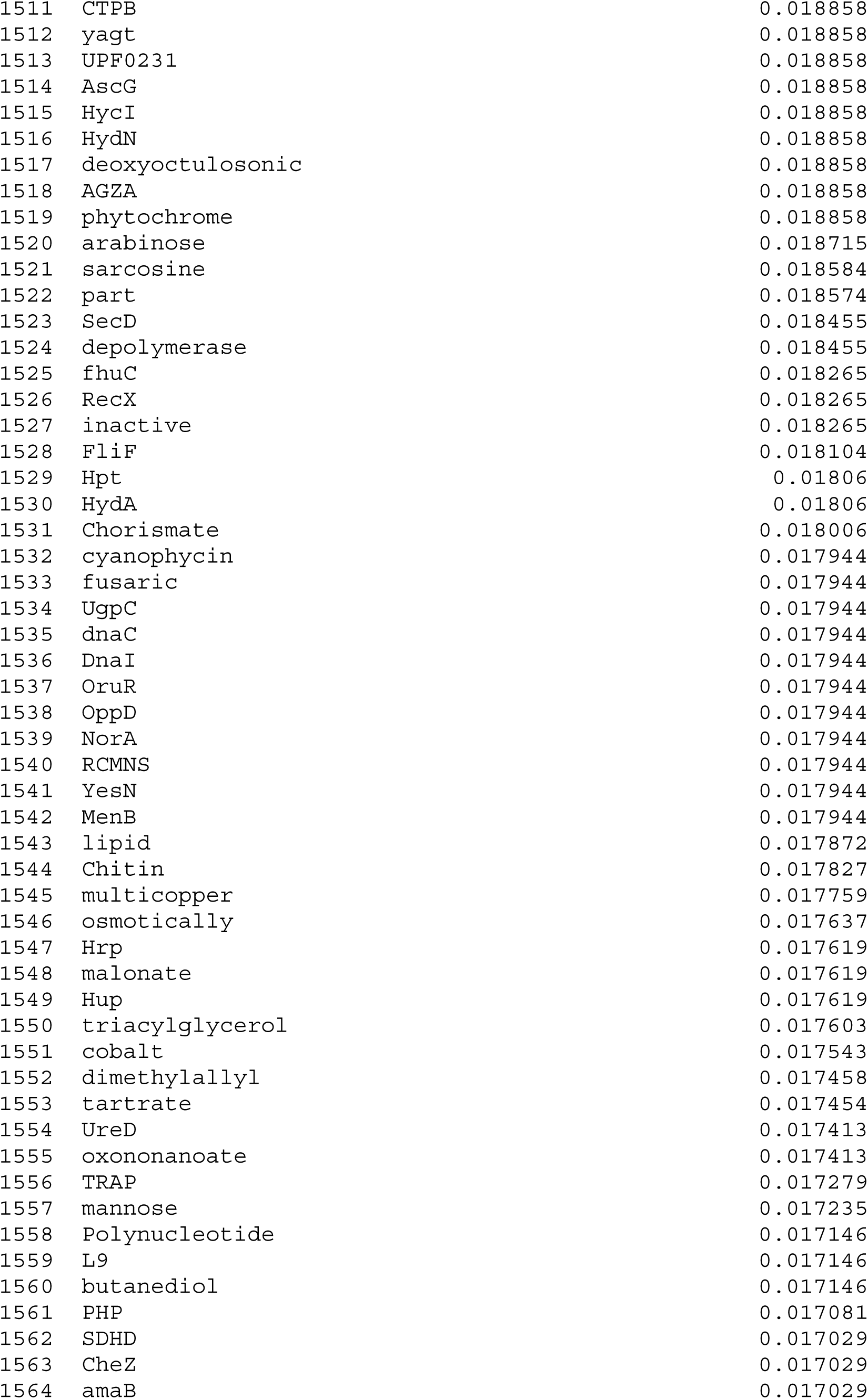

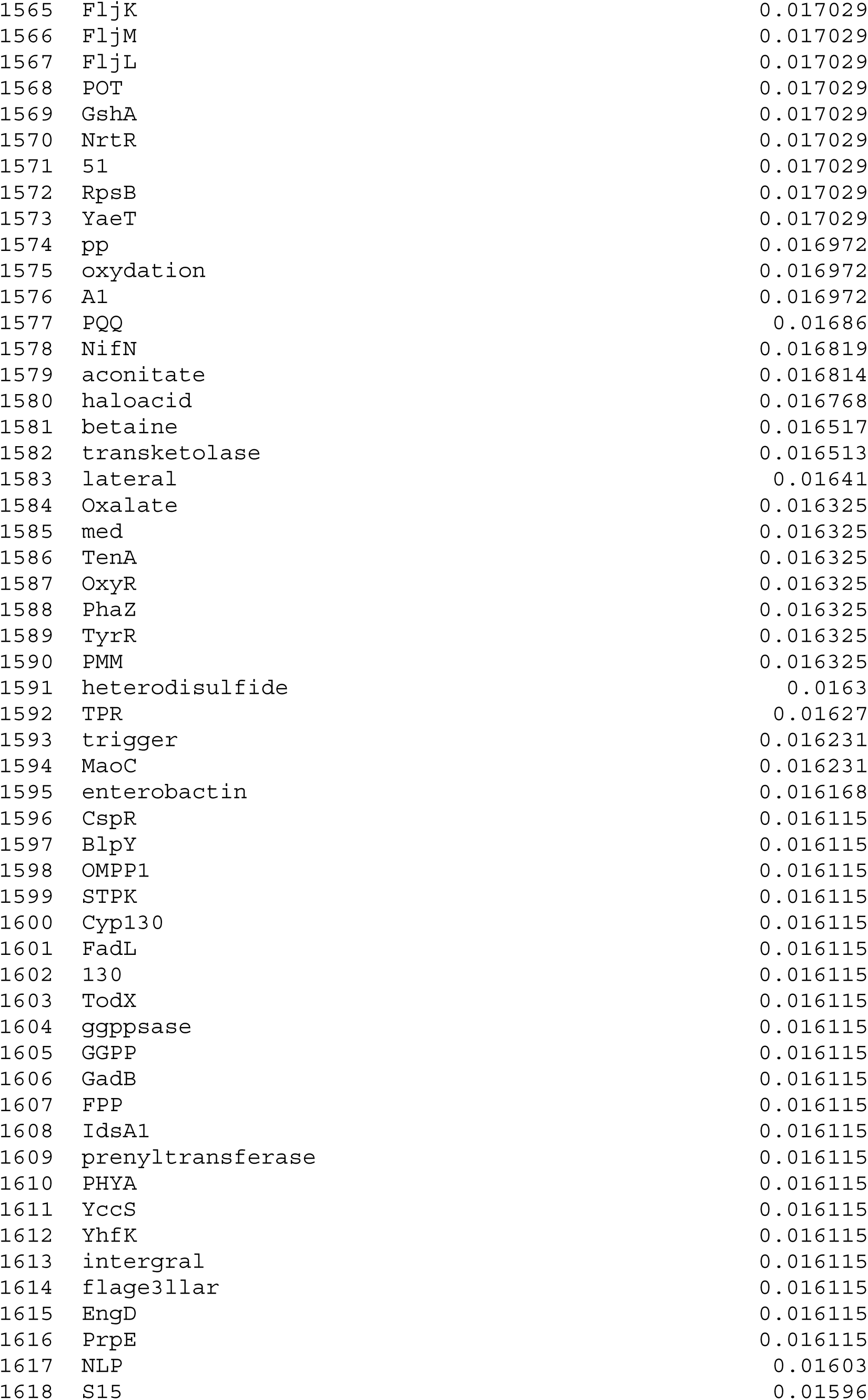

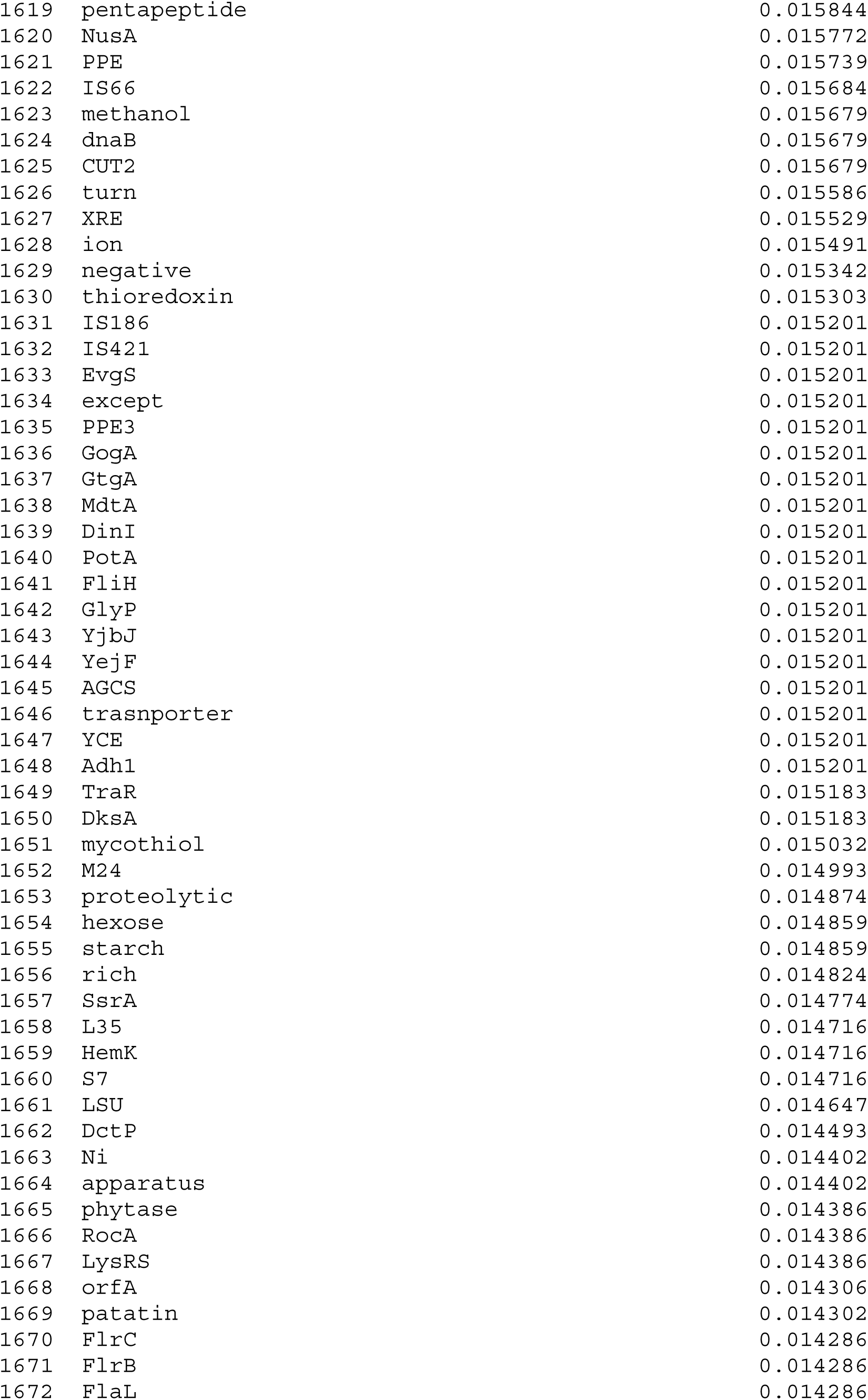

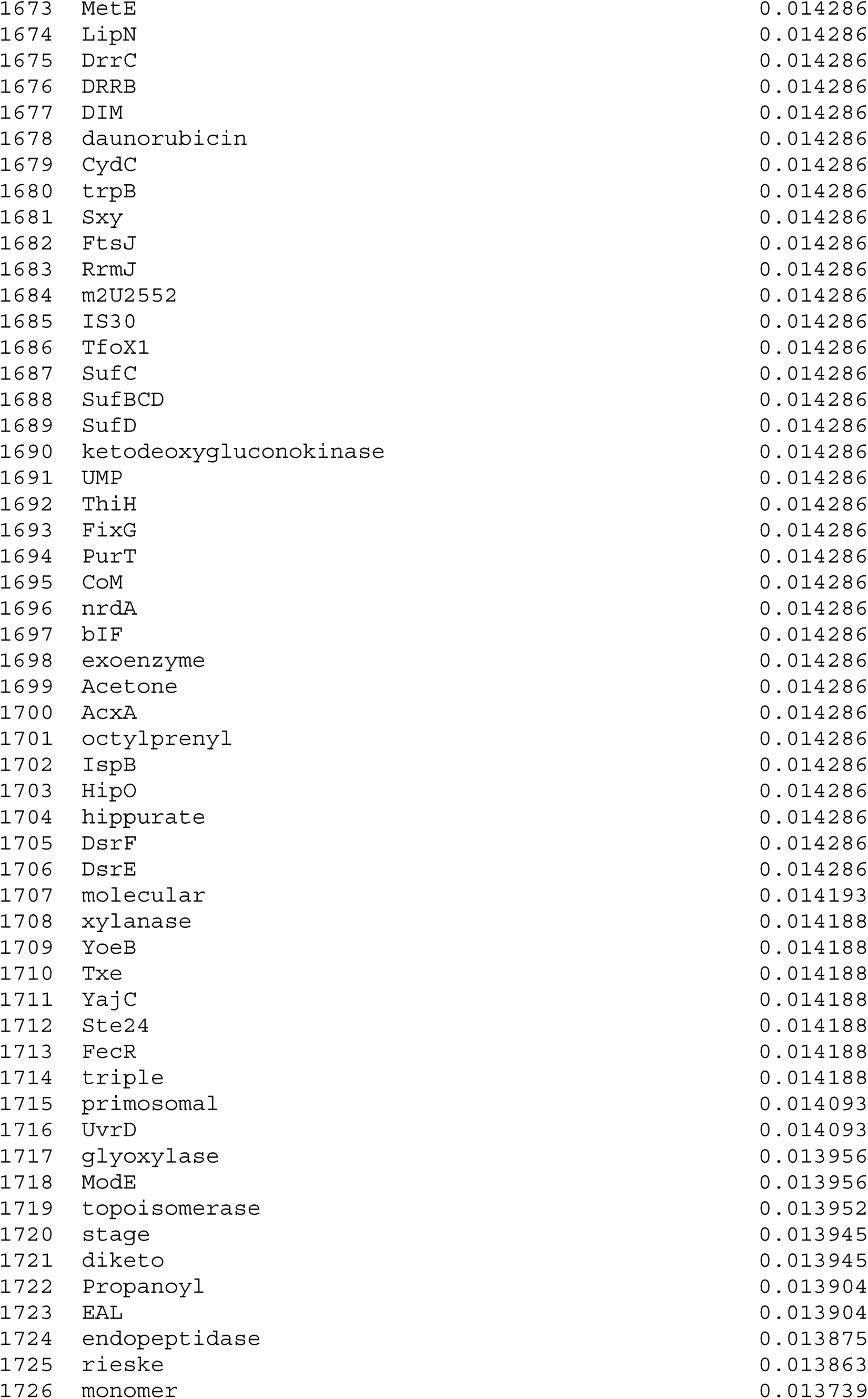

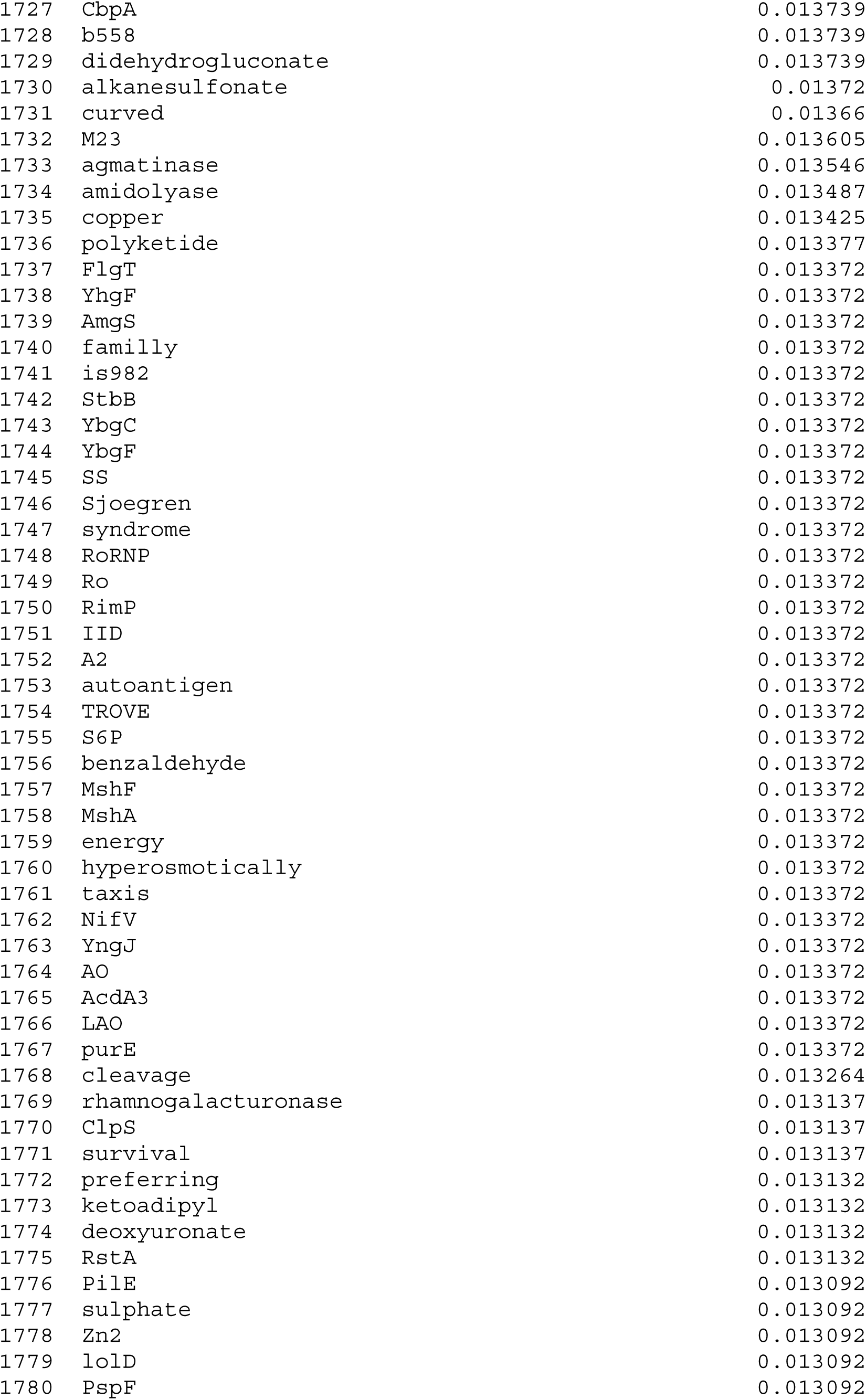

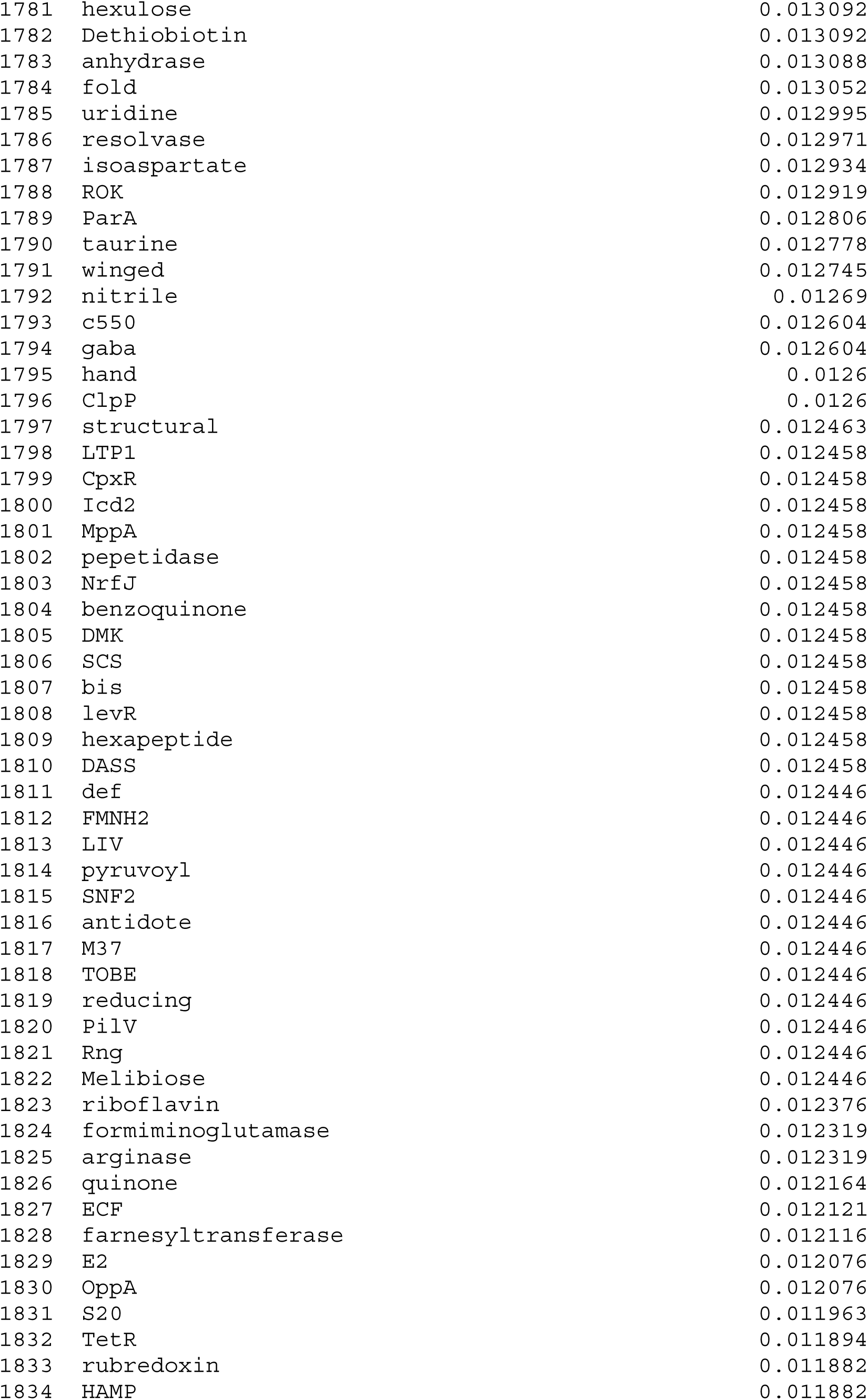

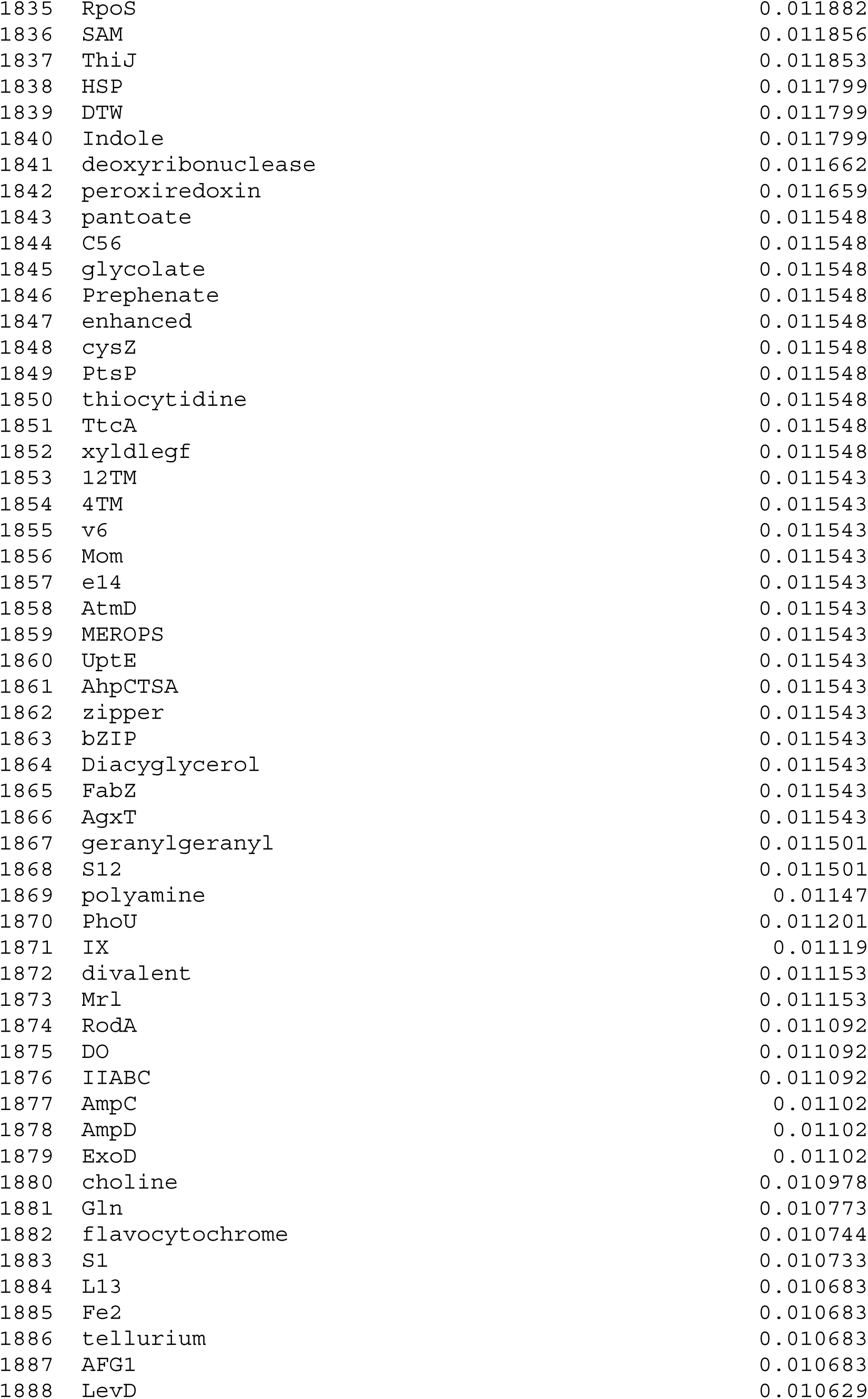

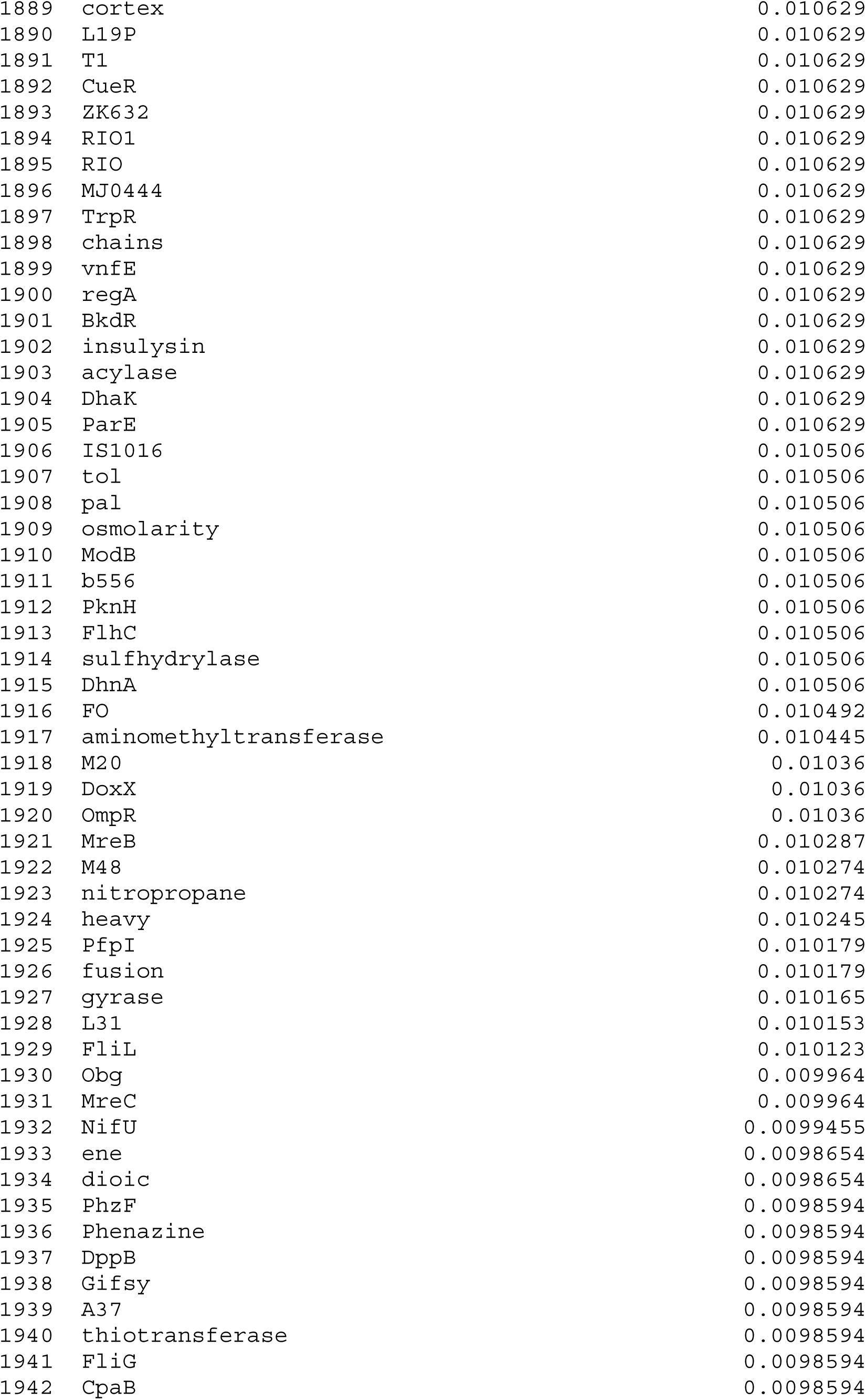

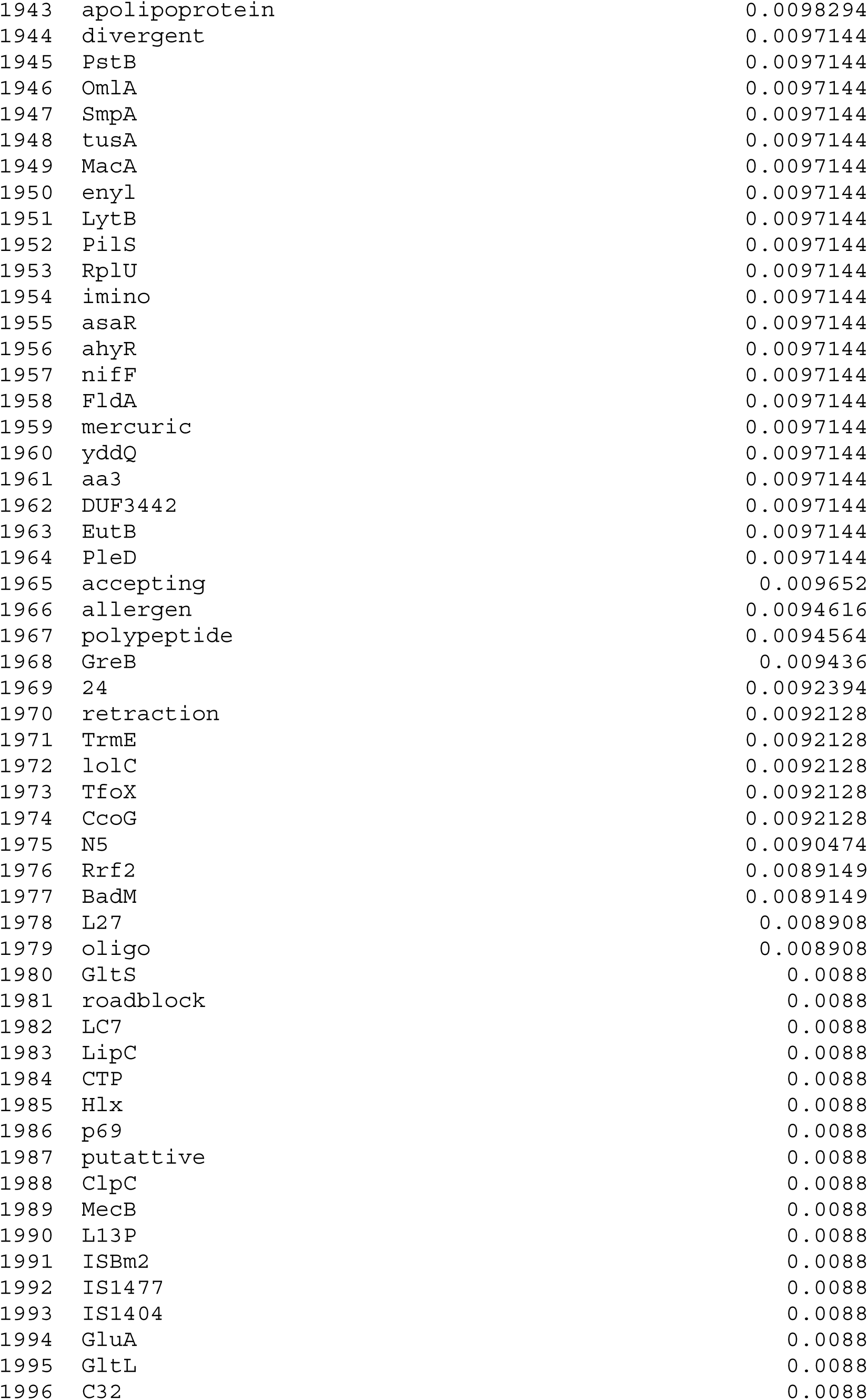

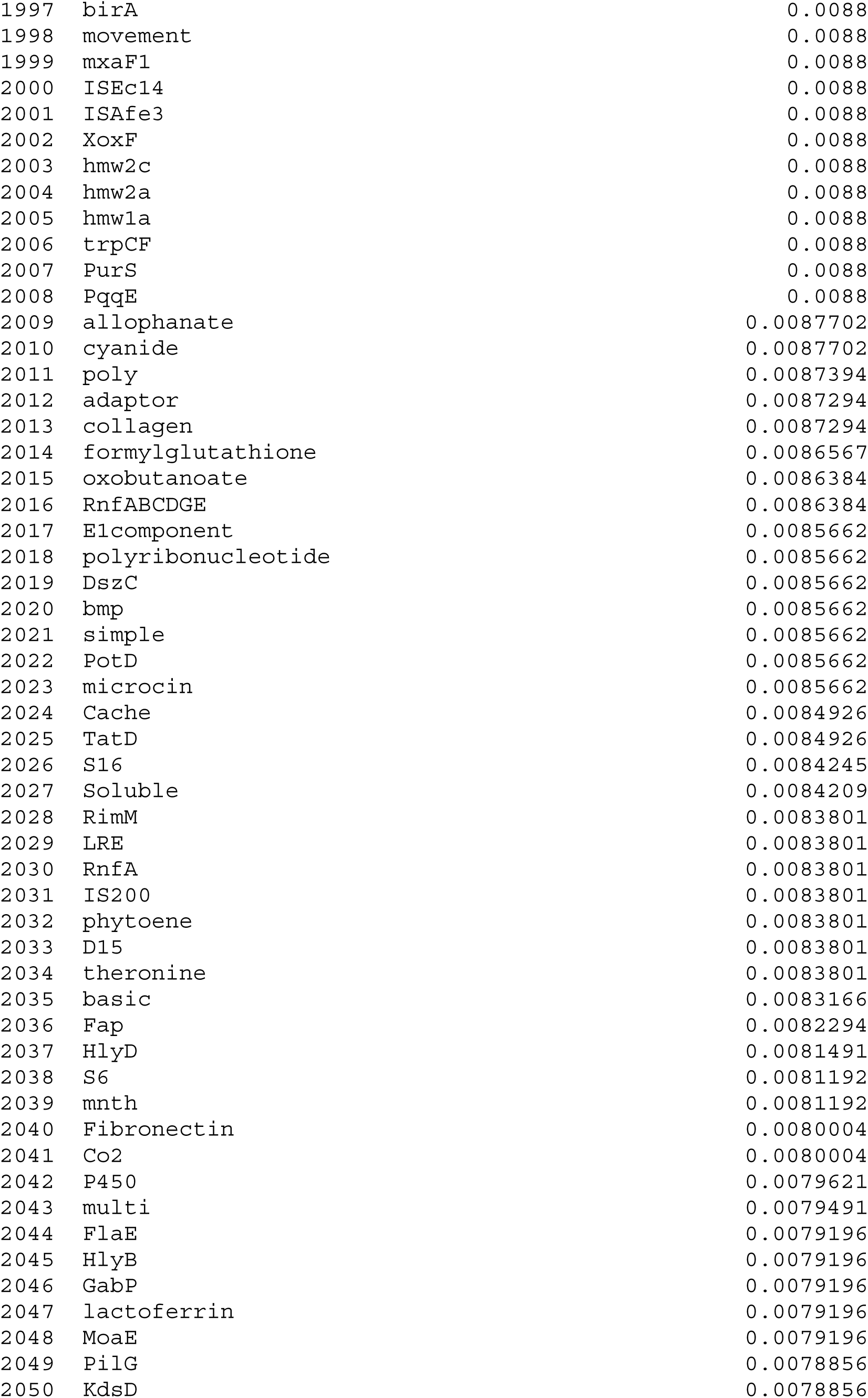

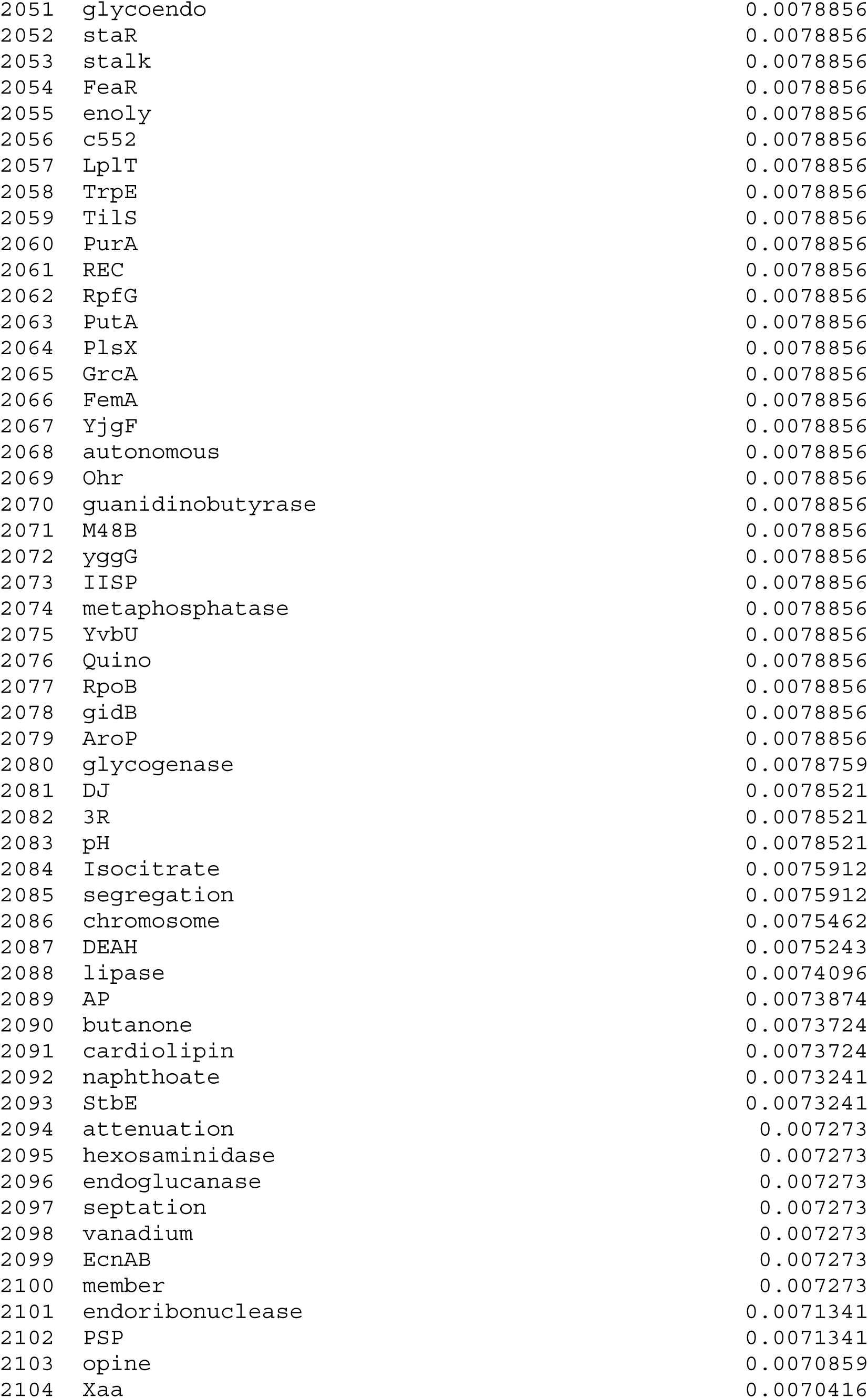

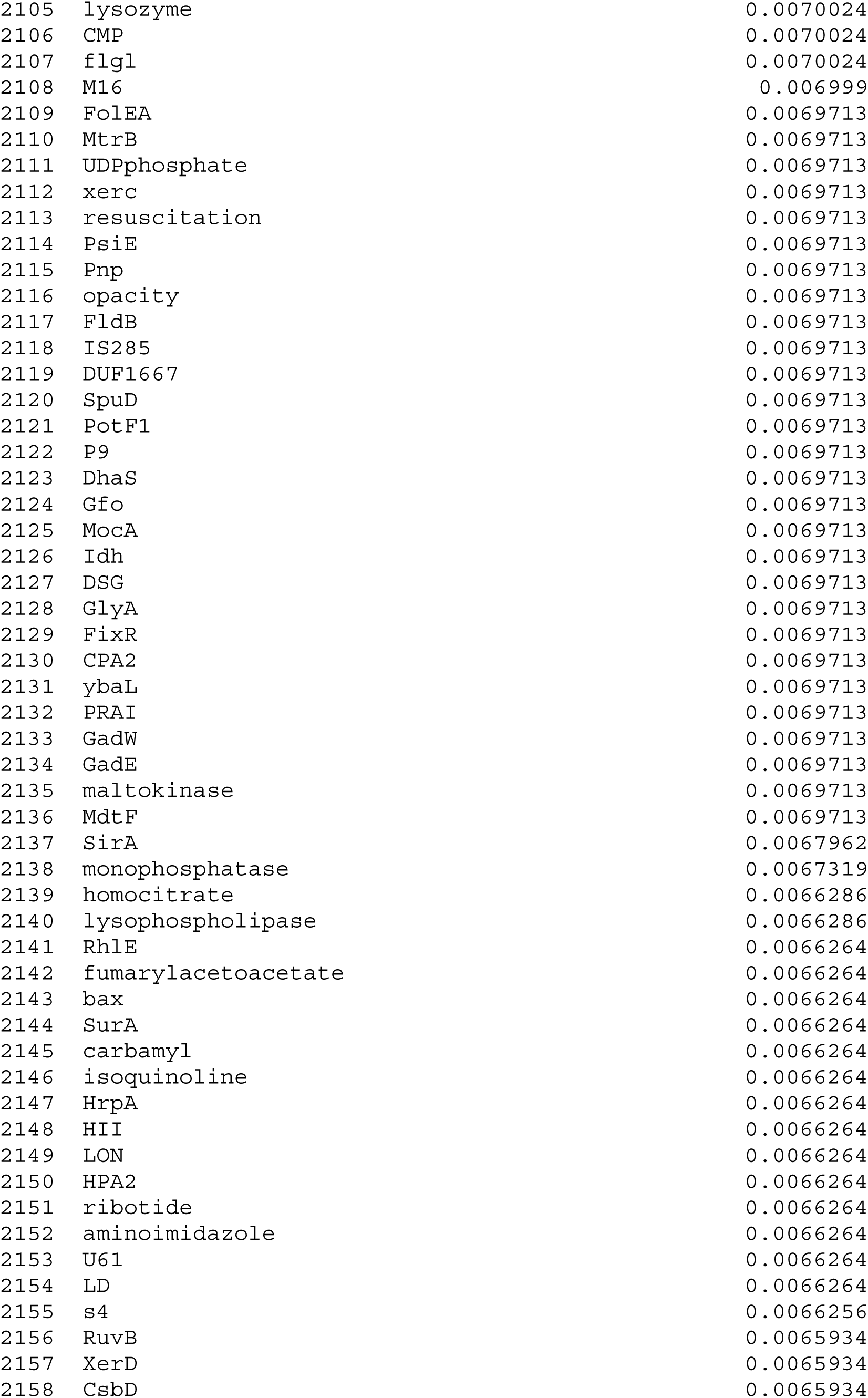

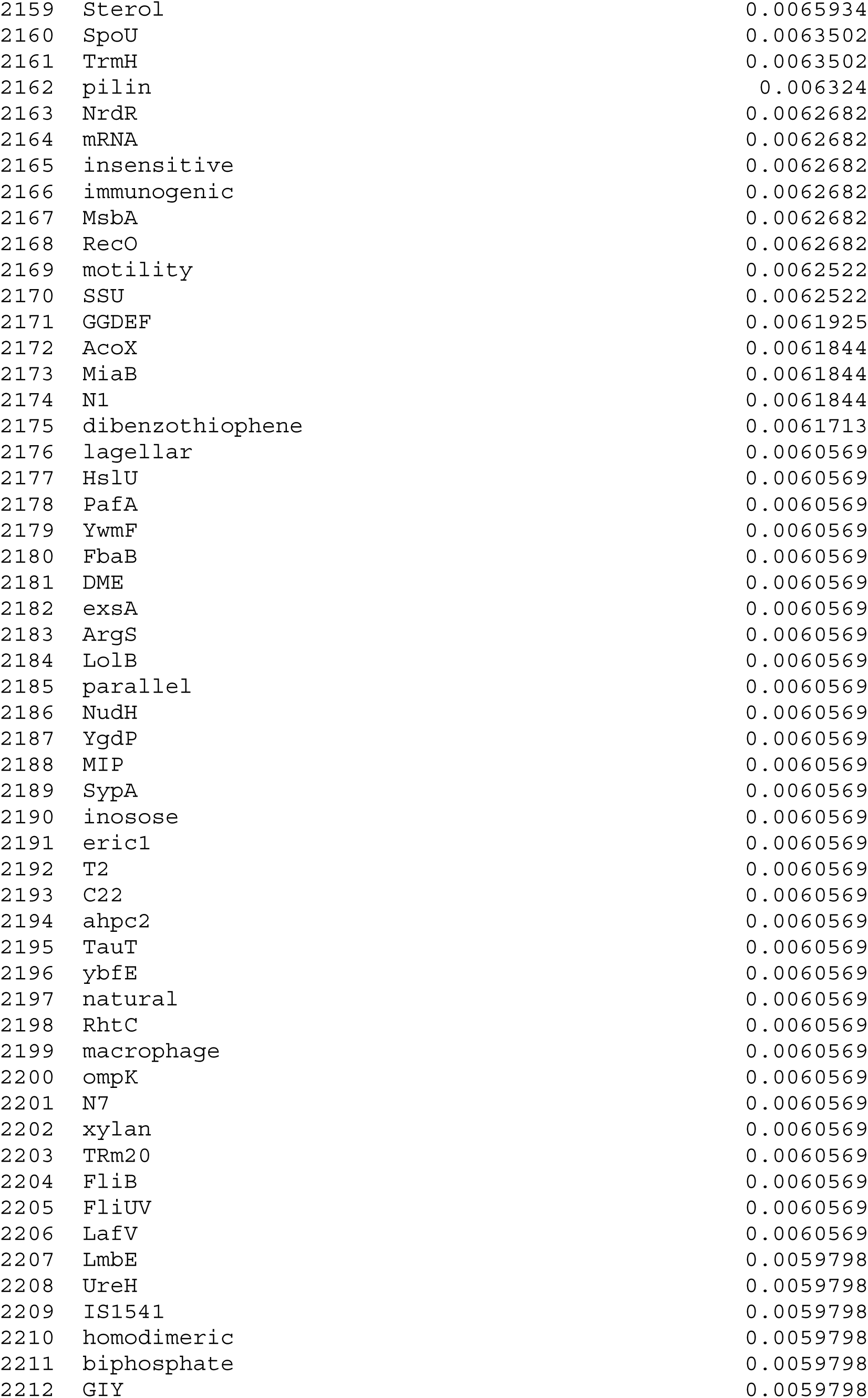

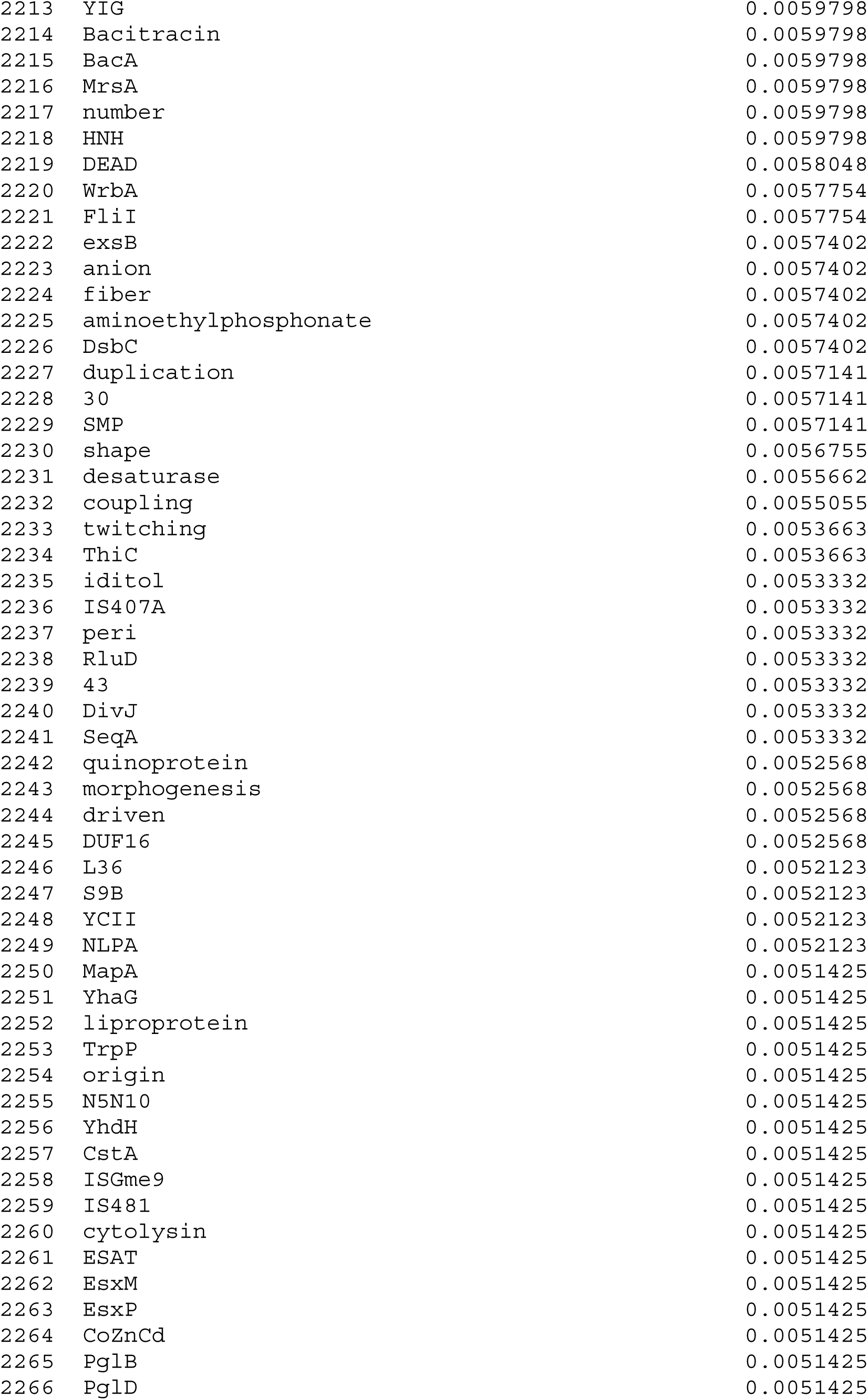

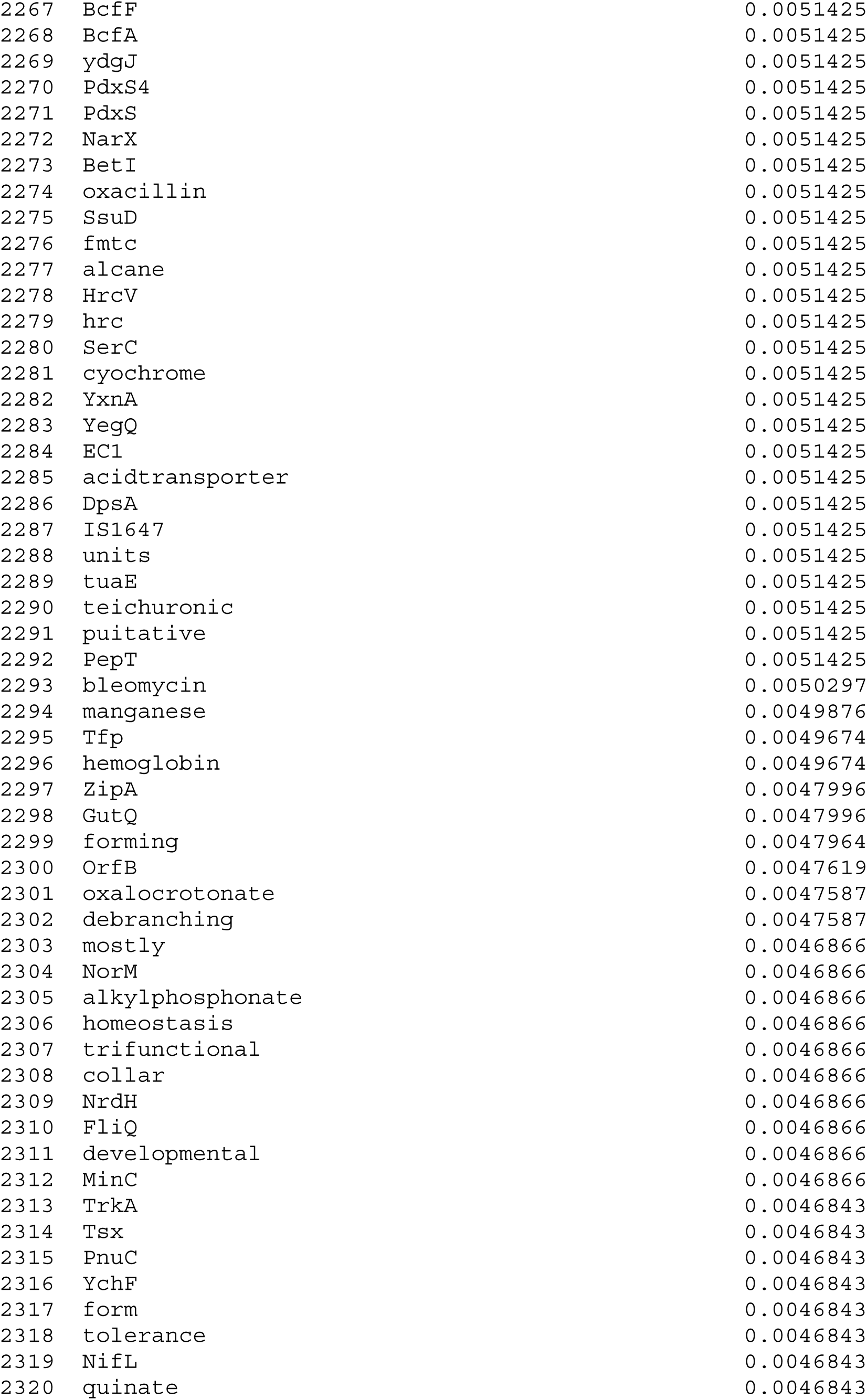

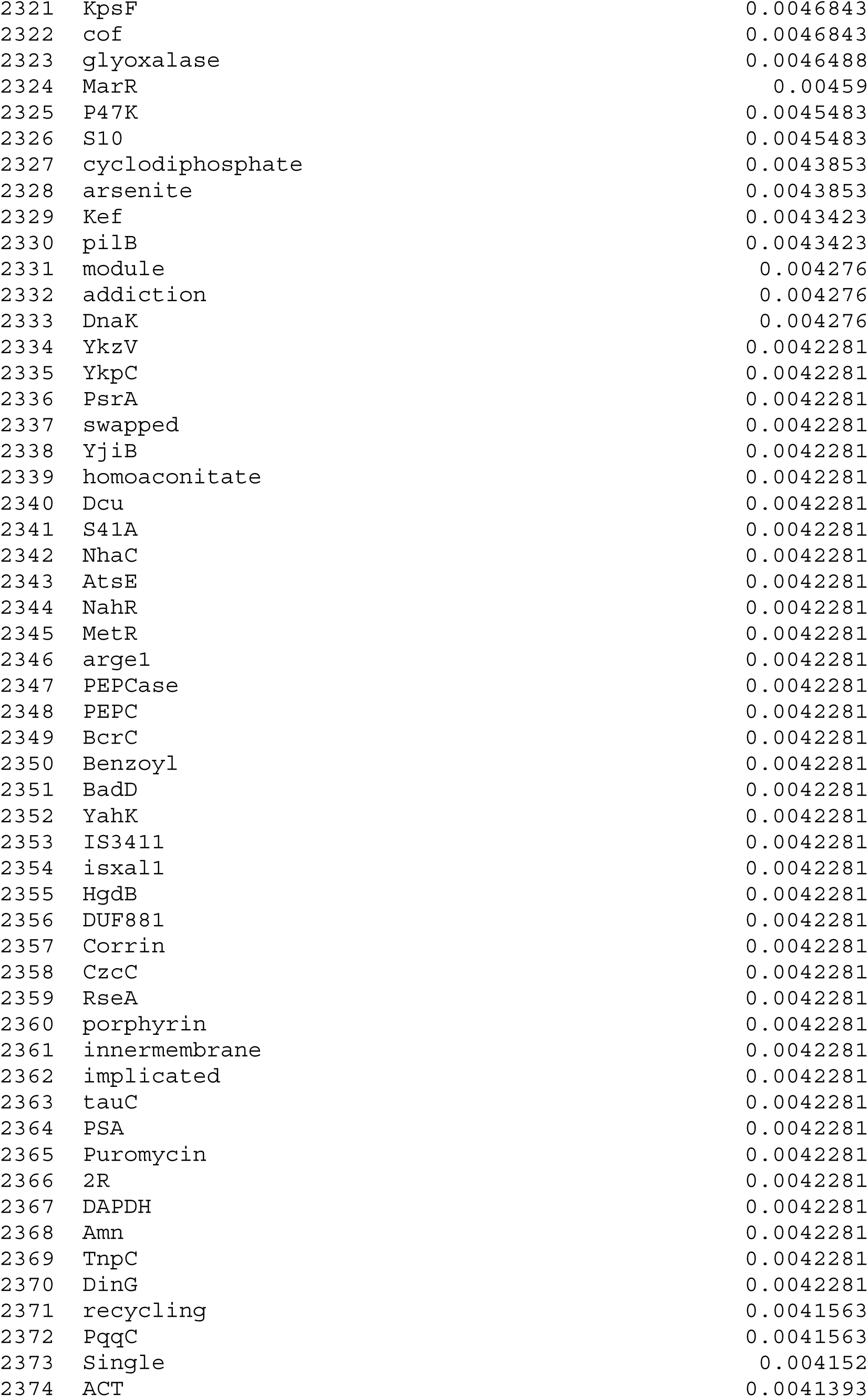

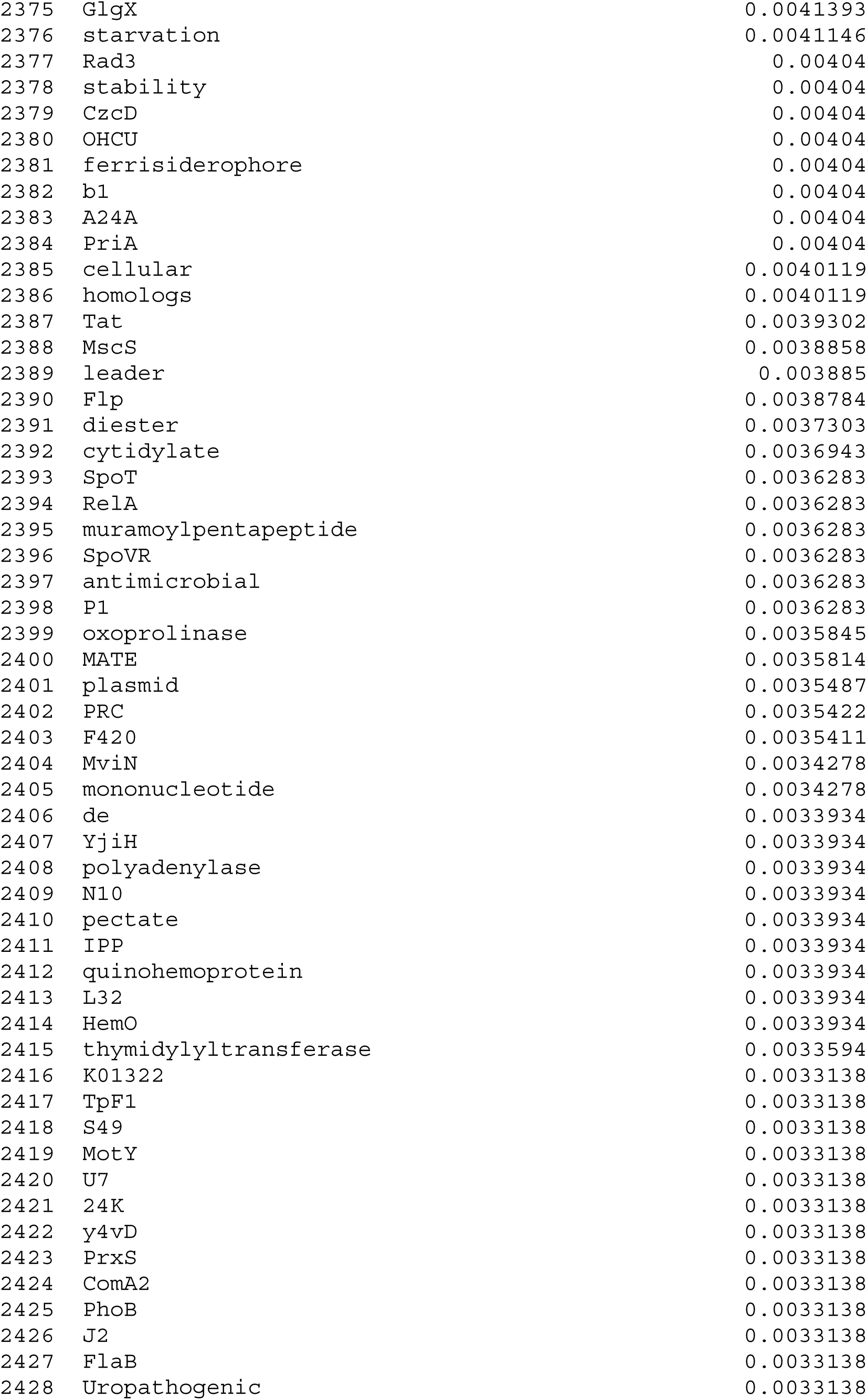

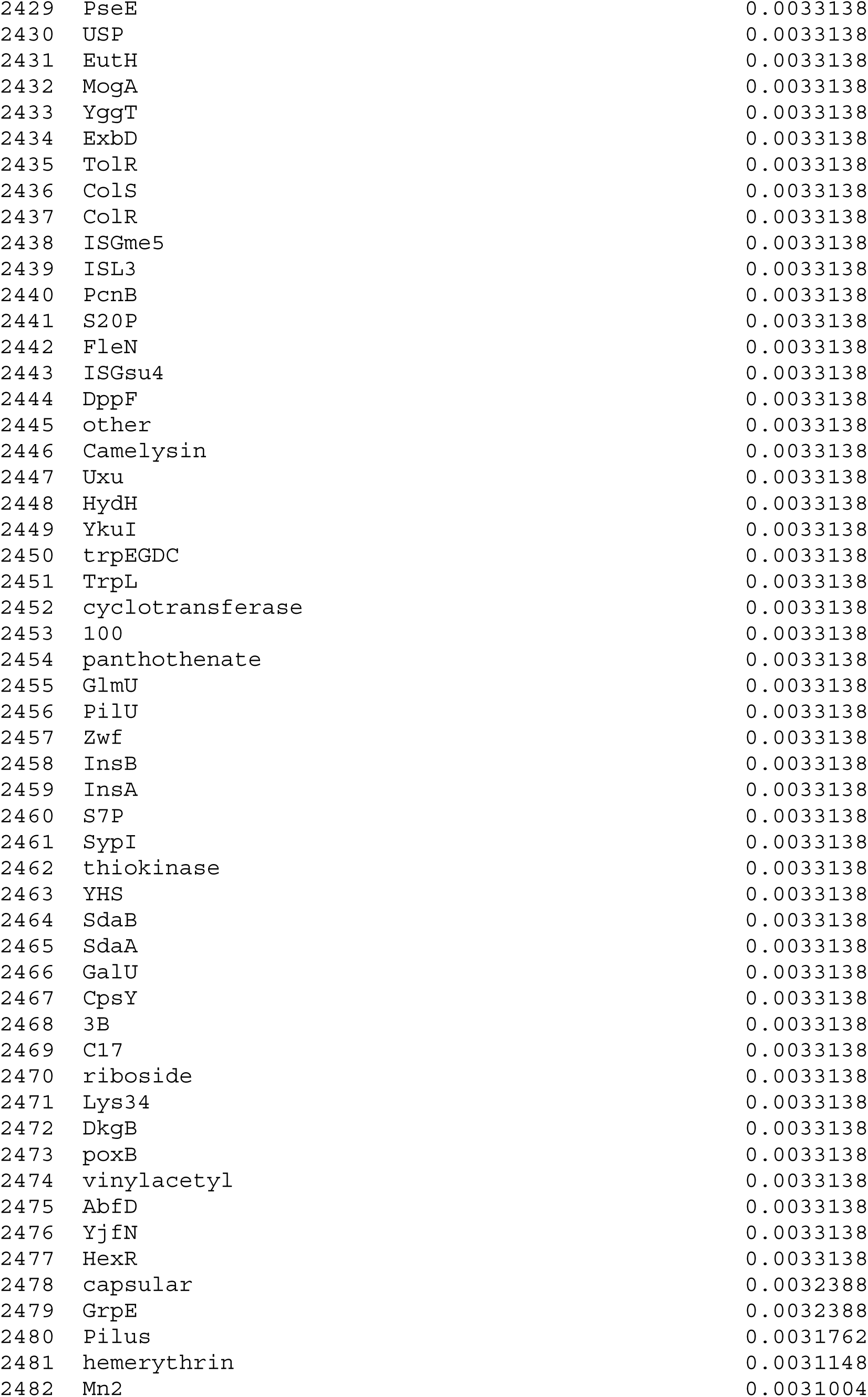

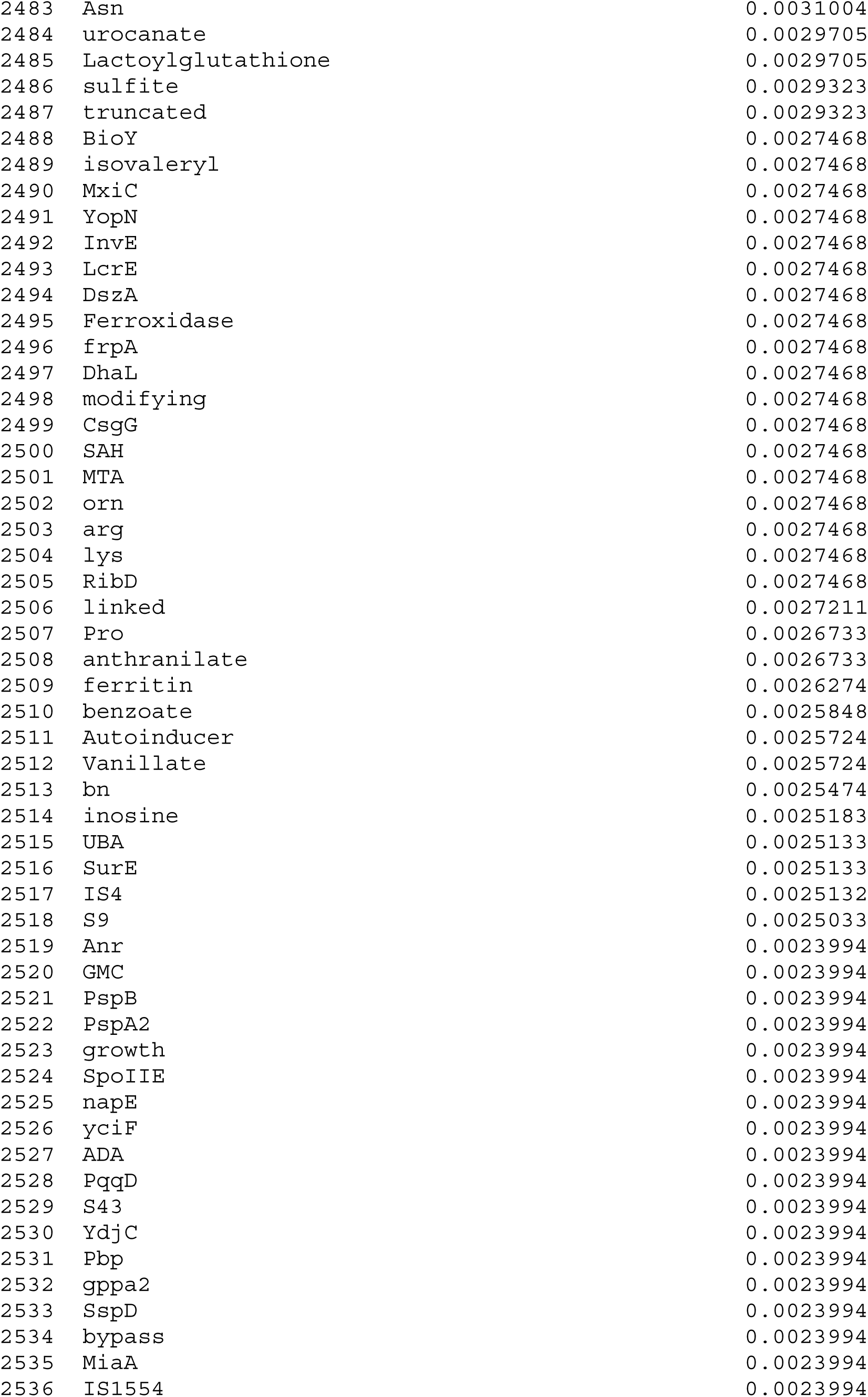

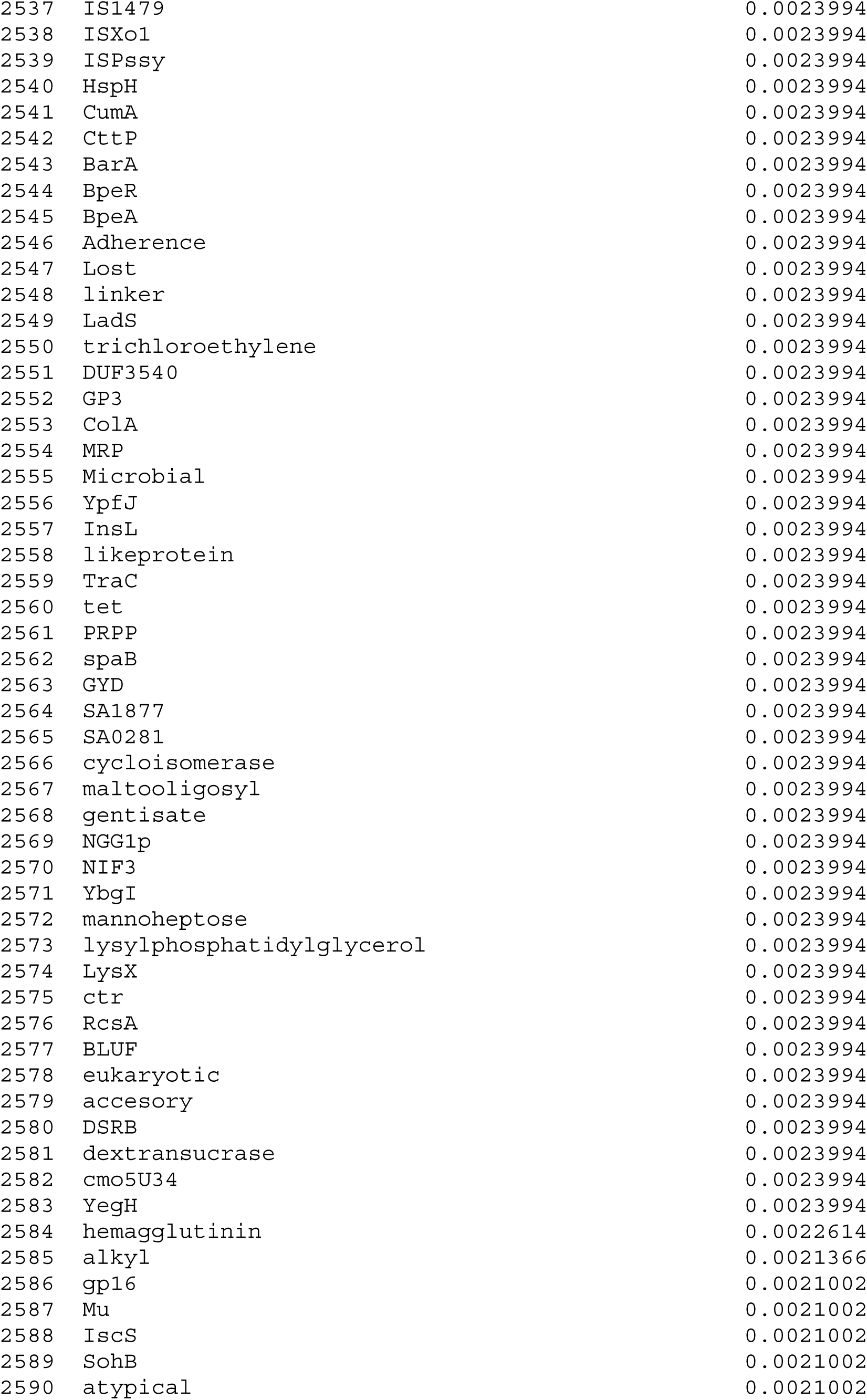

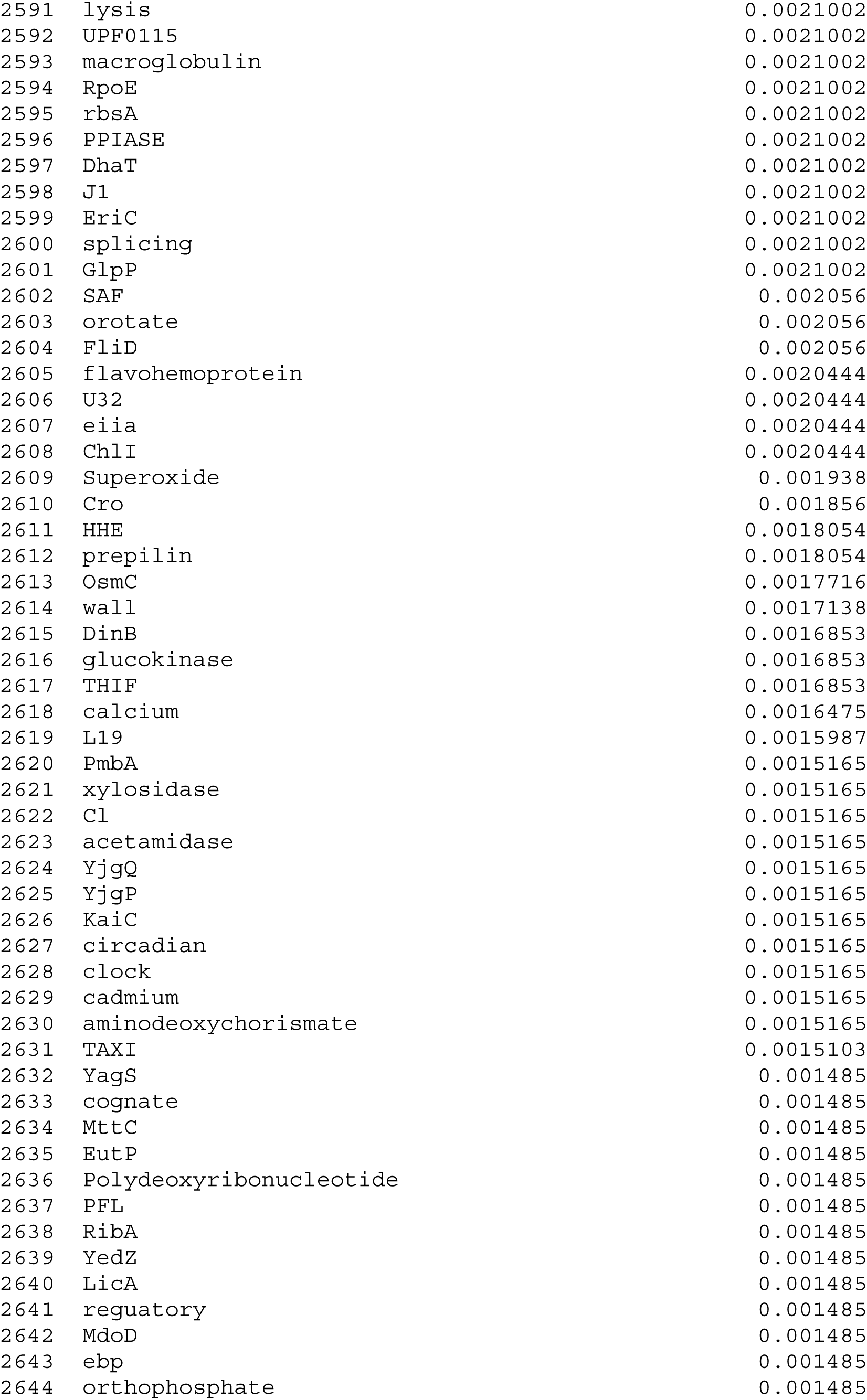

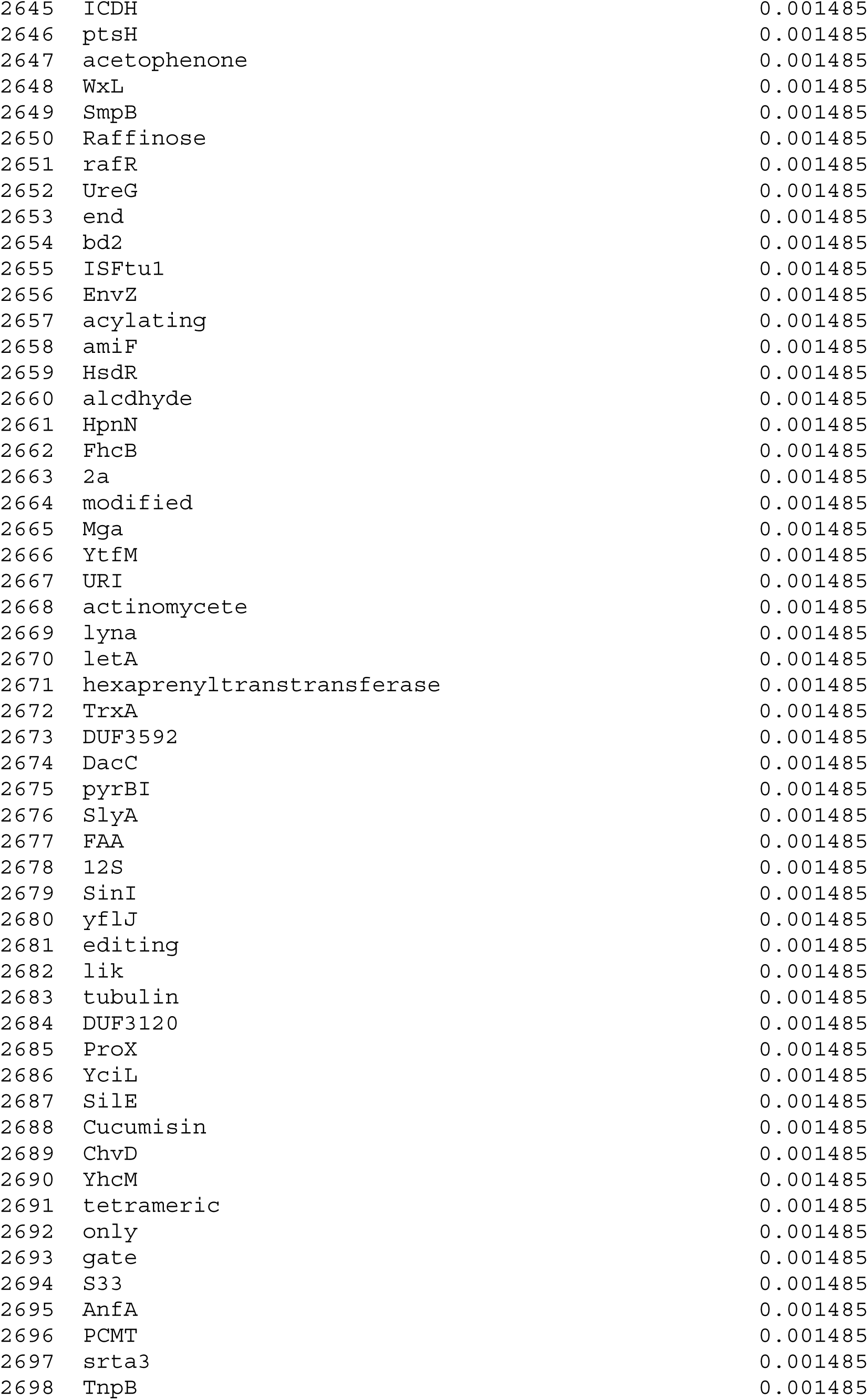

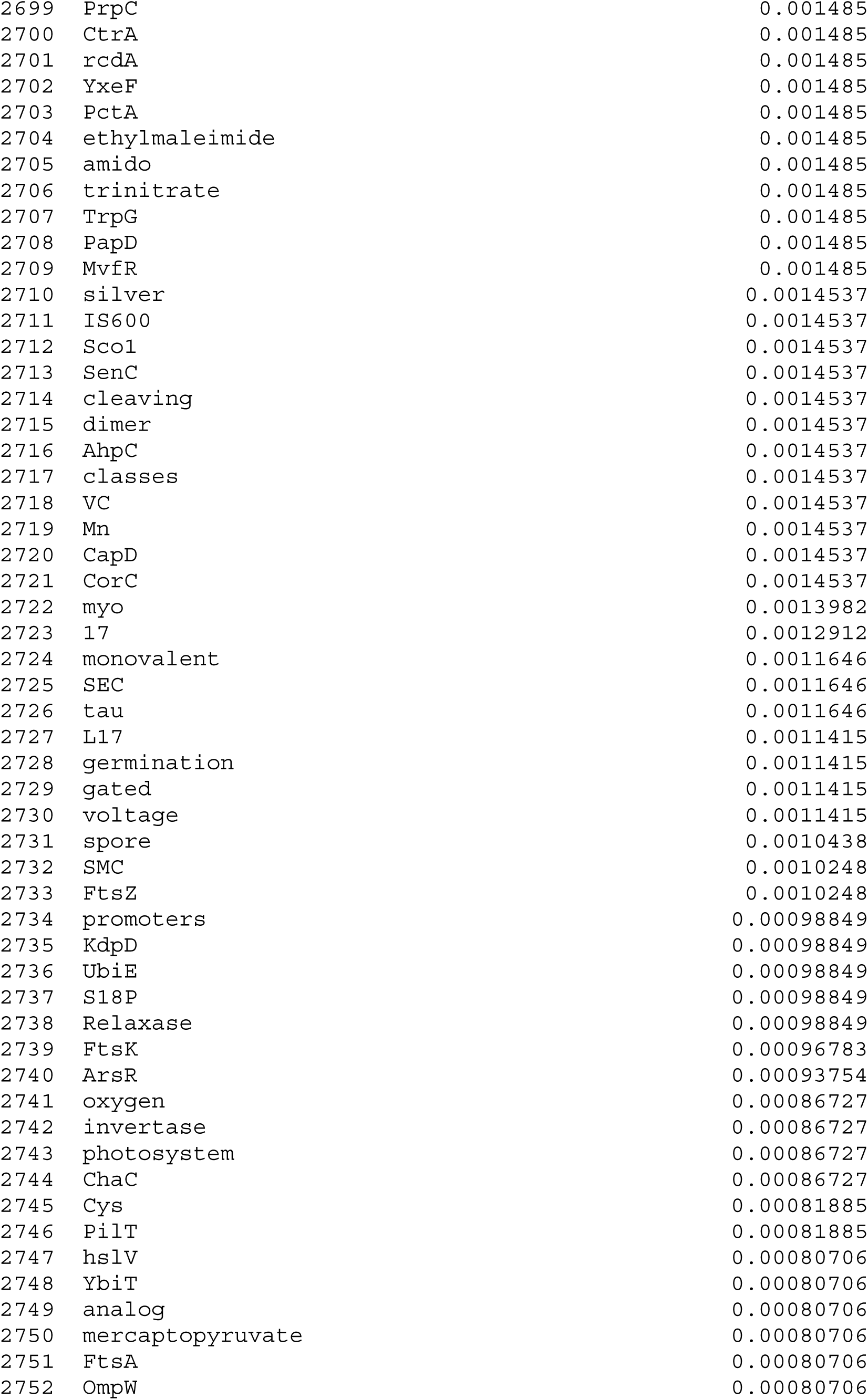

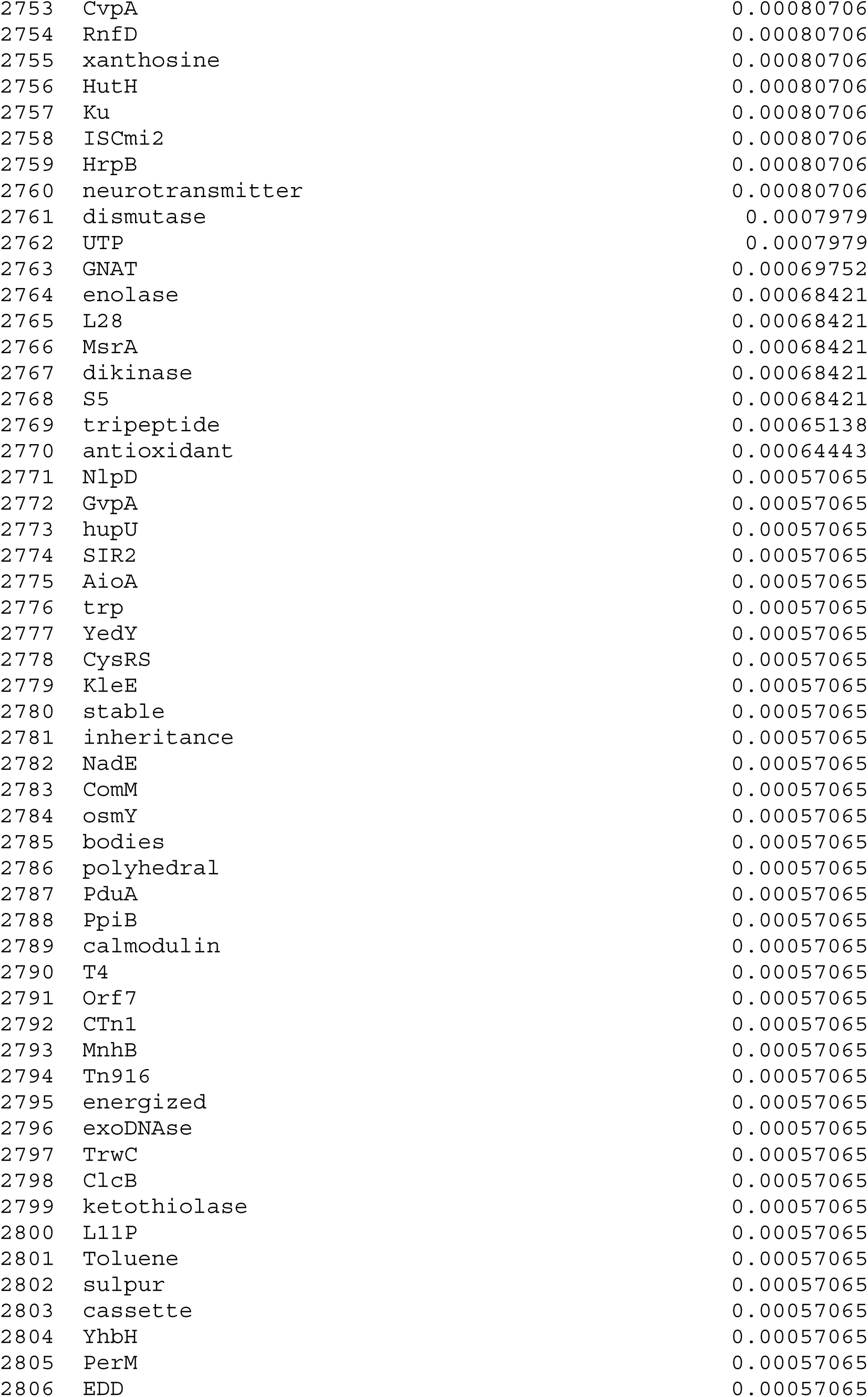

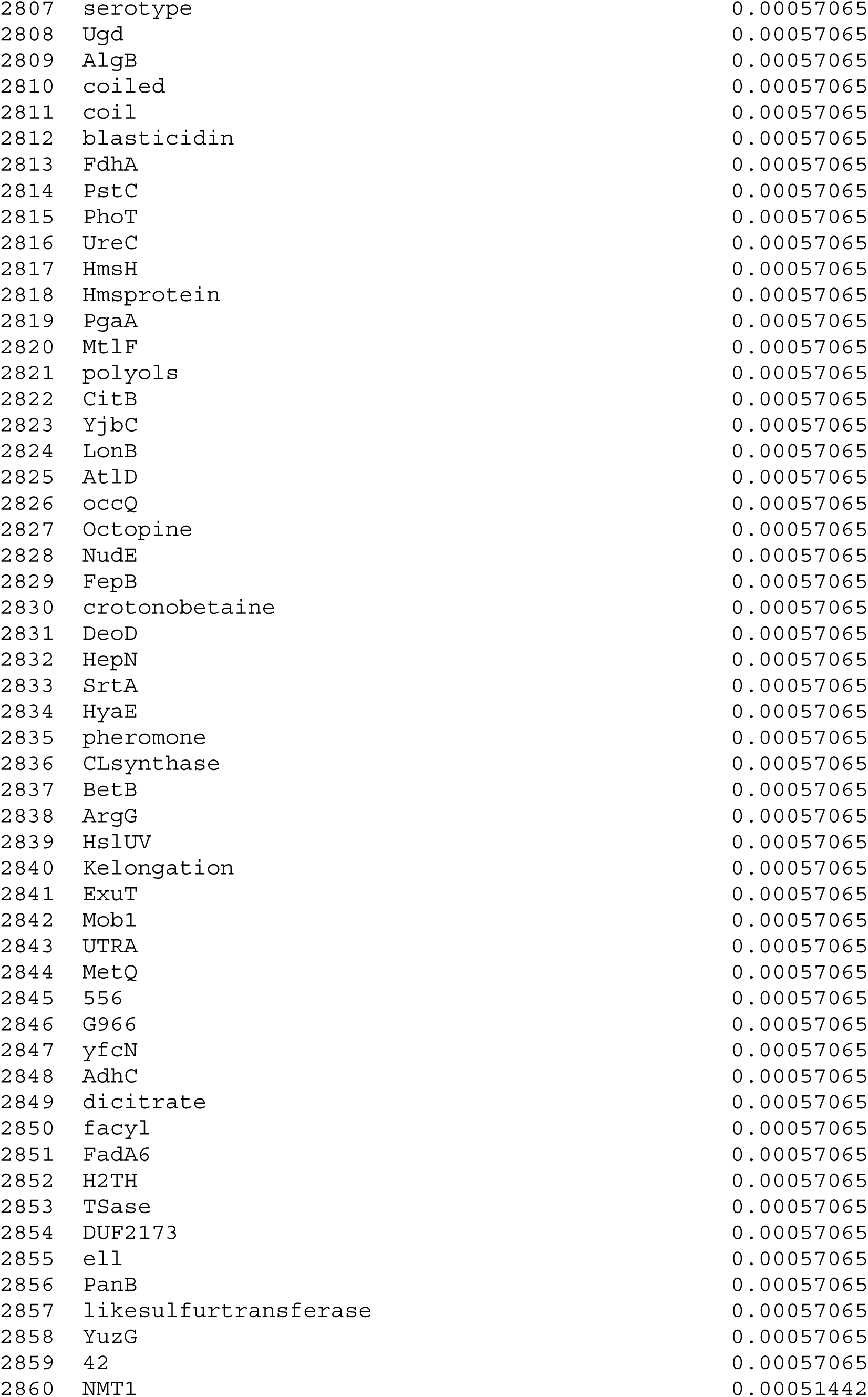

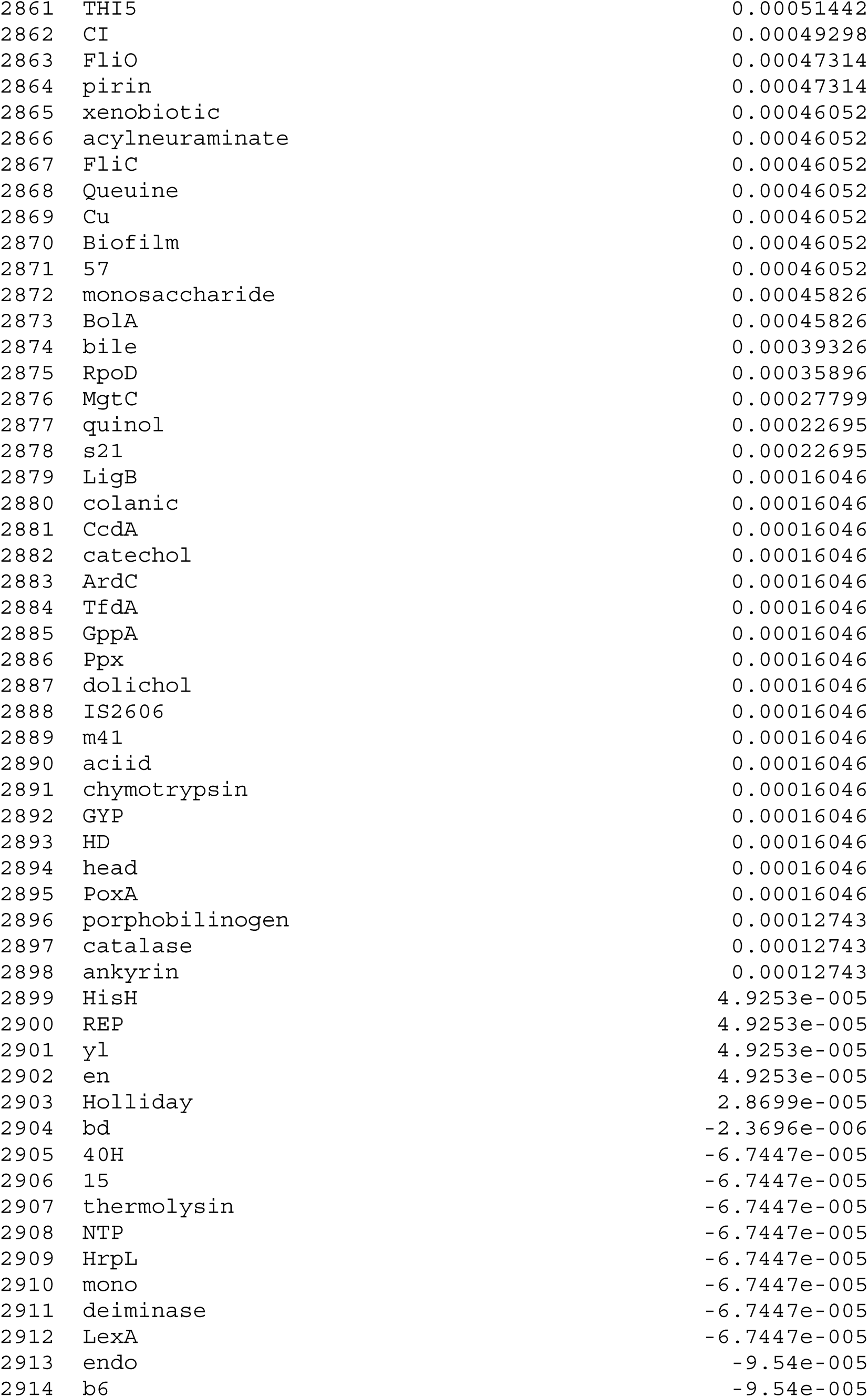

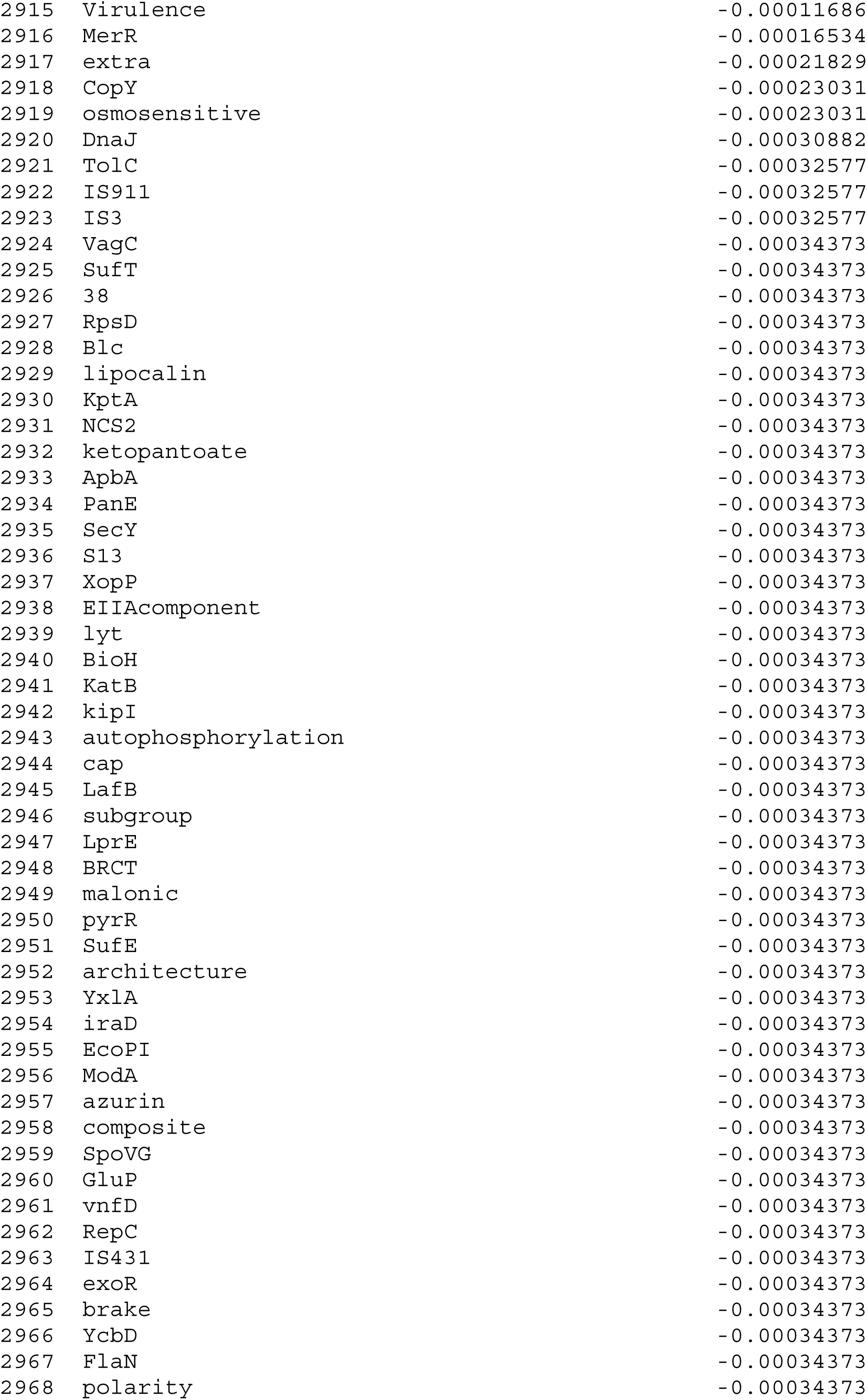

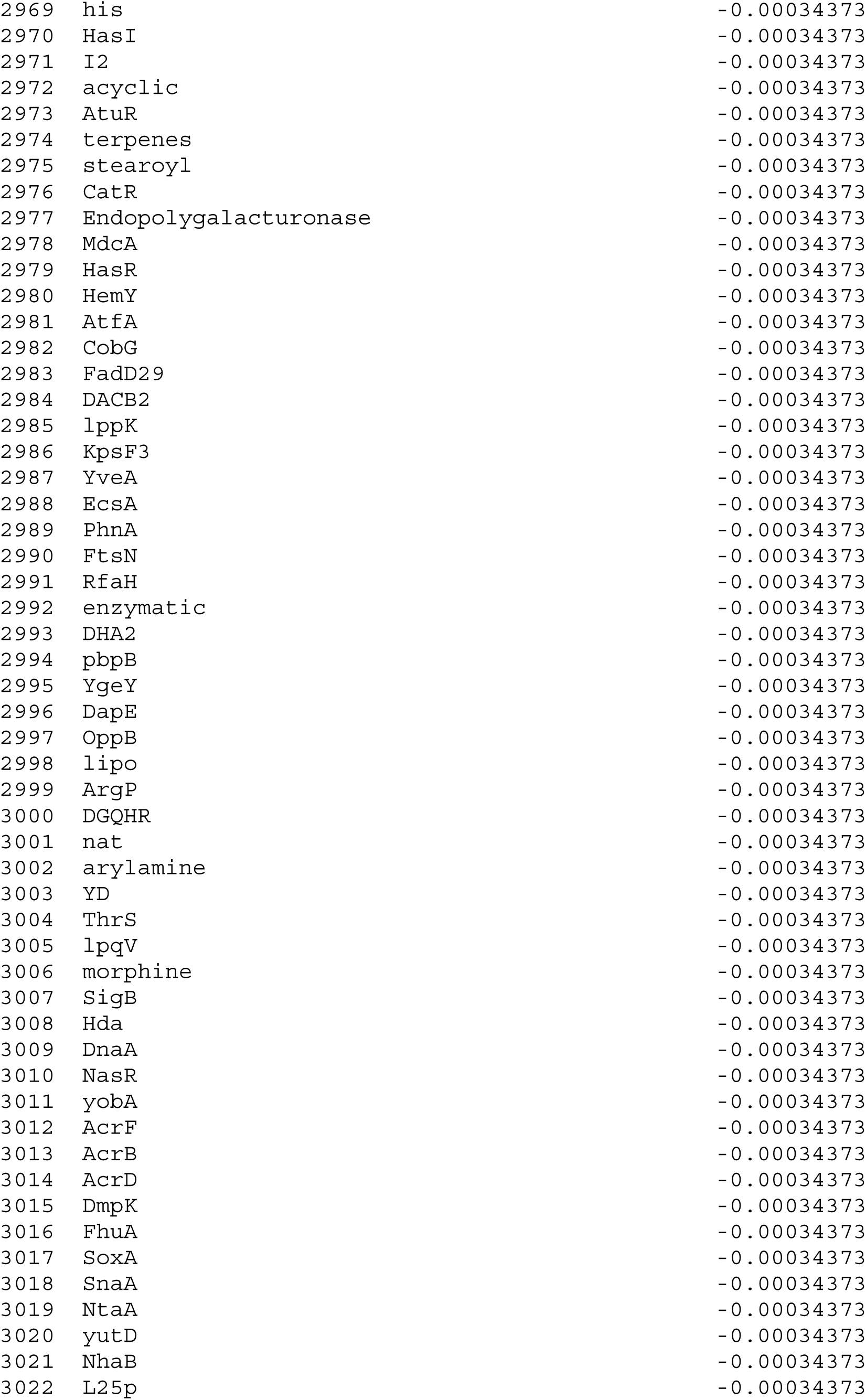

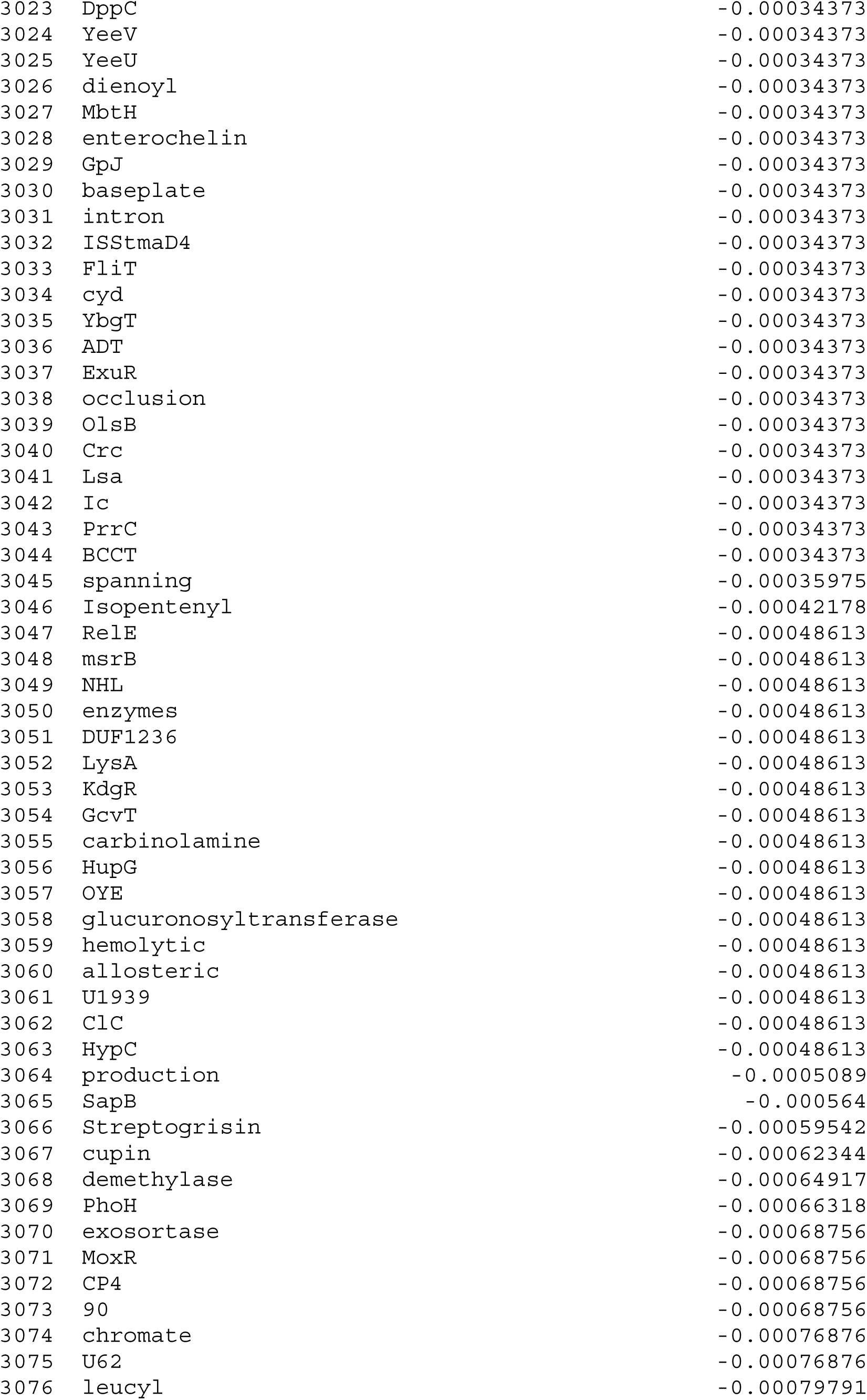

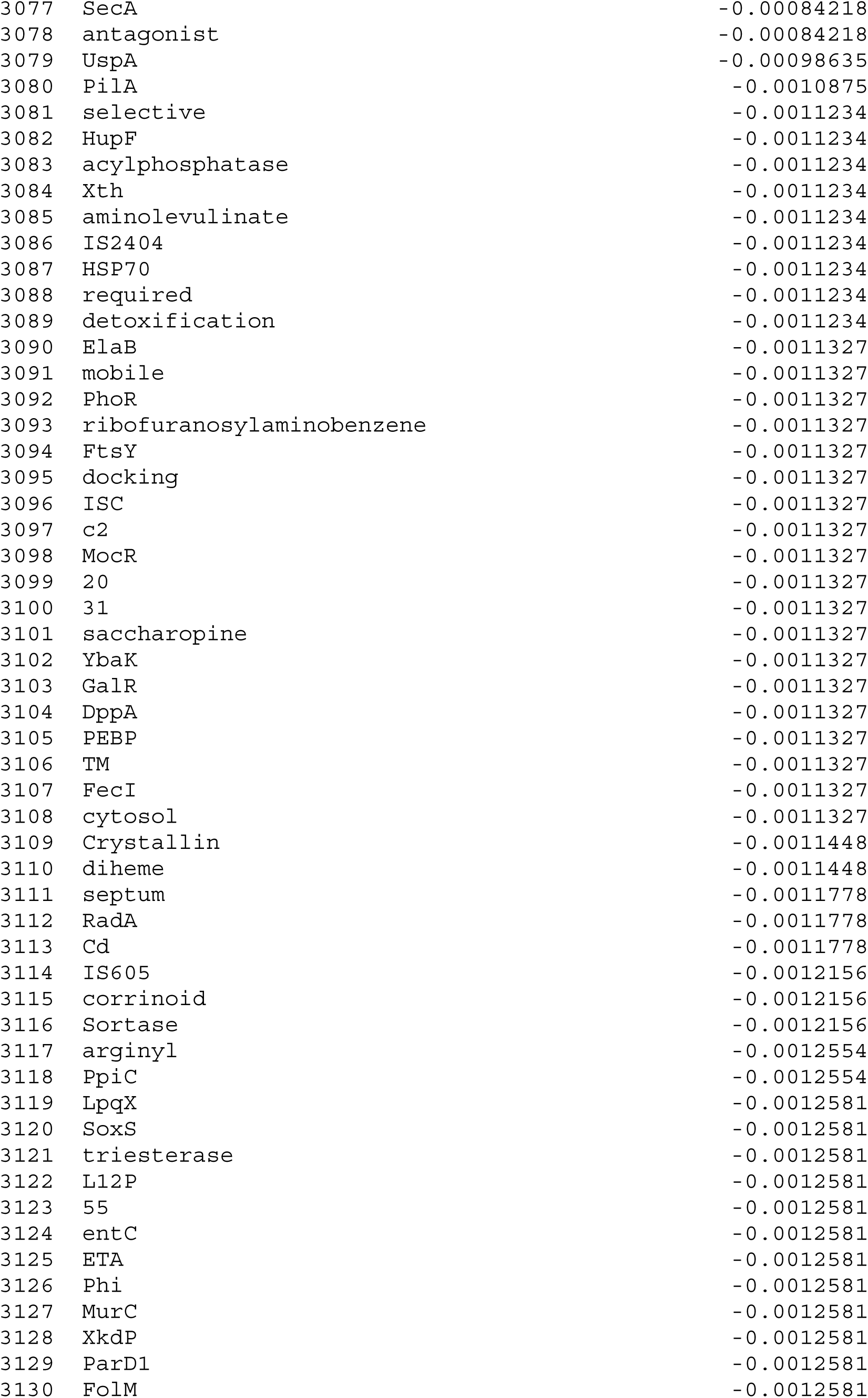

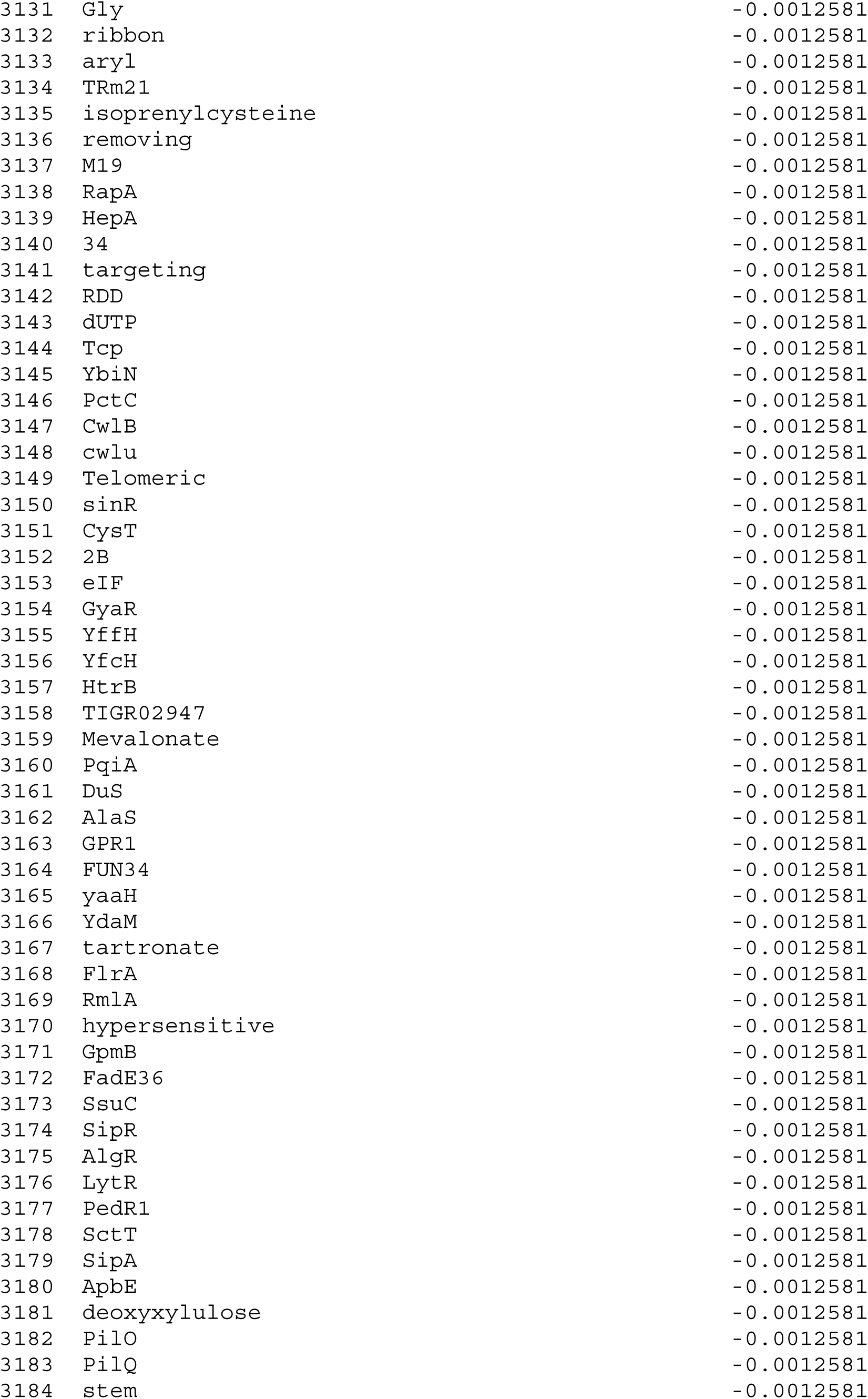

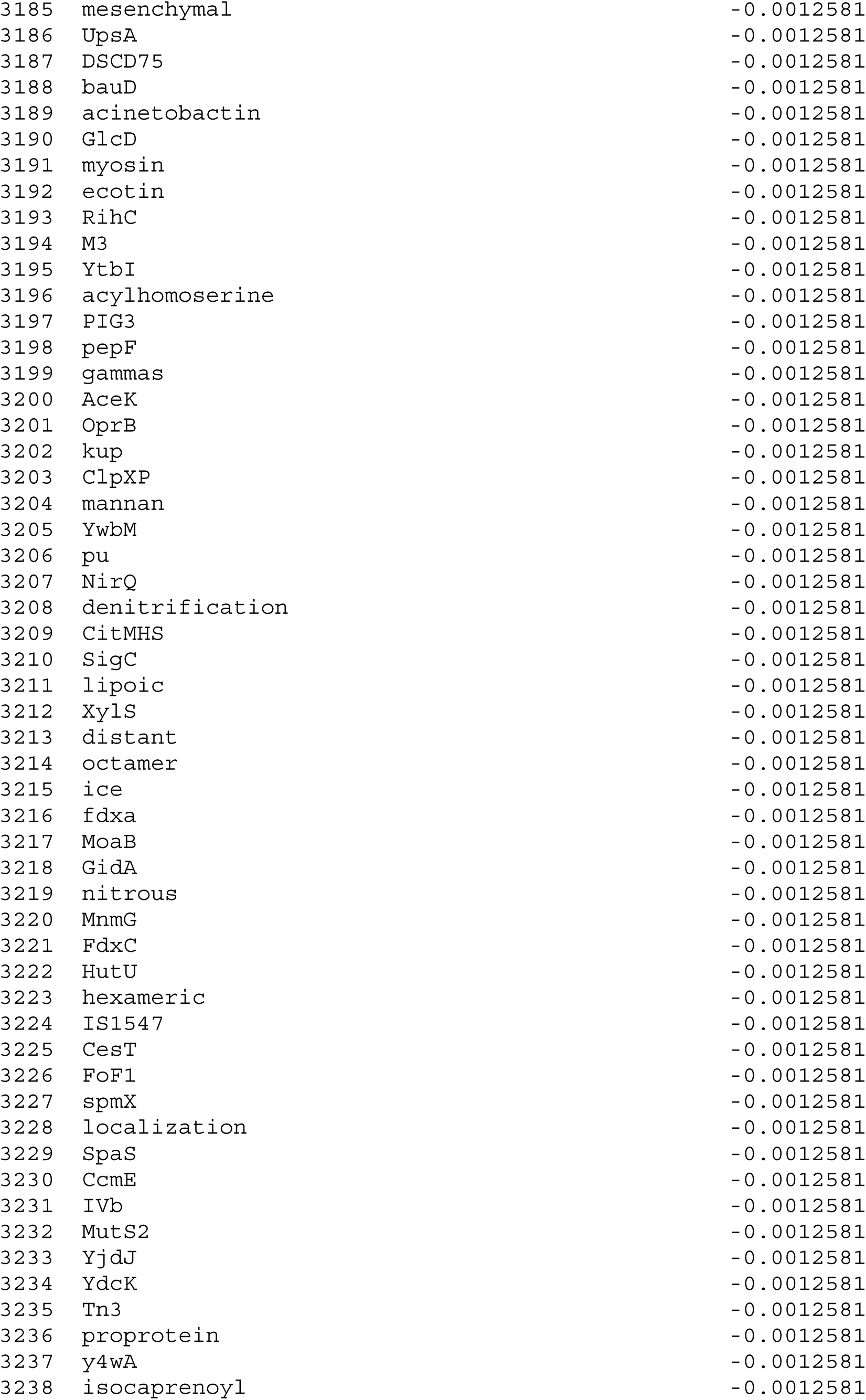

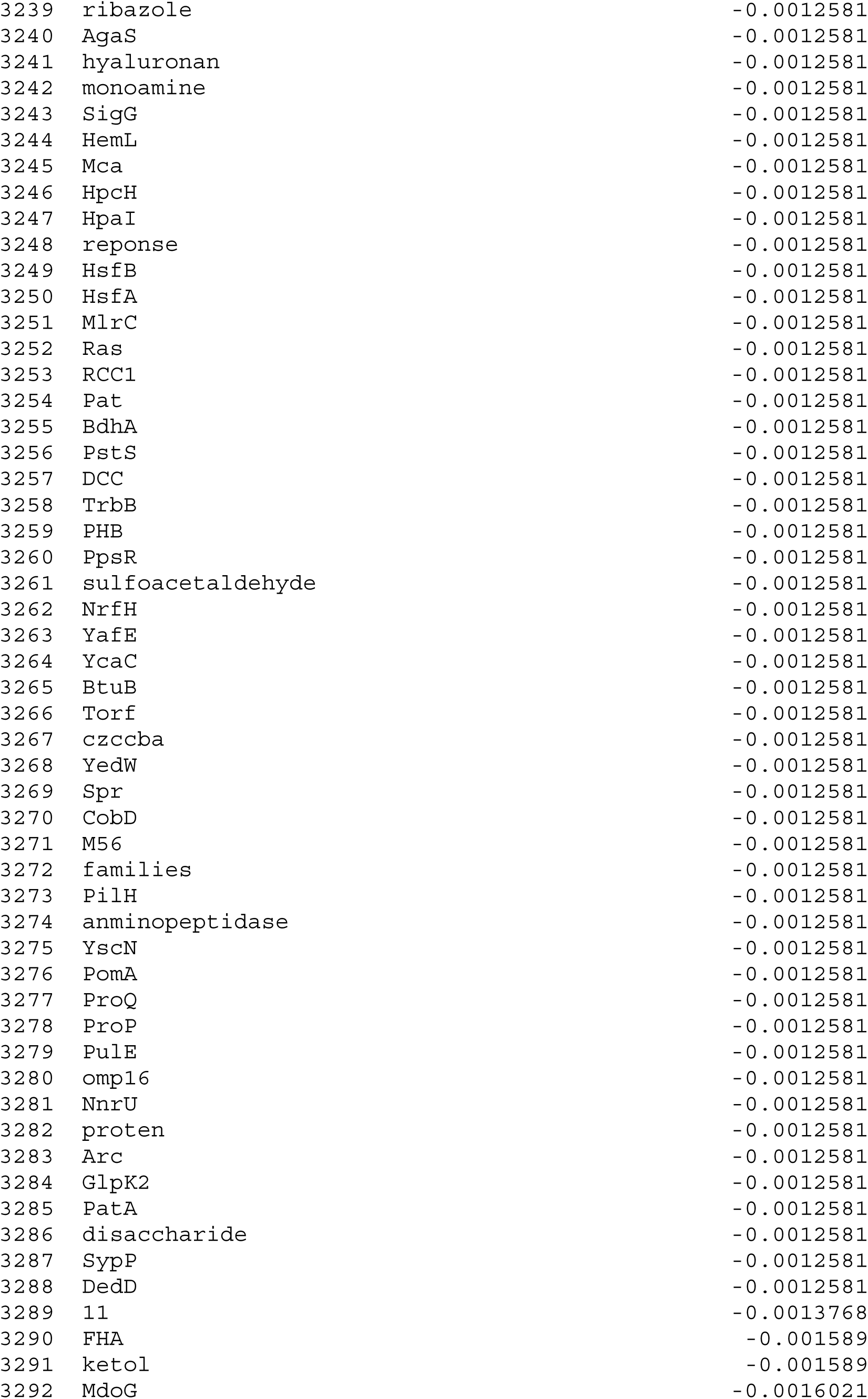

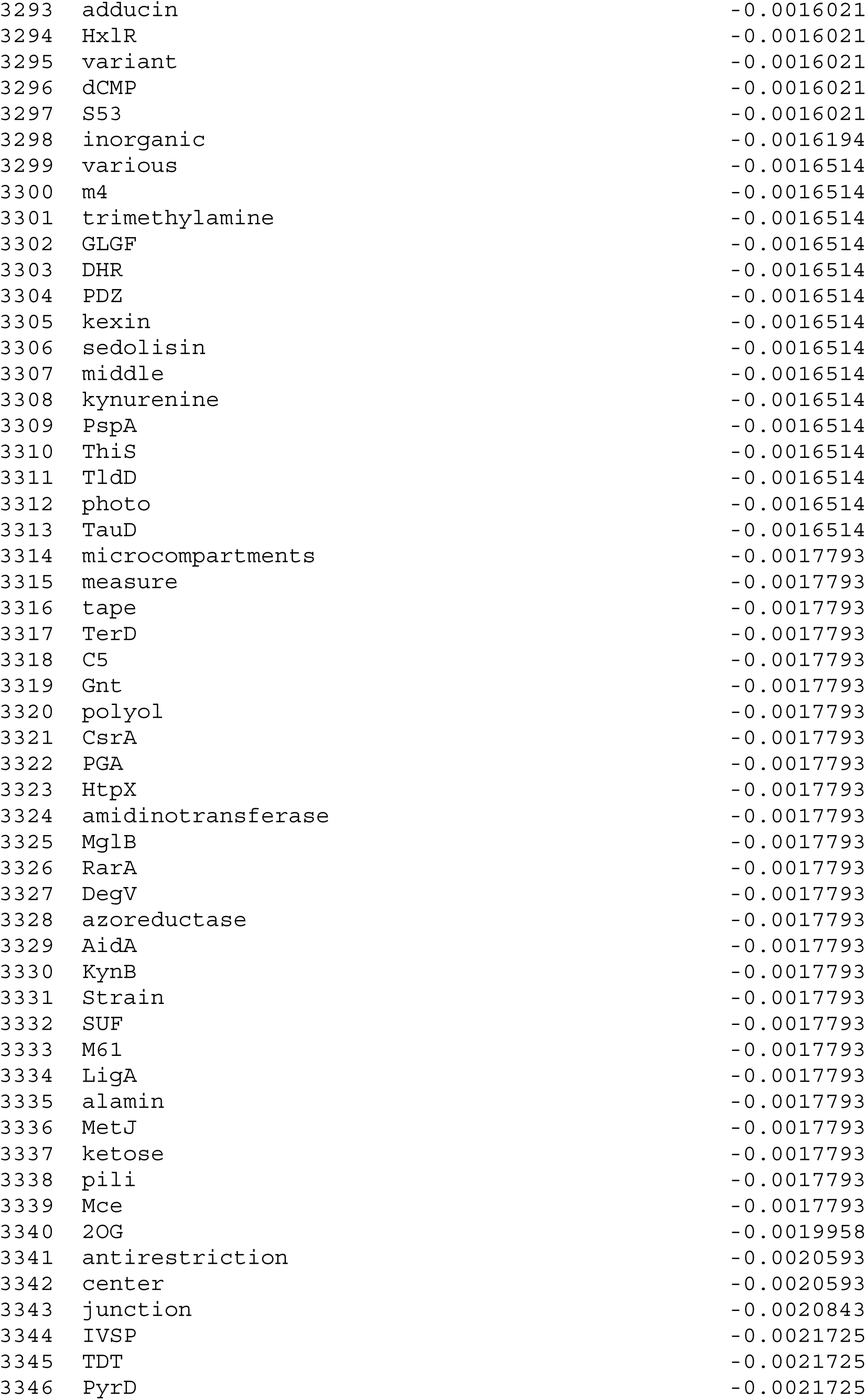

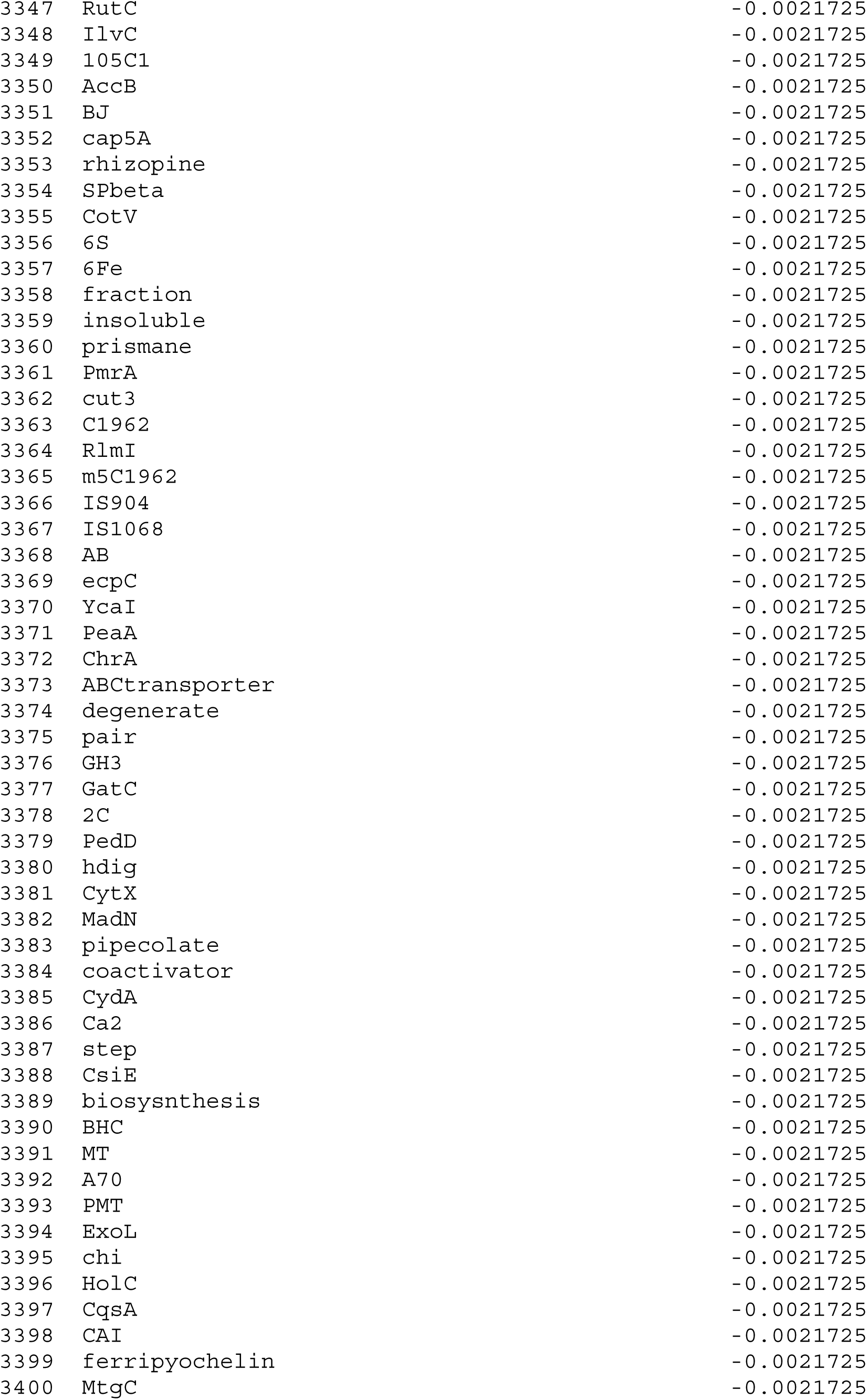

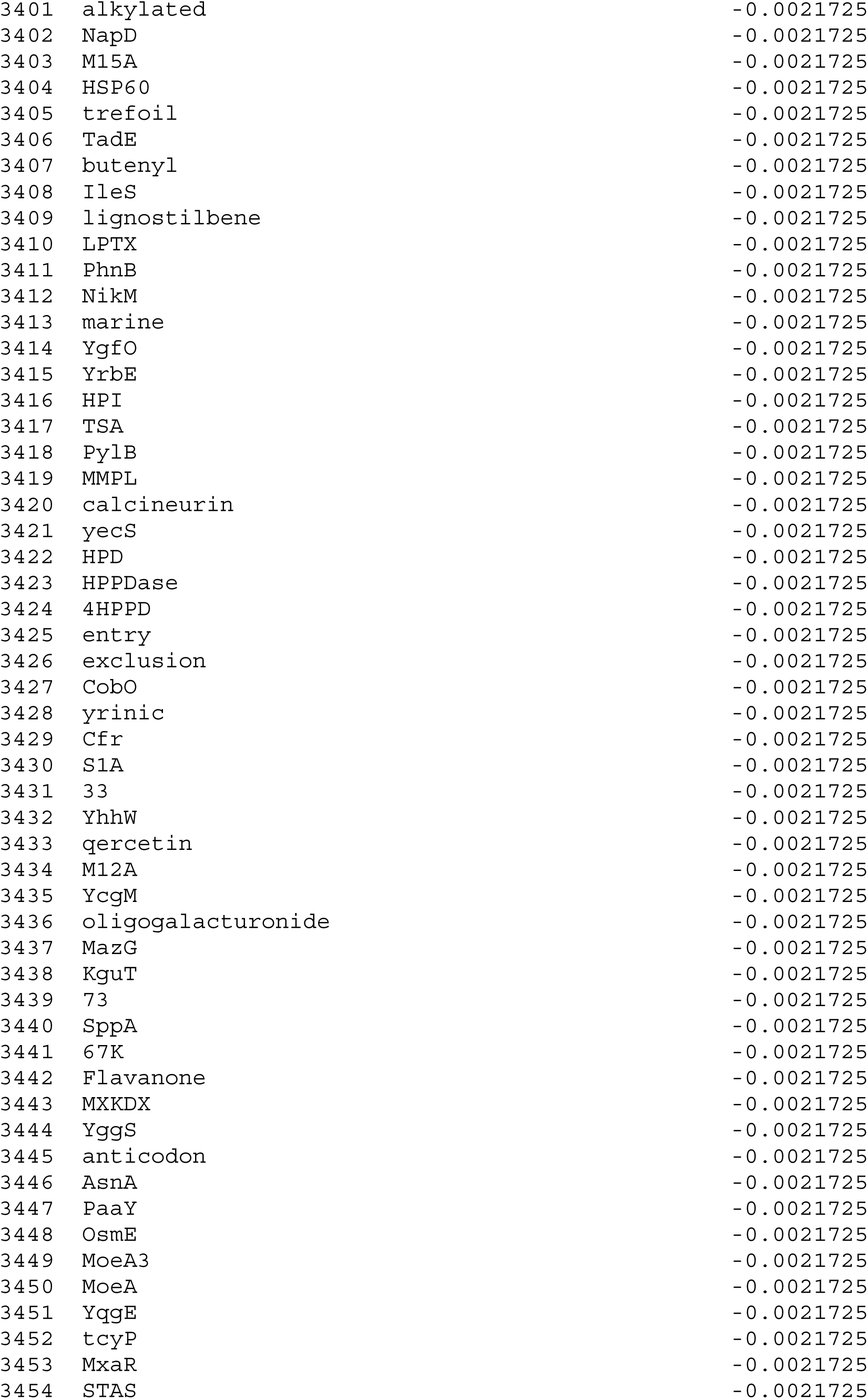

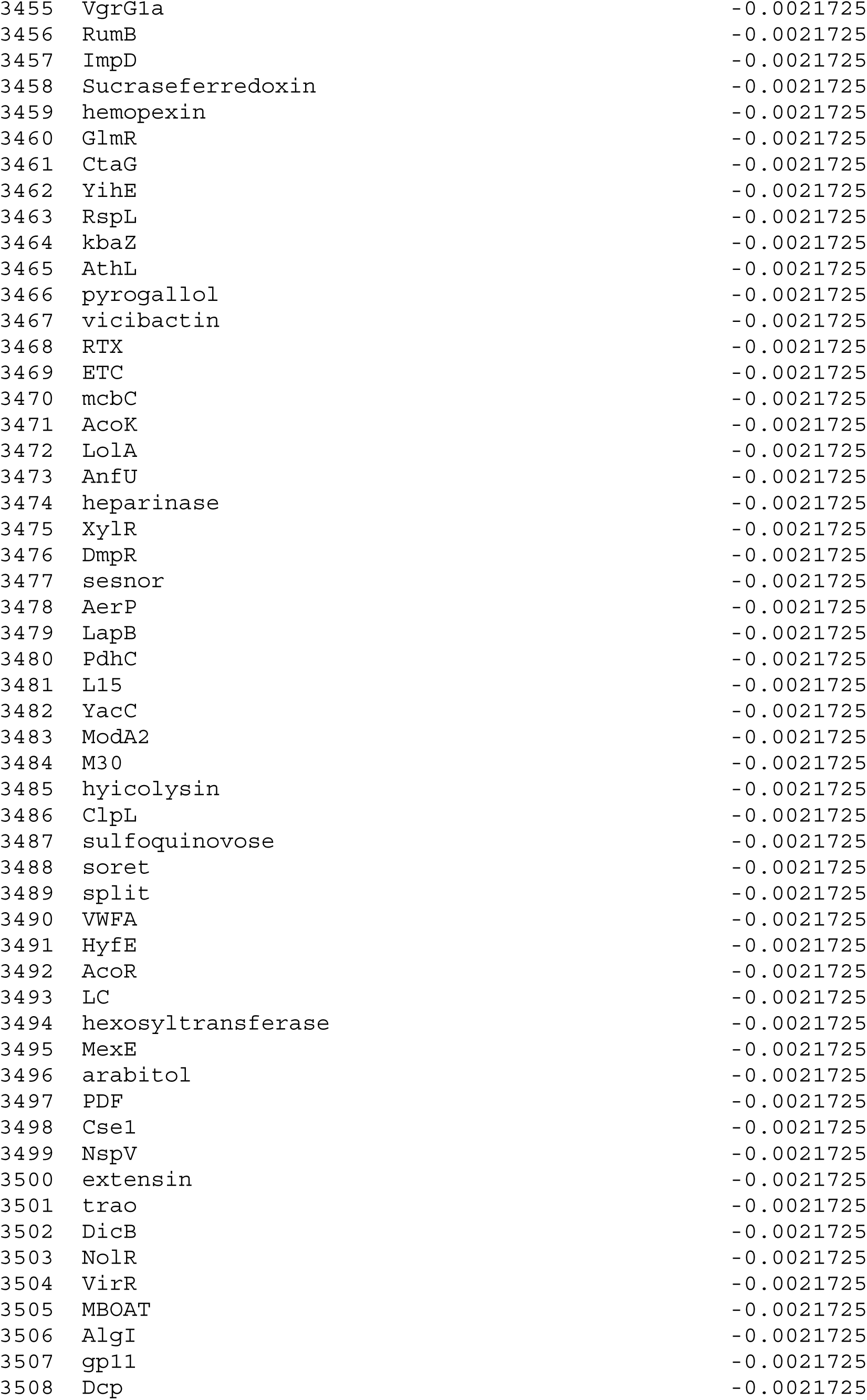

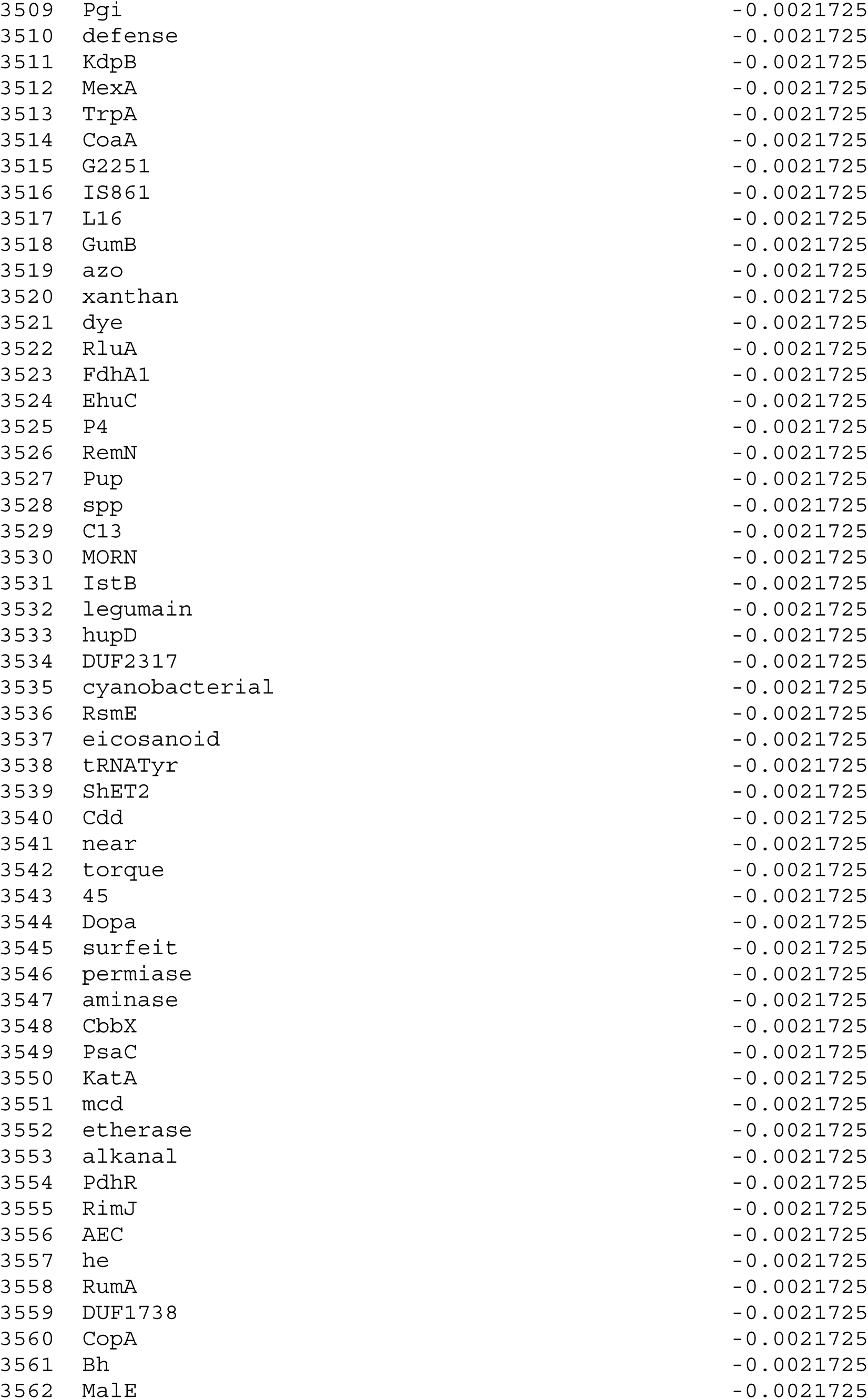

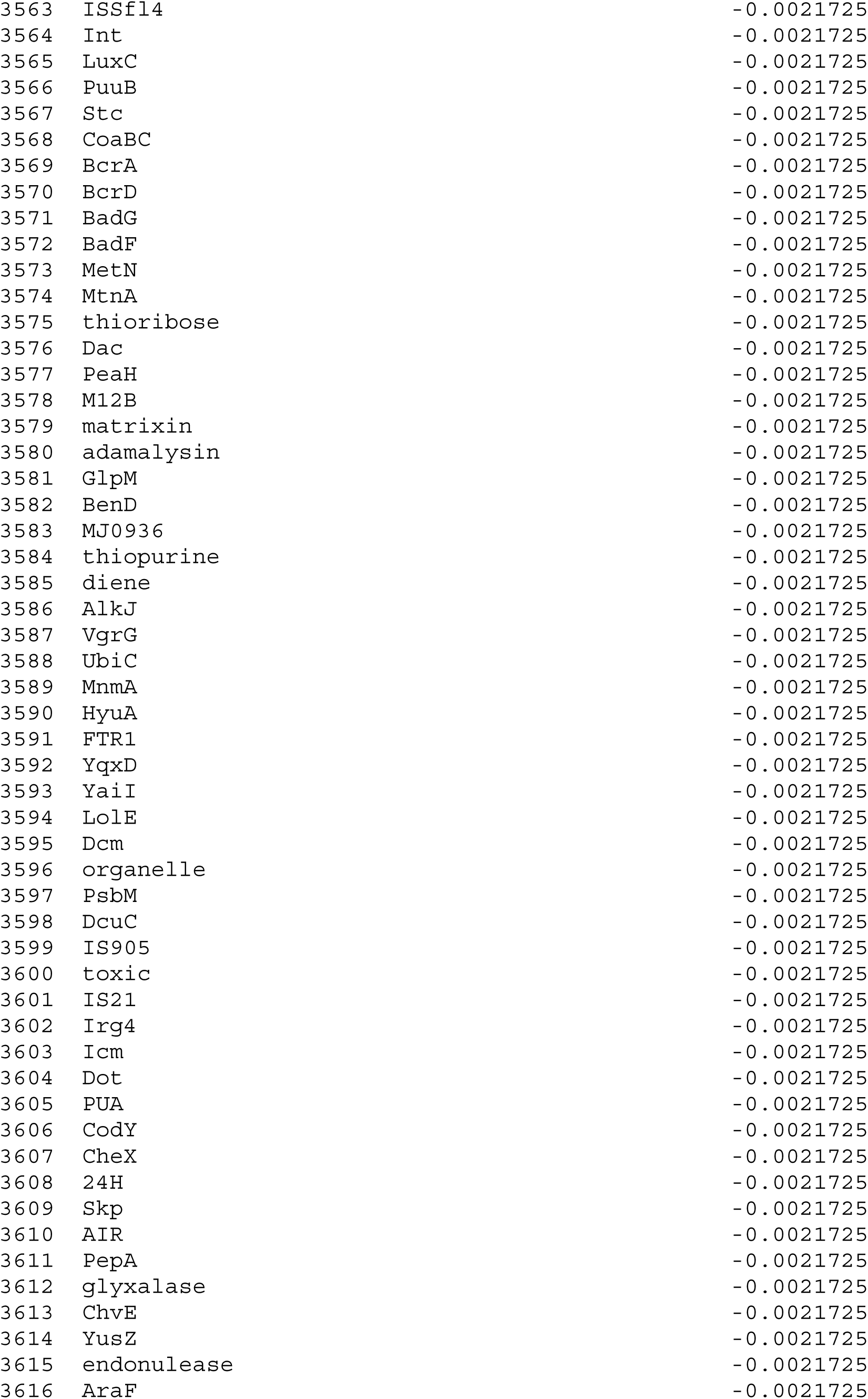

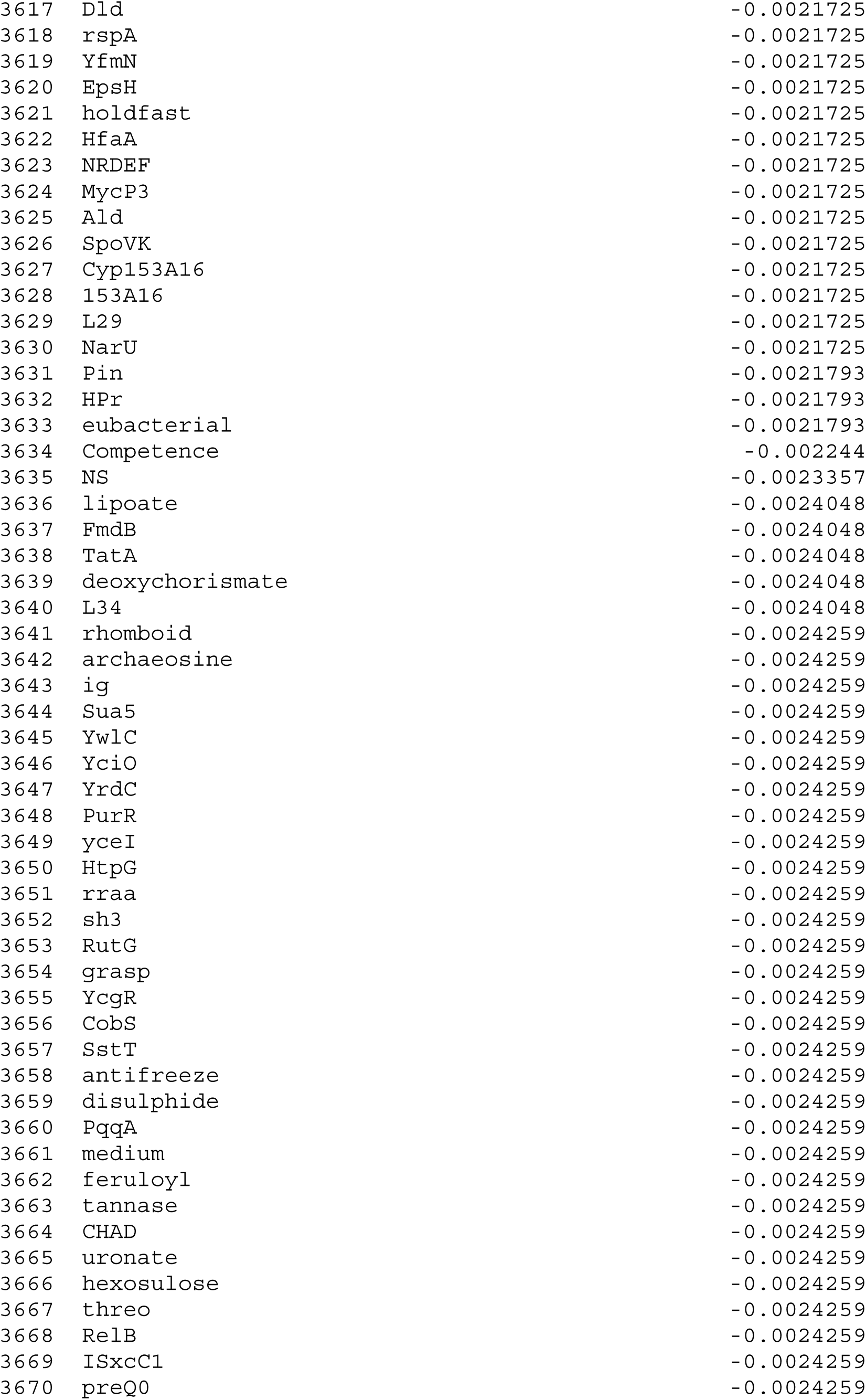

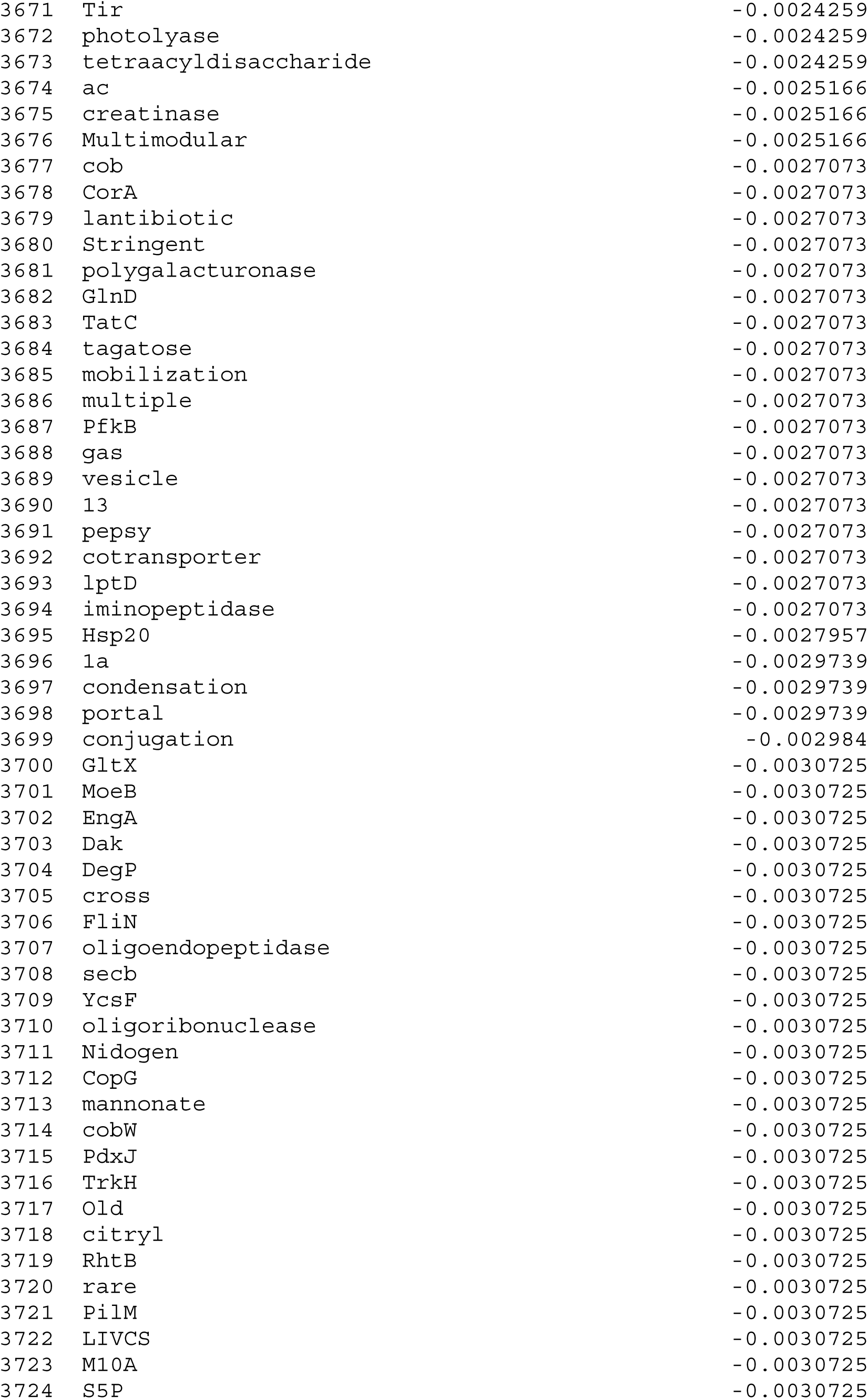

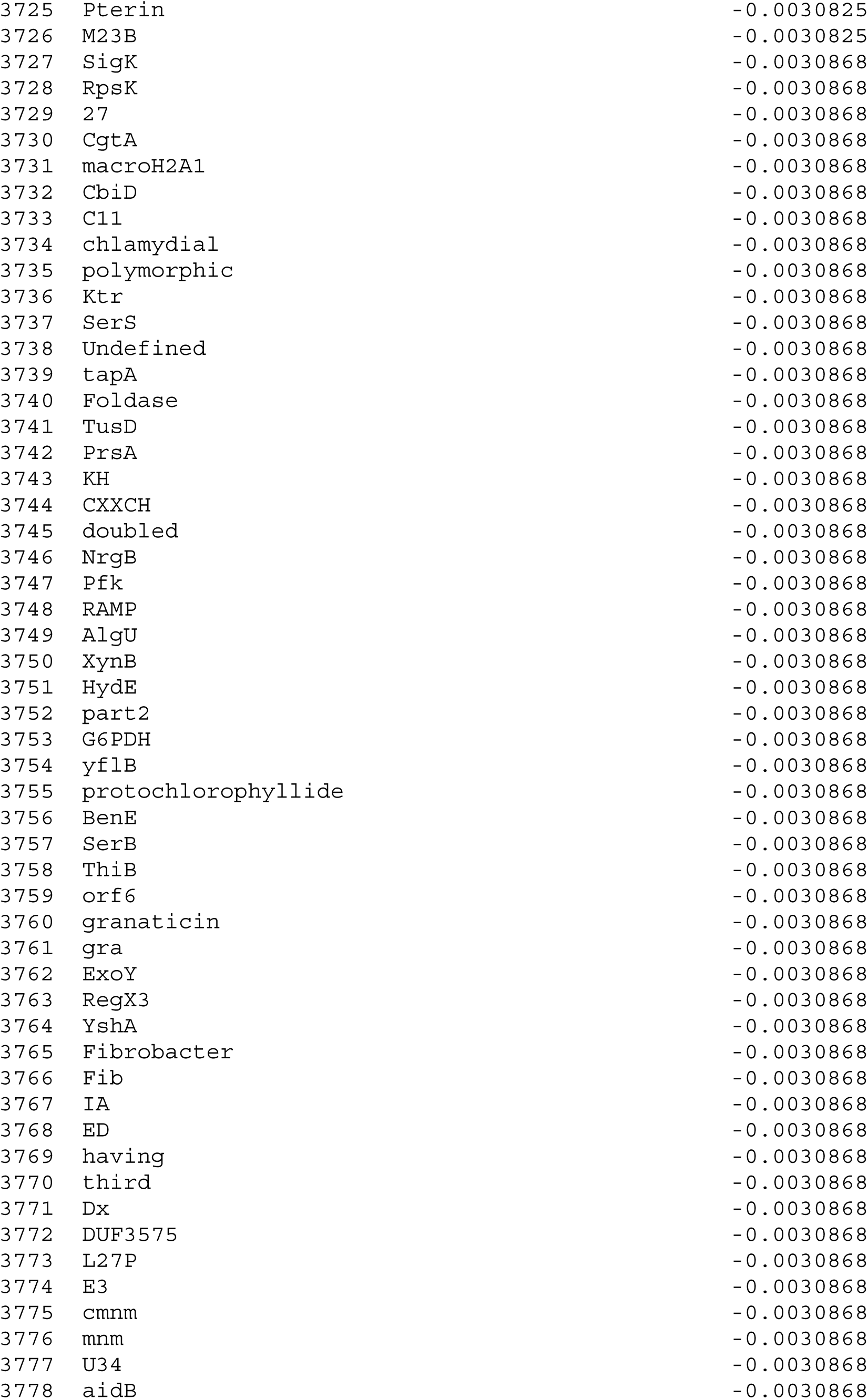

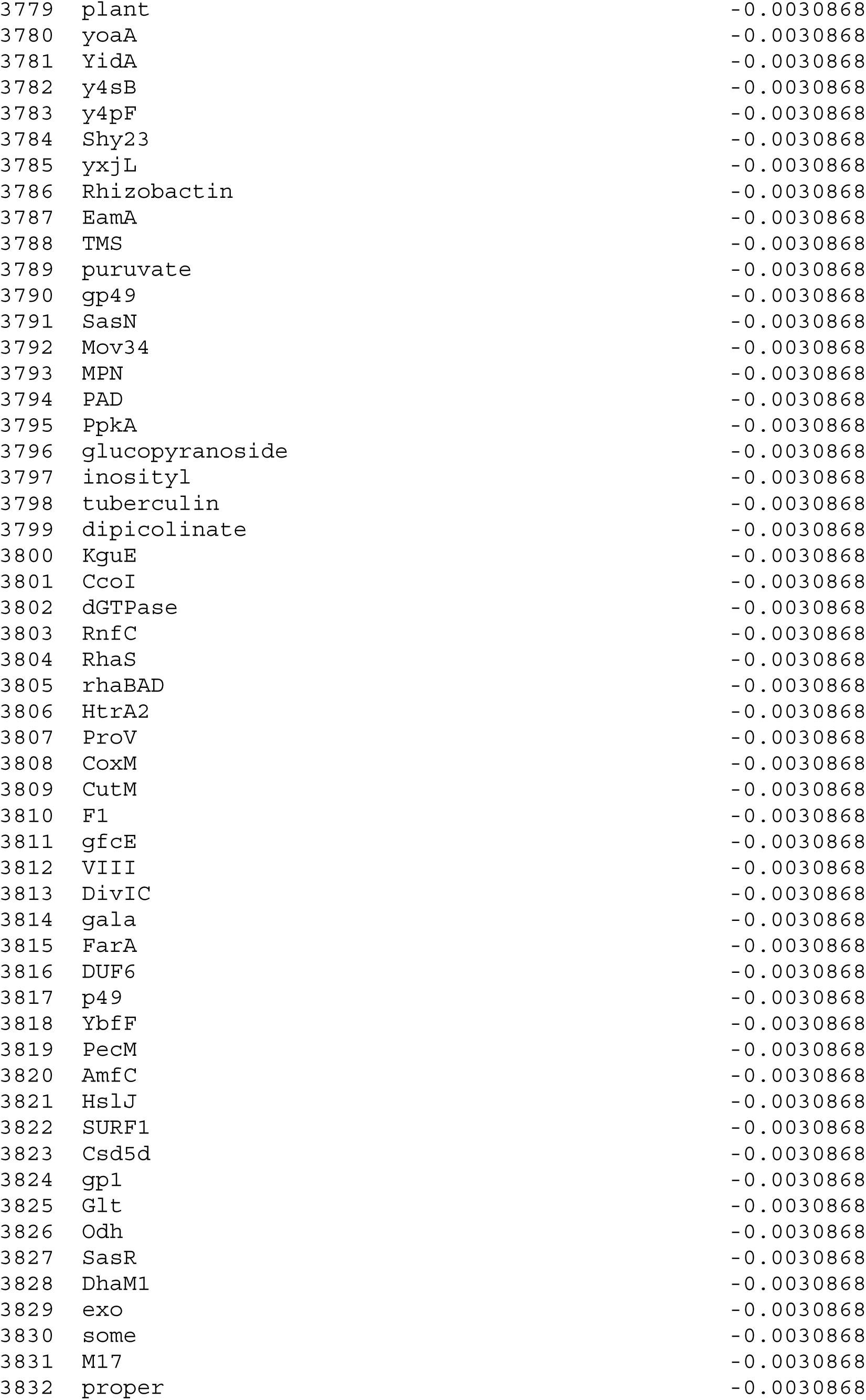

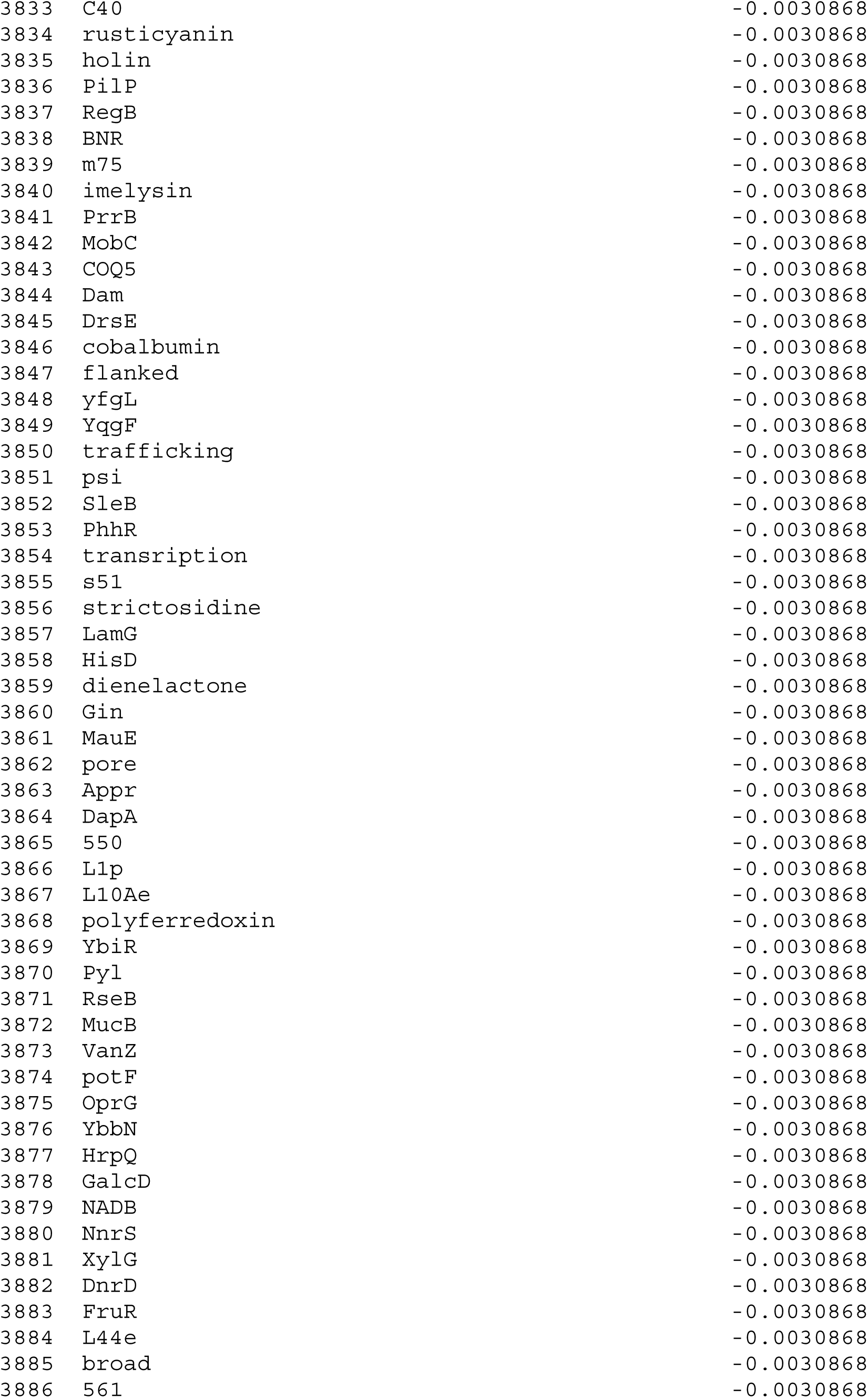

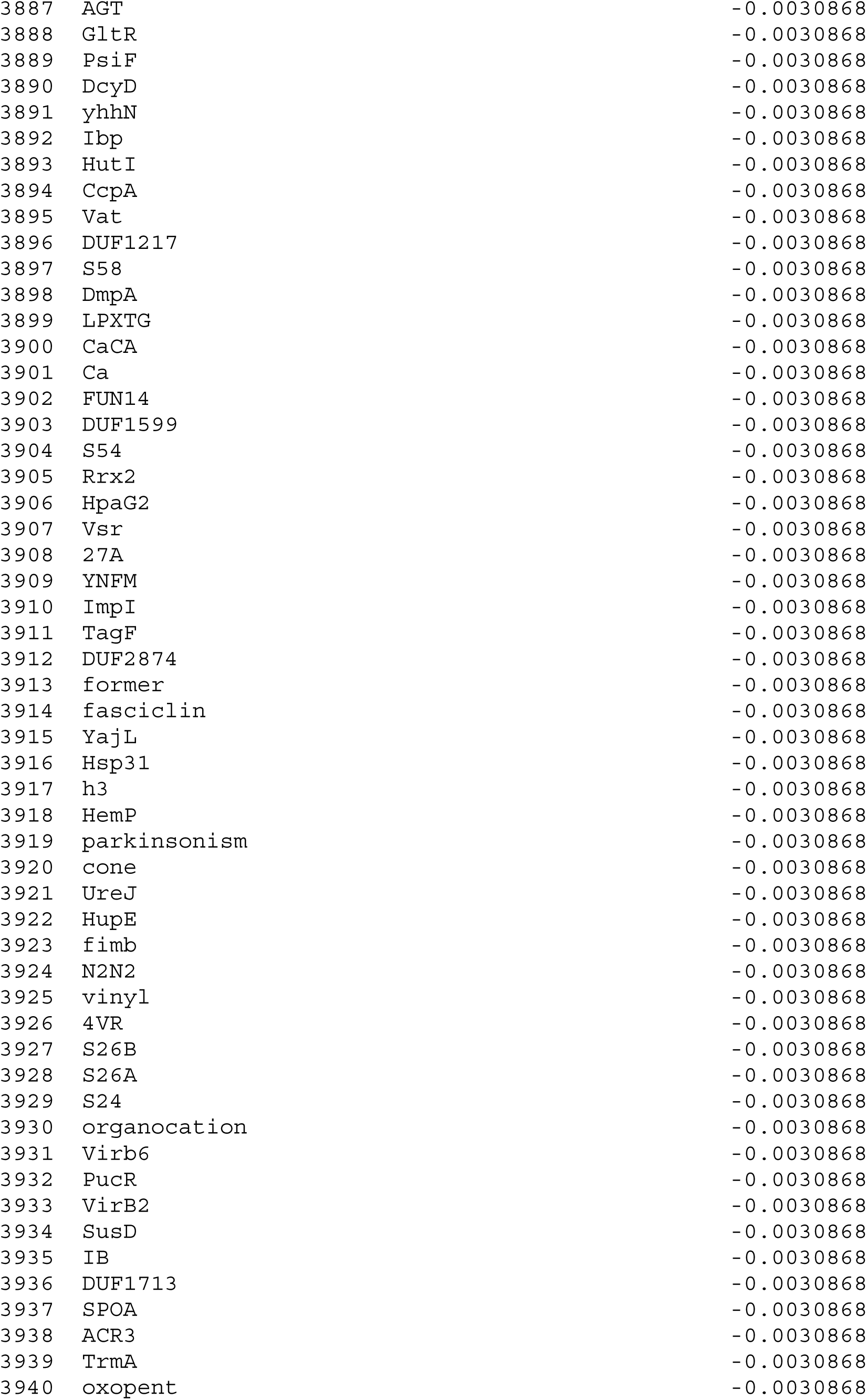

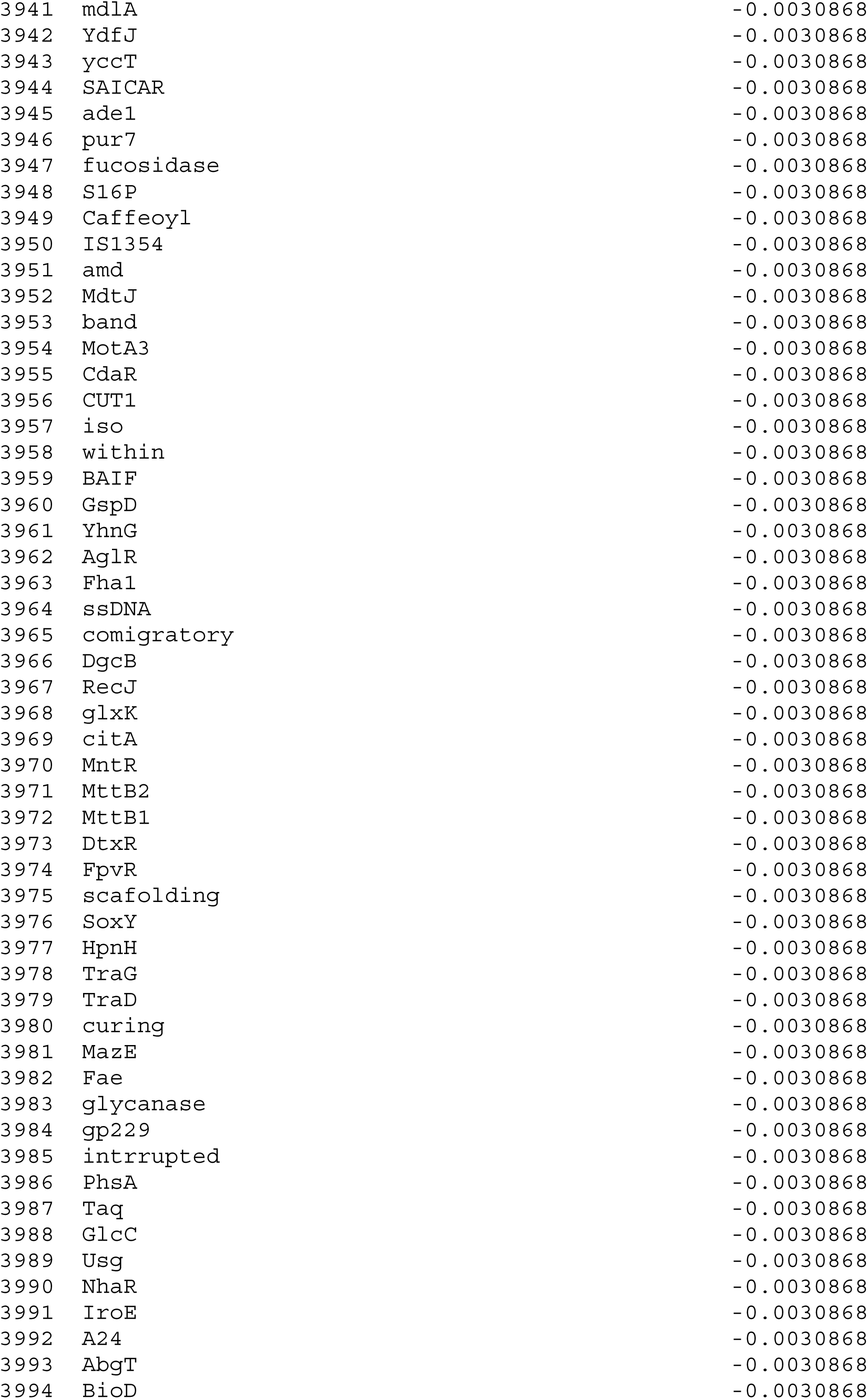

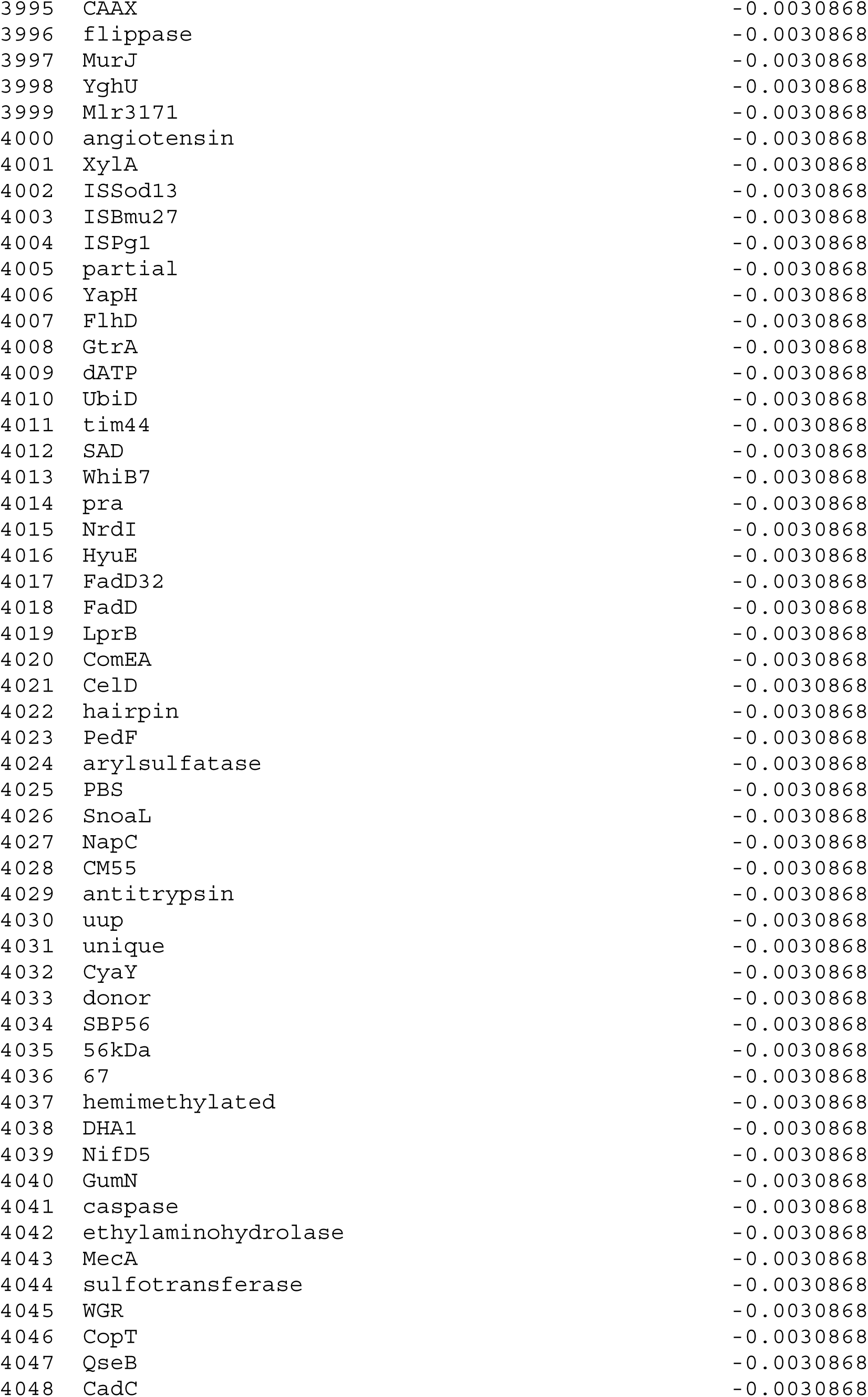

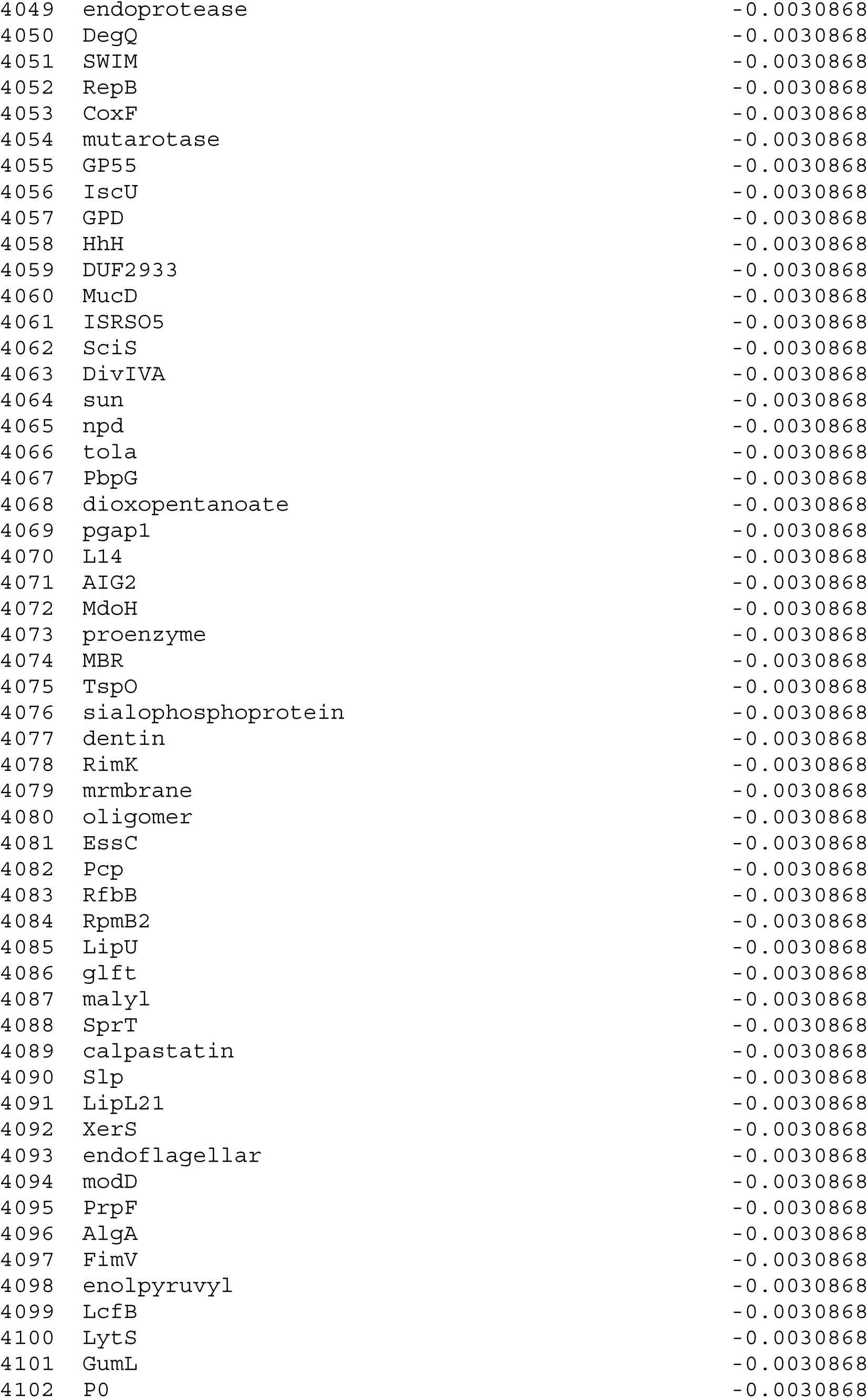

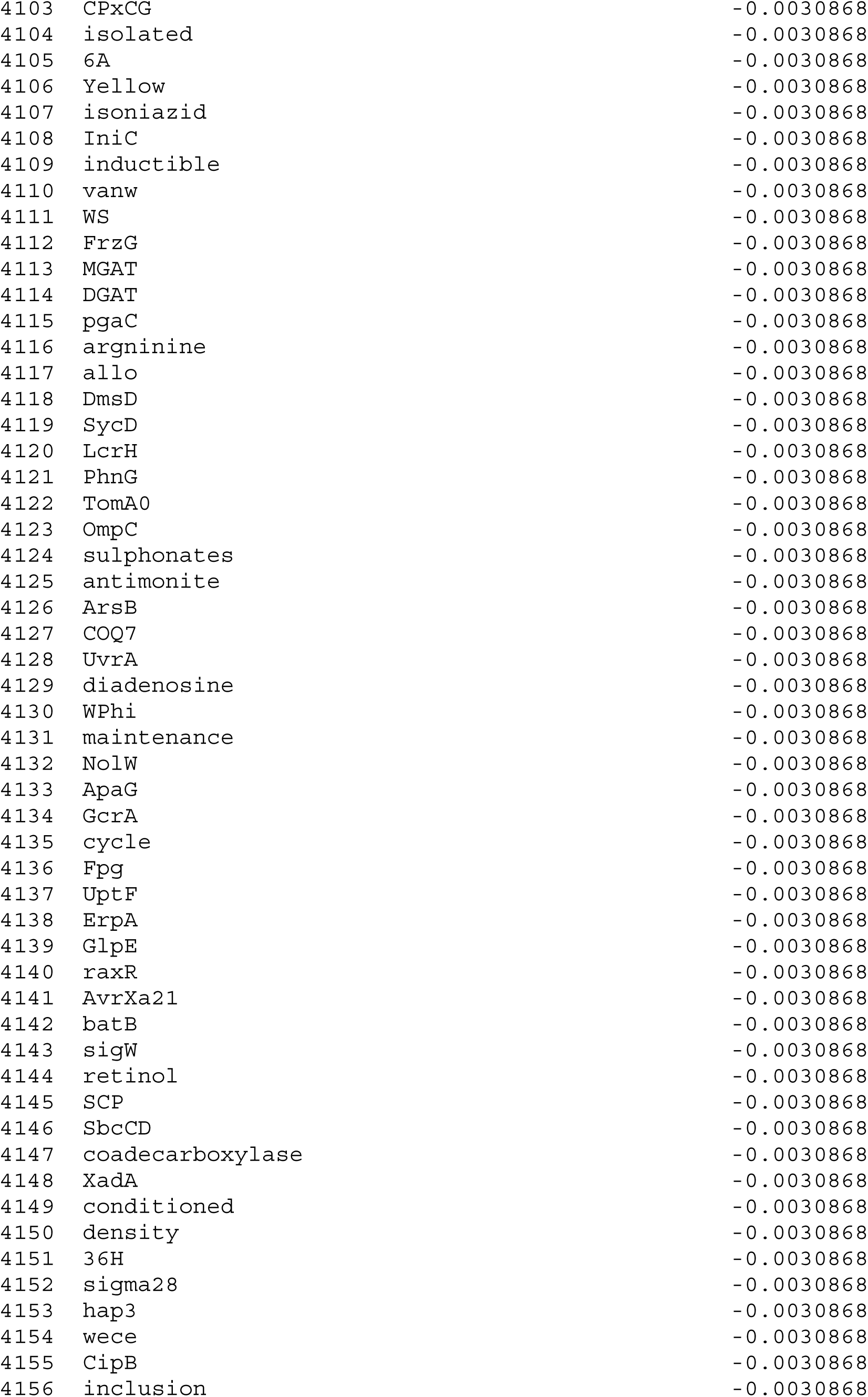

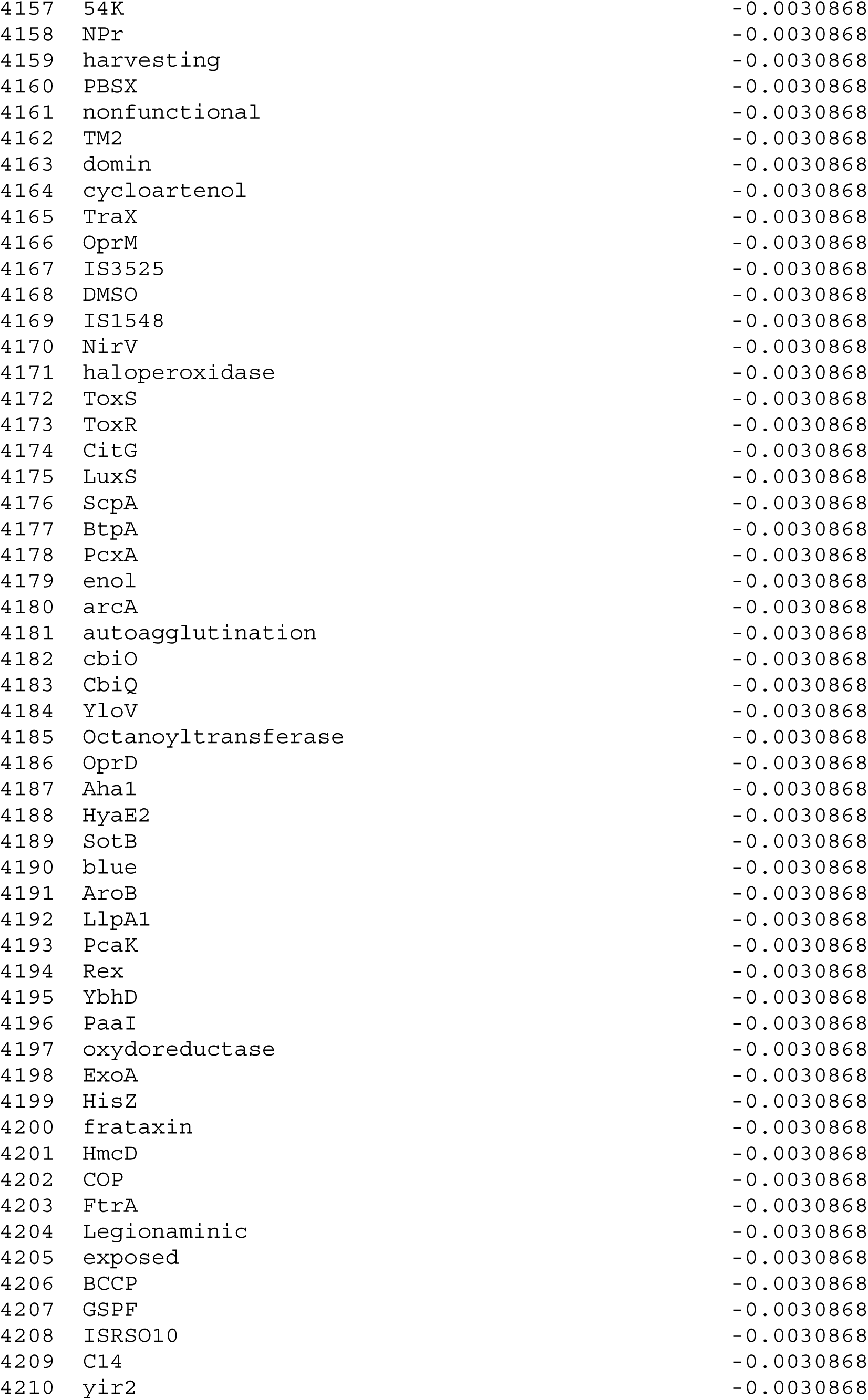

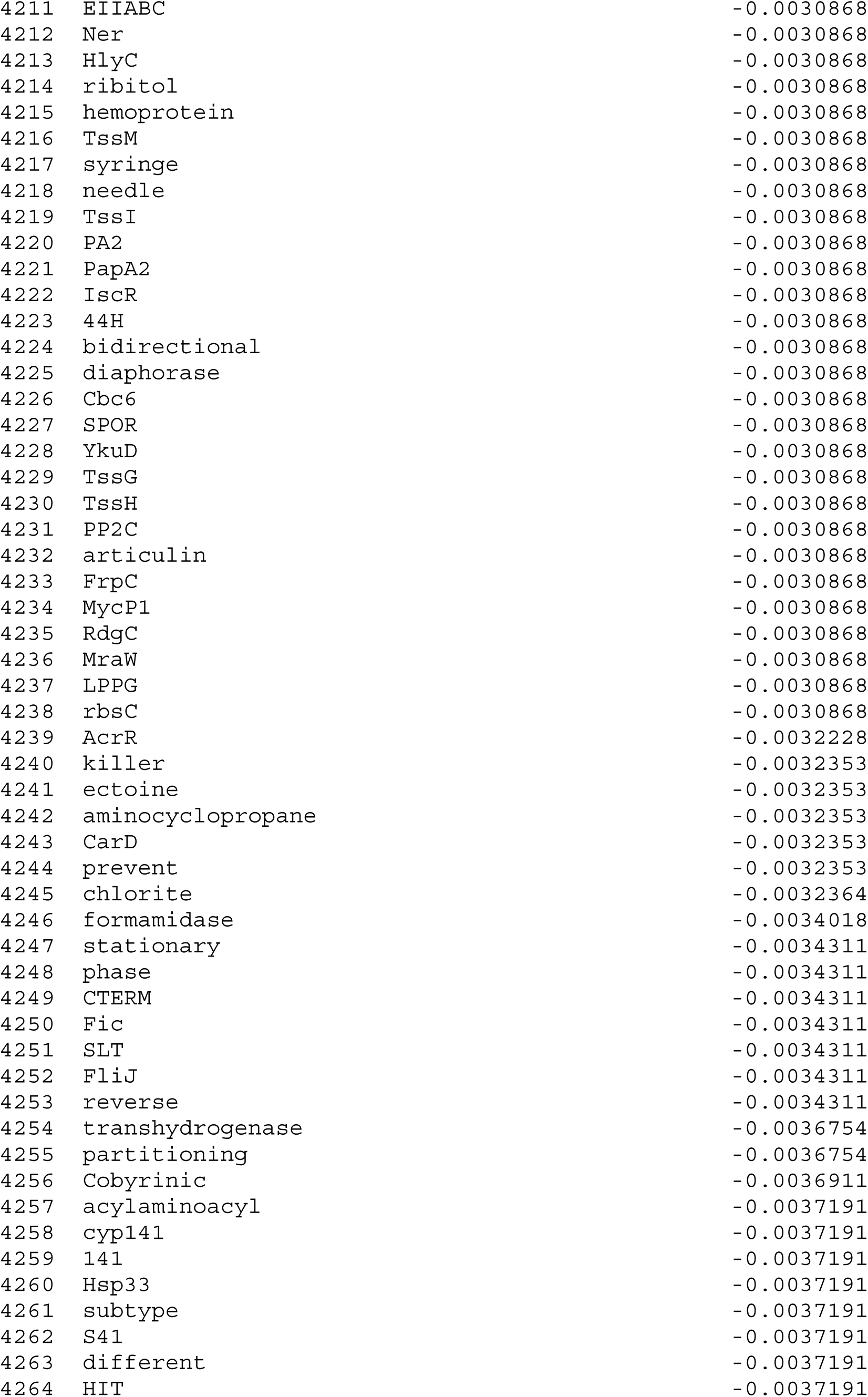

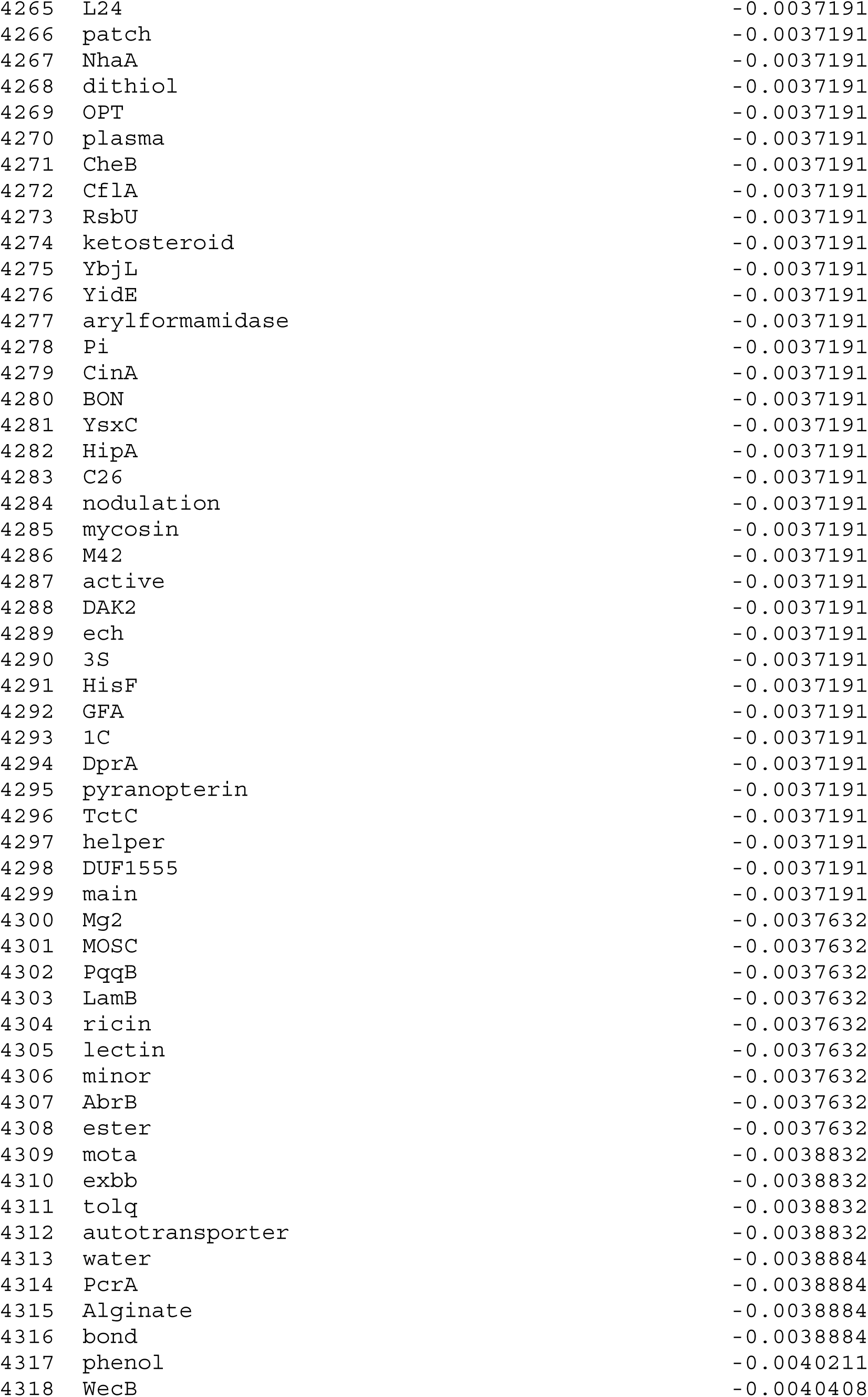

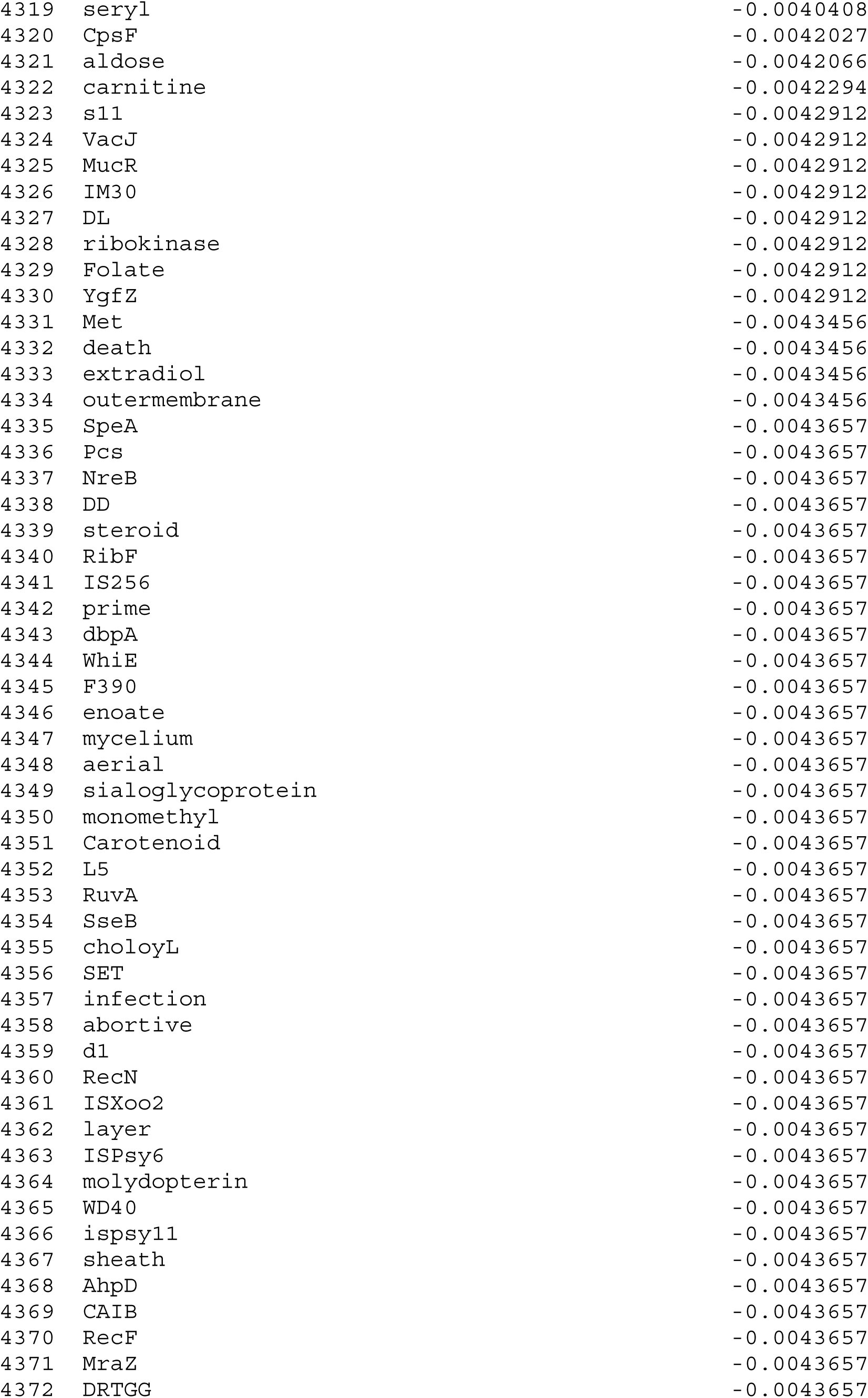

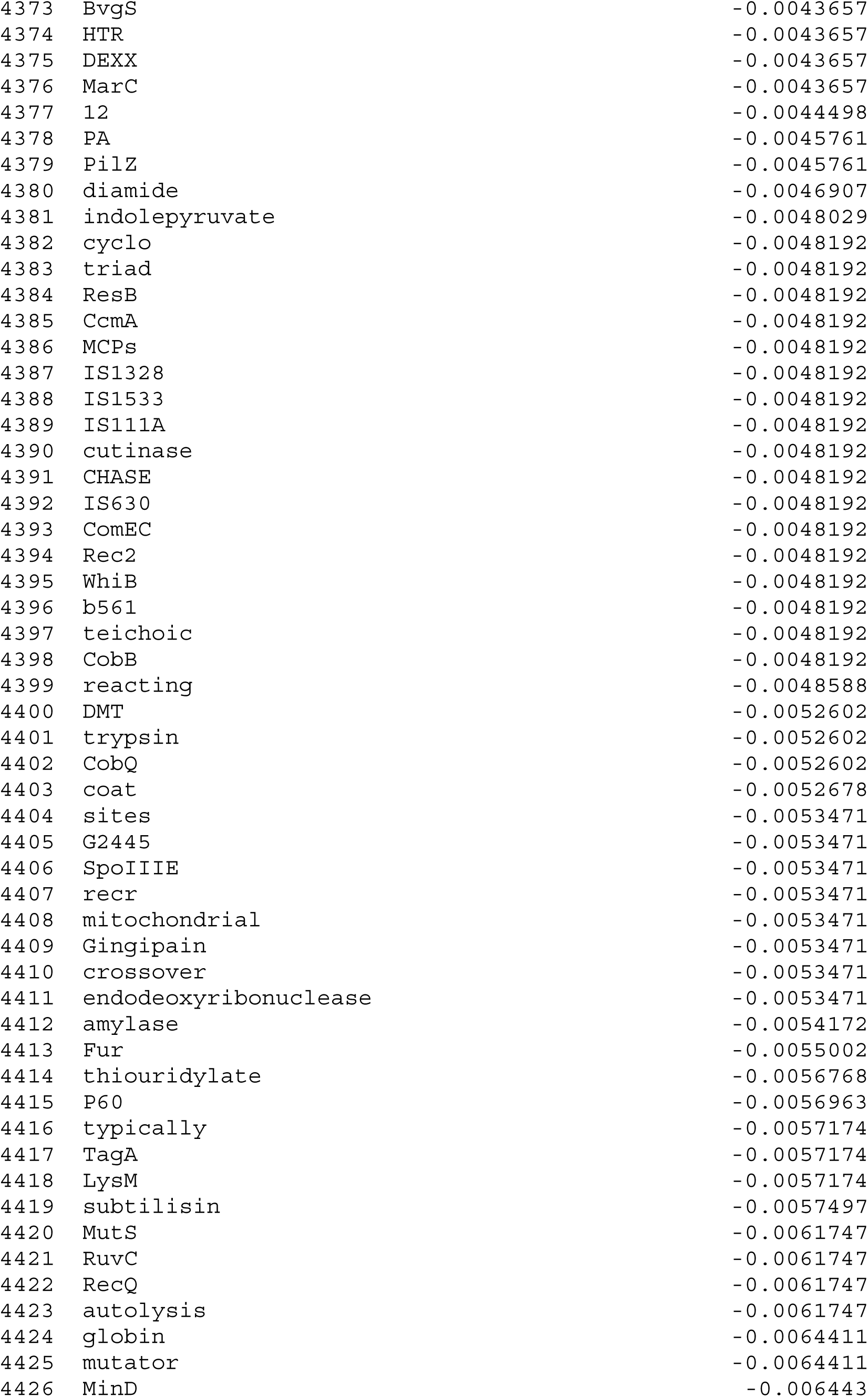

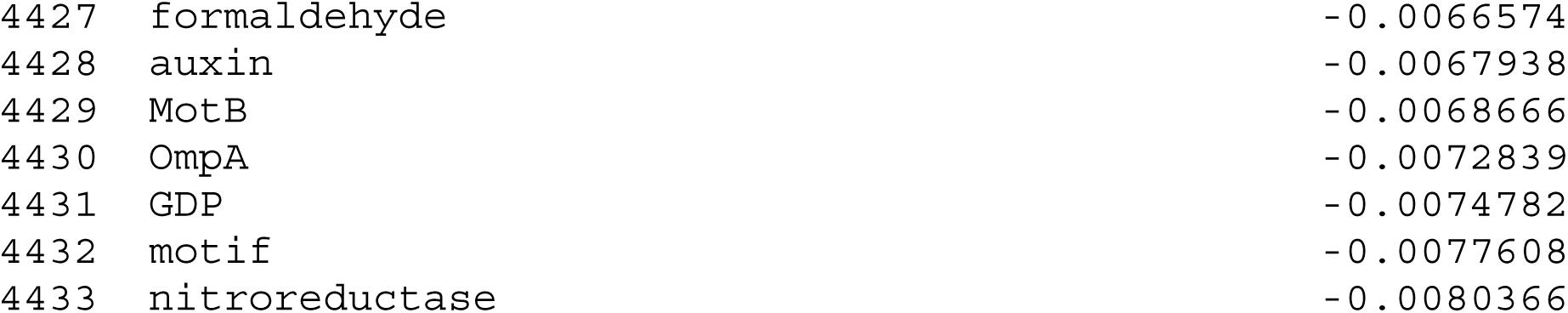

